# Regional heterogeneity in gene expression, regulation and coherence in hippocampus and dorsolateral prefrontal cortex across development and in schizophrenia

**DOI:** 10.1101/426213

**Authors:** L Collado-Torres, EE Burke, A Peterson, JH Shin, RE Straub, A Rajpurohit, SA Semick, WS Ulrich, Consortium BrainSeq, C Valencia, R Tao, A Deep-Soboslay, TM Hyde, JE Kleinman, DR Weinberger, AE Jaffe

## Abstract

Recent large-scale genomics efforts have better characterized the molecular correlates of schizophrenia in postmortem human neocortex, but not hippocampus which is a brain region prominently implicated in its pathogenesis. Here in the second phase of the BrainSeq Consortium (Phase II), we have generated RiboZero RNA-seq data for 900 samples across both the dorsolateral prefrontal cortex (DLPFC) and the hippocampus (HIPPO) for 551 individuals (286 affected by schizophrenia disorder: SCZD). We identify substantial regional differences in gene expression, in both pre- and post-natal life, and find widespread differences in how genes are regulated across development. By extending *quality* surrogate variable analysis (qSVA) to multiple brain regions, we identified 48 and 245 differentially expressed genes (DEG) by SCZD diagnosis (FDR<5%) in HIPPO and DLPFC, respectively, with surprisingly minimal overlap in DEG between the two brain regions. We further identified 205,618 brain region-dependent eQTLs (FDR<1%) and found that 124 GWAS risk loci contain eQTLs in at least one of the regions. We also identify potential molecular correlates of *in vivo* evidence of altered prefrontal-hippocampal functional coherence in schizophrenia. These results underscore the complexity and regional heterogeneity of the transcriptional correlates of schizophrenia, and suggest future schizophrenia therapeutics may need to target molecular pathologies localized to specific brain regions.

## Introduction

Schizophrenia is a psychiatric disorder that affects ~1% of the worldwide population and is linked to major socio-economic costs (Gore et al., 2011; Millan et al., 2016). As a highly heritable (Schizophrenia Working Group of the Psychiatric Genomics Consortium, 2014) disorder, it interferes with brain development and maturation, and has a wide range of severity and age of onset, though is typically diagnosed in young adults (Millan et al., 2016). Different treatments exist that have various degrees of efficacy against mainly so-called positive symptoms, though improving the treatment of these symptoms remains a crucial goal (Leucht et al., 2013; Millan et al., 2016).

Novel therapeutics used to treat psychiatric disorders have been lacking for many decades, primarily due to a lack of understanding of the underlying neural mechanisms that drive disease (Hyman, 2012). In disorders like schizophrenia, this problem is further exacerbated by the variety of complex and variable clinical characteristics. Genetic studies play a critical role in understanding the etiology of neurodevelopmental disorders and informing the development of novel therapeutics to treat these diseases. Recent efforts by the Psychiatric Genomics Consortium (PGC) have found hundreds of loci with a significant genome-wide association with risk for schizophrenia (Pardiñas et al., 2018; Schizophrenia Working Group of the Psychiatric Genomics Consortium, 2014). However, translating an association between a locus and a disease into therapeutic treatments is a challenging task: the biological importance of associated genetic variation must first be understood.

BrainSeq, a Human Brain Genomics Consortium consisting of six pharmaceutical companies working pre-competitively with the Lieber Institute for Brain Development, was initiated with the goal of generating publicly-available neurogenomic datasets (RNA sequence, genotype, and DNA methylation) in postmortem brain tissue to enhance the understanding of psychiatric disorders (BrainSeq Consortium, 2015). In the first phase of BrainSeq, we identified widespread genetic, developmental, and schizophrenia-associated changes in polyadenylated RNAs in the dorsolateral prefrontal cortex (DLPFC) (Jaffe et al., 2018) using polyA+ RNA sequencing (RNA-seq) to prioritize protein-coding changes in gene expression. Other research groups, in particular the Common Mind Consortium, have also focused on understanding gene expression and regulation in schizophrenic brain (Fromer et al., 2016). However, in general, the transcriptional landscape of hippocampus (HIPPO), another region prominently implicated in the pathogenesis of schizophrenia (Callicott et al., 1998; Rasetti et al., 2014; Weinberger, 1999), is much less well-explored, as current large consortia have prioritized neocortical brain regions like DLPFC (Fromer et al., 2016; PsychENCODE Consortium et al., 2015) despite differences in neuronal clonal organization (Xu et al., 2014) and the timing of their formation (Rice and Barone, 2000; Weinberger, 1999).

Here, in the second phase of the BrainSeq Consortium, we studied the expression differences between DLPFC and HIPPO by performing RNA-seq using RiboZero on 900 tissue samples across 551 individuals (286 with schizophrenia) in both DLPFC (n=453) and in HIPPO (n=447, **Figure S1, Figure S2 A**). We quantified expression of multiple feature summarizations of the Gencode v25 reference transcriptome, including genes, exons and splice junctions. These summarization methods are consistent with (Jaffe et al., 2018) and allow the exploration of both the annotated and unannotated transcriptome. We compared expression within and across brain regions, modeled age-related changes in controls using linear splines, integrated genetic data to perform expression quantitative trait loci (eQTL) analyses, and performed differential expression analyses controlling for observed and latent confounders. By extending the experiment-based quality surrogate variable (qSVA) framework (Jaffe et al., 2017), we adjusted for RNA degradation across multiple brain regions and reduced false positives. We also explored at the molecular level evidence from *in vivo* imaging studies that the pattern of functional coherence between DLPFC and HIPPO is altered in patients with schizophrenia. Together these results help shed light on the complex regional and molecular heterogeneity of schizophrenia-associated gene expression in human brain.

## Results

The BrainSeq Phase II dataset was created to better understand the differences in brain gene expression in developmental regulation, genetic control and subsequent dysregulation in schizophrenia (**Figure S1**, Material and Methods: postmortem brain tissue acquisition and processing, RNA sequencing). After processing the RNA-seq data, we performed extensive quality control to identify low quality samples and resolve potential sample swaps prior to filtering features with low expression across both brain regions together (**Figure S3**, **Figure S4**, Materials and Methods: RNA-seq processing pipeline, feature filtering by expression levels, genotype data processing, and the experiment-based qSVA approach to RNA quality control). We analyzed 24,652 genes, 396,583 exons, and 297,181 exon-exon junctions that were expressed across experiment-specific subsets of the 900 samples for all subsequent analyses.

### Regional differences in expression between brain regions across development

We first explored differences in expression between DLPFC and HIPPO among non-psychiatric controls (Materials and Methods: differential expression by brain region in prenatal and adult samples). We compared the regions using both adult (age>=18 years, range: 18 to 96 years) and prenatal (age<0, range: 14 to 22 post-conception weeks) samples, separately (**Figure S2 B**, total adult n=460, prenatal n=57). We also used 26 subjects (adult n=8, prenatal n=18) across 52 samples from the same two regions available in the smaller BrainSpan project as a potential replication dataset (BrainSpan, 2011) (**Figure S2 C**). Replication rate of differentially expressed features was determined across several Bonferroni-adjusted p-value thresholds while further requiring a consistent directionality of the expression differences (**Figure 1 A**,Materials and Methods: Replication analysis with BrainSpan). This conservative analysis identified 1,612 and 32 genes differentially expressed between DLPFC and HIPPO among adult and prenatal age groups, respectively, (**Figure 1 B**) at Bonferroni-adjusted p-value <0.01 and replicating in BrainSpan. These results echo earlier reports (Sousa et al., 2017) suggesting cortical gene expression during fetal life reflects less differentiation of cell types and connectivity patterns compared with adult cortex. Moreover, individual exons from a total of 3,375 genes and exon-exon junctions from 37 genes were differentially expressed between these two regions in adults (**Figure 1 B**). Transcript-level results were mostly complementary in adults as only one differentially expressed transcript was detected in the prenatal age group (**Figure S5**). These differential expression results in adults reflect fewer reads spanning exon-exon junctions and differences in complexity between gene and exon read assignments and coverage, particularly using ribosomal depletion. The modest differential expression results for the prenatal age group could be due at least in part to the relatively small sample sizes in both BrainSeq Phase II and BrainSpan.

**Figure 1.**
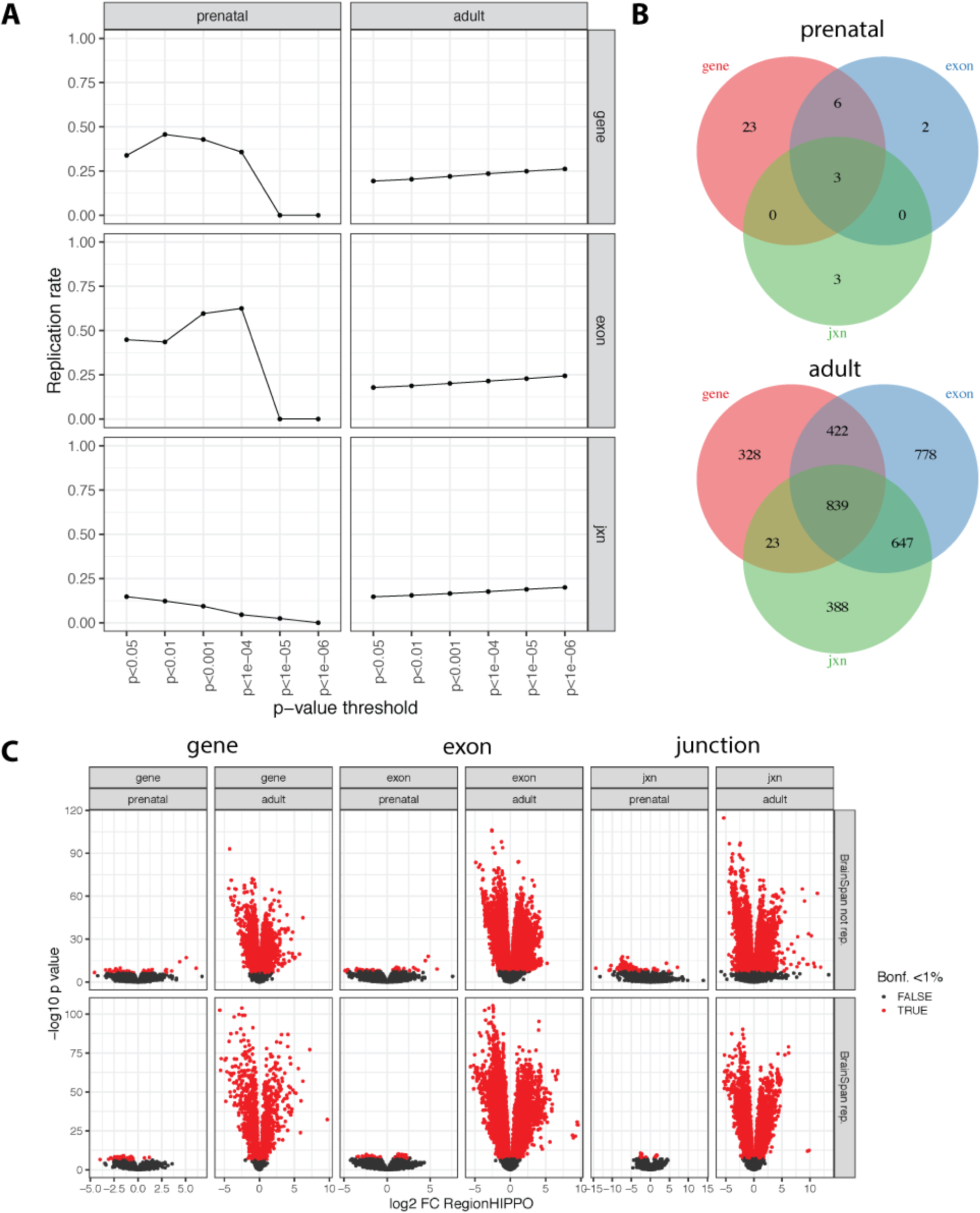
(**A**) Replication rate with consistent fold change for the adult and prenatal differences between brain regions using the BrainSpan dataset (n=16 for adult, n=36 for prenatal) across p-value thresholds for the exon, gene and exon-exon junction features. (**B**) Differentially expressed features grouped by gene ID for the adult and prenatal age groups. (**C**) Volcano plots of the differential expression signal between brain regions for each feature type and age group stratified by whether the differential expression replicated in BrainSpan (p<0.05 and consistent fold change).

Overall, there were large brain regional expression differences, particularly among adults by effect size as shown in **Figure 1 C**. However, we note that many of the strongest effects did not replicate in BrainSpan which could be due to the smaller sample size in that dataset. We did observe, however, higher replication rates when only considering directionality (**Figure S6**). In the adult samples, enriched gene ontology (GO) terms and Kyoto Encyclopedia of Genes and Genomes (KEGG) pathways for differentially expressed genes (DEGs) include biological processes such as axonogenesis and neurogenesis that are consistently observed across the three main expression features (**Figure S7 A**) with higher expression in DLPFC than HIPPO. We also observed DEG enrichment for ion channel activity and binding molecular functions (**Figure S7 B**), synaptic membrane and transporter complex cellular components (**Figure S7 C**)and GABAergic and cholinergic synapse pathways (**Figure S7 D**, exons and junctions only). In the prenatal age group, although overall there were fewer enriched GO terms and KEGG pathways (**Figure S8**) for the regionally DEGs, likely because there were fewer DEGs, increased positive regulation of neurogenesis (**Figure S8 A**) as well as relatively increased basal transcription machinery binding (**Figure S8 B**), nuclear chromatin (**Figure S8 C**), and biosynthesis of amino acids (**Figure S8 D**) were found in HIPPO compared with DLPFC. These findings are likely consistent with the earlier neuronal maturation of HIPPO compared with DLPFC (Rice and Barone, 2000). These analyses suggest expression landscapes in HIPPO and DLPFC begin to diverge in prenatal life and further diverge across brain development and aging.

### Developmental differences in expression between brain regions

We hypothesized that these differences in expression between brain regions could arise from differences in the cellular components of these regions. Using DNA méthylation data we estimated cell type proportions (Materials and Methods: DNA méthylation) for embryonic stem plus precursor cells (ESC+NPC), glial and neuronal cells and found that the differences in cell composition between DLPFC and HIPPO within the prenatal age group get larger with advancing months of gestation (**Figure 2 A**) which supports the results from the above differentially expressed genes between the two brain regions in comparing the prenatal and adult age groups (**Figure 1 B**). Given the changes in cell composition between the brain regions over development, we expected to find large expression differences across development.

**Figure 2.**
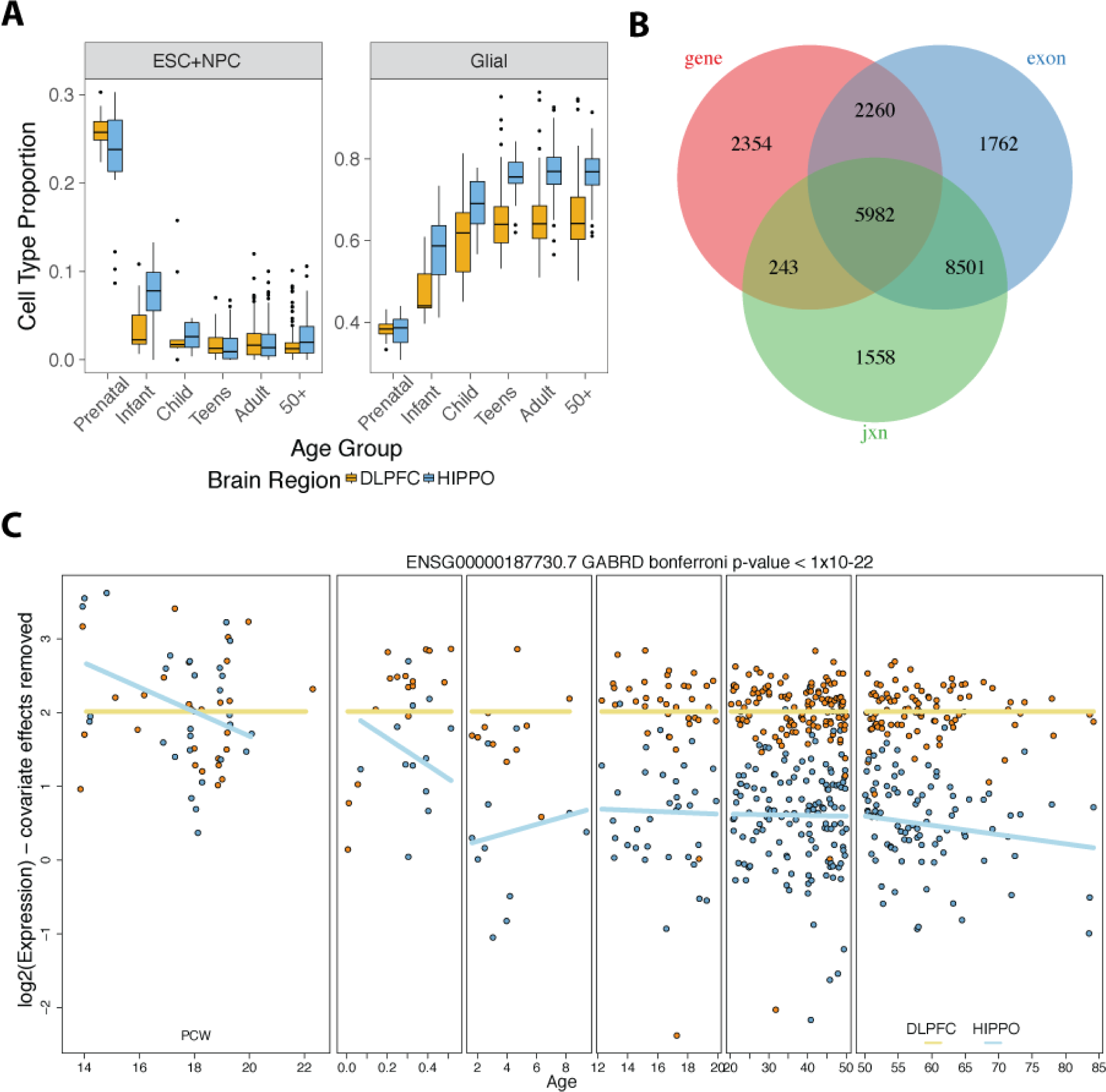
(**A**) Estimated cell type proportions for precursor cells (ESC+NPC) and glial cells across age groups using DNA methylation data. Neuronal cells are not shown. (**B**) Differentially expressed features between brain regions across development grouped by gene ID. (**C**) *GABRD* is differentially expressed between DLPFC and HIPPO across development. Y-axis shows the residualized expression after removing modeled covariates. PCW: post-conception weeks.

We therefore partitioned age into six age groups and used a linear spline model to identify changes in expression between DLPFC and HIPPO across development and aging among all non-psychiatric controls (**Figure S2 B**, DLPFC n=300, HIPPO n=314). We Bonferroni-adjusted the p-values and used BrainSpan (**Figure S2 C**, n=79) to survey the features that replicated in this external dataset (p<0.05). Exploration of the top 8 principal components (PCs) showed that brain region, pre- versus post-natal age (**Figure S9**, PC1 mean 20.5% and PC2 mean 10.9%, **Figure S9**) and sex (**Figure S10**) represented the main sources of variation across all feature types, while race was not associated with the top 8 PCs (**Figure S11**). The BrainSpan dataset presented similar patterns (age: **Figure S12**, sex: **Figure S13**, race: **Figure S14**) and we included these and other covariates as adjustment variables in the model (Materials and Methods: Differential expression over development).

We identified widespread differences in transcriptional regulation between the DLPFC and HIPPO over development, with 10,839 genes differentially expressed between these regions dependent on age period (Bonferroni < 0.01) that are nominally replicated in BrainSpan (p<0.05, **Figure 2 B**). Of these genes, 5,982 (55%) contain differentially expressed exons and splice junctions that replicated in BrainSpan (Materials and Methods: Replication analysis with BrainSpan). For example, *GABRD* (**Figure 2 C**), which encodes a subunit of the major inhibitory neurotransmitter receptor in the mammalian brain, shows decreased expression in HIPPO compared to DLPFC in several of the age groups. Enriched GO terms among differentially expressed genes dependent on development at the gene, exon or exon-exon junction levels include regulation of GTPase activity and dendrite development biological processes (**Figure S15 A**), GTPase regulator activity molecular function (**Figure S15 B**), synaptic membrane and postsynaptic density cellular components (**Figure S15 C**), and calcium signaling and RNA degradation KEGG pathways (**Figure S15 D**). These results suggest unique developmental profiles of expression in the HIPPO and DLPFC, and likely further associate with the shifting cellular landscapes of development in these brain regions.

### Unique schizophrenia-associated expression differences in HIPPO compared to DLPFC

Given the different expression levels and developmental regulation between DLPFC and HIPPO, we next asked whether different genes were associated with schizophrenia diagnosis in the two regions. As we have shown previously (Jaffe et al., 2017), RNA degradation is a major if not universal confounder in SCZD cases versus non-psychiatric controls comparisons, due in part to different ante- and post-mortem conditions that is not fully addressed by adjusting for observable quality metrics (e.g. RIN, mitochondrial mapping rate, pH, etc) (**Figure S16 A-B**). The effect of RNA degradation can be better adjusted for by using the *quality* surrogate variable analysis (qSVA) framework which is based on an *ex vivo* RNA degradation experiment (Jaffe et al., 2017). We modified the qSVA framework so that it could be applied to more than one brain region by identifying a common set of expressed regions (Collado-Torres et al., 2017) that are associated with degradation across both brain regions (**Figure S17**, Material and Methods: Degradation data generation, determining multi-region quality surrogate variables). Exploratory data analysis led us to exclude 43 HIPPO samples prepared with a different sequencing kit, as well as age <17 samples, given that there are no SCZD cases in that age range in the BrainSeq Phase II dataset (**Figure S16 C-F**, **Figure S2 D**). We identified 15 and 16 quality surrogate variables (qSVs) for DLPFC and HIPPO respectively, and 22 when using both regions together. The top qSV for each brain region is similarly associated with RIN and SCZD diagnosis (**Figure S18**) as well as with other technical covariates. Degradation quality plots for each brain region suggest that the adapted qSVA workflow is effective in reducing the confounding by RNA degradation for the SCZD case-control analysis (**Figure S19**, Material and Methods: DEqual plots). For example, a naïve model that does not adequately control for degradation, i.e. one based only on observed quality control metrics (e.g. RIN, pH, mito mapping rate), generated 6,429 differentially expressed by SCZD status in HIPPO at FDR<5%.

In contrast, when using qSVA, in HIPPO we identified 48 significantly differentially expressed genes between patients and controls (27 Control > SCZD, down; 21 Control < SCZD, up) while in DLPFC we identified 245 genes significantly differentially expressed (142 Control > SCZD, down; 103 Control < SCZD, up) at FDR< 5%. These numbers of genes scaled with the false discovery rate, as there were 101 genes (52 down, 49 up) in HIPPO and 632 in DLPFC (379 down, 253 up) at FDR<10%. We considered a more liberal set of DE genes in hippocampus (FDR<20%, N=332: 171 down, 161 up) than DLPFC (FDR<10%, N=632) for gene set enrichment analyses to ensure a sufficient number of input genes for inference. Interestingly, among the two sets of genes associated with differential expression based on diagnosis in the two regions, there was remarkably little overlap at the different feature levels (**Figure 3 A-B** for genes, **Figure S20** for other expression features) and also little overlap across expression features (**Figure S21 A-E**) when grouped by Gencode gene ID.

**Figure 3.**
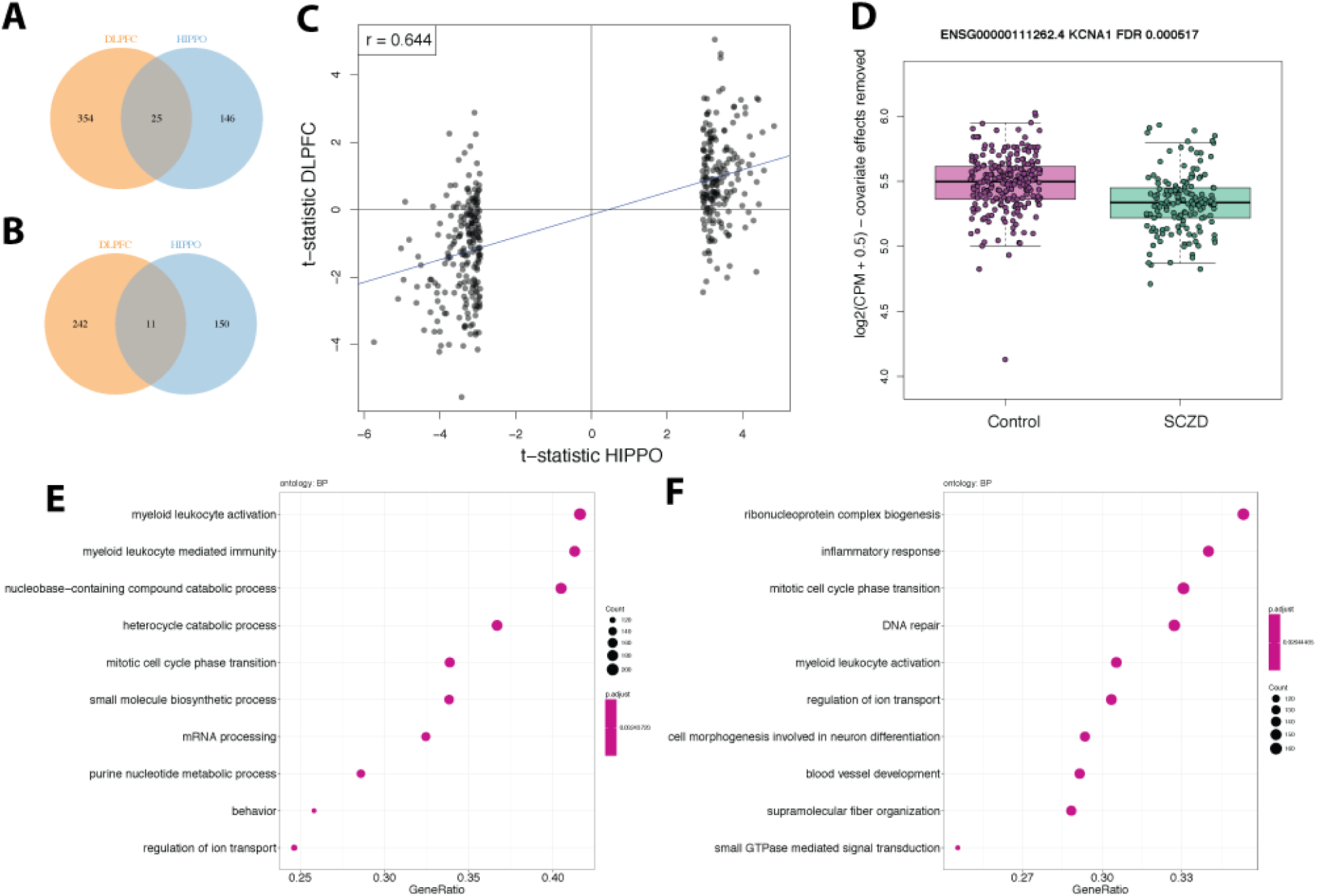
(**A**) Venn diagram of differentially expressed genes with higher expression in non-psychiatric controls than SCZD affected individuals by brain region (DLPFC: FDR <10%, HIPPO: FDR <20%). (**B**) Similar to (**A**) but for genes with higher expression in SCZD than in non-psychiatric controls. (**C**) T-statistics for the top 400 differentially expressed genes in HIPPO compared to their t-statistics in DLPFC. (**D**) *KCNA1* is differentially expressed in DLPFC. (**E**) Gene set enrichment (GSE) analysis on DLPFC with biological process ontology terms and HIPPO (**F**).

We used previously published data from BrainSeq Phase 1 (BSP1) and the CommonMind Consortium (CMC) to examine replication of these differences in DLPFC, which specifically could assess differences in library preparation protocols (polyA+ versus RiboZero) for a largely overlapping sample set and independently sequenced subjects in different labs, respectively. At the gene level, the DLPFC data significantly correlate (ρ=0.81) with the DLPFC BrainSeq Phase I (BSP1) results based on PolyA+ (**Figure S22**) as well as the CommonMind Consortium (CMC) dataset based on RiboZero (ρ=0.53, **Figure S22 B**) that we had previously re-processed (Fromer et al., 2016; Jaffe et al., 2018). In line with the lack of regional overlap among DE genes, the t-statistics for the top 400 differentially expressed genes by SCZD status in HIPPO do not predict the t-statistics in DLPFC (**Figure 3 C**), further showcasing the striking regional specificity. Similar comparisons against BSP1 (ρ=0.70) and CMC (ρ=0.18, **Figure S22**) show greater concordance with our DLPFC results than HIPPO. For example, among the genes identified in BSP1 at FDR<10% that replicated in CMC, *KCNA1* is differentially expressed in DLPFC but not in HIPPO (**Figure 3 D**) despite BSP1 and DLPFC having different RNA-seq library preparation protocols.

We performed gene set enrichment analyses at the gene level for DEGs based on diagnosis in each brain region, which did show more overlap in biological processes than individual DEGs had suggested. In both DLPFC and HIPPO we found that myeloid leukocyte activation and regulation of ion transport processes (**Figure 3 E-F**, **Figure S23 A**, **Figure S24** A), peptidase and ATPase activity molecular functions (**Figure S23 B**, **Figure S24 B**) were enriched. For cellular components, axon and endosome membrane were enriched in DLPFC (**Figure S23 C**) while nuclear chromosome and actin cytoskeleton were enriched for HIPPO (**Figure S24 C**). For KEGG, we found enrichment of Alzheimer’s and Parkinson’s disease pathways for DLPFC (**Figure S23 D**) and cytosolic DNA-sensing and influenza A pathways for HIPPO (**Figure S24 D**), while phagosome and tuberculosis pathways were enriched in both brain regions. We also performed GO enrichment analyses (**Figure S25**) and found primarily enrichment across genes with decreased expression among SCZD cases for both brain regions. We found several biological processes related to immune processes among differentially expressed genes in both regions (**Figure S25 A**), but no overlap among enriched molecular functions across brain regions and expressed features (**Figure S25 B**), with neuronal components enriched in DLPFC and phagocytic vesicle in HIPPO across expression features (**Figure S25 C**), and no overlap among brain regions in KEGG pathways with protein processing in endoplasmic reticulum appearing in HIPPO and tuberculosis and Ras signaling pathway in DLPFC (**Figure S25 D**). We identified 111 region-dependent DEGs using interaction modeling between SCZD diagnosis status and brain region (**Figure S26 A**, 424 with support at the gene, exon or junction expression level) such as *SCL02A1* (**Figure S26 B**) that are enriched for axonogenesis, axon development, postsynaptic density and neuron to neuron synapse biological processes and cellular components (**Figure S26 C-D**). These interaction DEGs that showed differential effects across the two brain regions had minimal overlap with either the SCZD DEGs in HIPPO (4/48) or DLPFC (5/245). These results in total suggest largely unique differentially expressed genes and processes associated with schizophrenia in the HIPPO compared with DLPFC and underscore the incompleteness of the molecular pathology associated with schizophrenia based on only DLPFC or HIPPO. The enrichment of immune processes in genes relatively decreased in expression in schizophrenia echoes several recent studies that have not confirmed earlier proposals of an upregulated immune response in the brain of patients with this illness (Birnbaum et al., 2018; Plavén-Sigray et al., 2018).

### Decreased regional coherence in schizophrenia via co-expression analysis

The clinical imaging research literature contains a number of reports of altered patterns of correlated measures of HIPPO and DLPFC in SCZD subjects compared to non-psychiatric controls (Meyer-Lindenberg et al., 2001; Weinberger et al., 1992). Evidence of altered coherence of gene expression across these regions in schizophrenia has not previously been explored. Using the subset of individuals with data in both brain regions (n=265 subjects, 530 RNA-seq samples), we first computed correlations across individuals for each gene to determine how genes were co-expressed across these brain regions. Those genes most consistently co-expressed (at FWER<5%) across these regions were enriched for GO terms related to the immune response, response to virus, cytokine activity (**Figure S27 A-C**) and cytokine-cytokine receptor interaction KEGG pathways (**Figure S27 D**), suggesting that immune processes are particularly coherent across these brain regions within individuals. Interestingly, there was no association between gene co-expression across brain regions and subsequent differential expression in schizophrenia, supporting the regional heterogeneity across brain region by diagnosis described above (**Figure S28**).

We next assessed decreased coherence at the subject-, rather than gene-, level across the entire transcriptome, which would be more comparable to identifying decreased connectivities via neuroimaging and other studies (Friston et al., 2016). We found that SCZD-affected individuals had significantly lower transcriptome-wide correlation across DLPFC and HIPPO for all expression feature types we assessed (**Figure S29**, gene-level p=0.0164). This difference was accentuated for the 48 SCZD DEGs in HIPPO (**Figure S30 A**, p= 1.89×10^−9^) and missing for the DLPFC, BSP1 and CMC DEGs (**Figure S30 B-D**). We further refined these analyses by computing more specific individual-level correlations for sets of genes defined by biological functions (grouped by GO term and KEGG pathways). We identified 10 cellular component, 14 biological processes and 3 molecular function GO terms as well as 13 KEGG pathways with significant correlation differences (at FDR<5%) between SCZD-affected individuals and non-psychiatric controls (**Table S3**). Consistent with the dysconnection hypothesis and prior evidence from neuroimaging studies (Friston et al., 2016; Meyer-Lindenberg et al., 2001; Weinberger et al., 1992), 35 out of the 40 (87.5%) of the enriched terms and pathways had decreased correlation in the SCZD affected individuals (p=4.53×10^−6^, χ_1^2^_=20.0, **Table S3**). We highlight gene sets involving the dendritic spine neck (**Figure S31 A**) and positive regulation of long term synaptic depression (**Figure S31 B**) having decreased coherence in SCZD affected individuals, which have been previously linked to schizophrenia (Crabtree and Gogos, 2014; Hasan et al., 2012; Penzes et al., 2011). We further identified that coherent activation of the immune response (**Figure S31 C**) across regions was also decreased in SCZD individuals, in agreement with our differential expression analyses among individual features. At the pathway level, the lysosome pathway (**Figure S31 D**) is among the pathways with decreased correlation which is in agreement with previous studies that identified the lysosome as one of the components with dysregulated function in schizophrenia (Zhao et al., 2015). These findings provide novel insight into the potential molecular mechanisms underlying decreased functional coherences between the hippocampus and frontal cortex in schizophrenia.

### Differences in genetic regulation of expression between brain regions

The altered connection hypothesis of schizophrenia is partially supported by the GWAS findings (Friston et al., 2016; Pardiñas et al., 2018; Schizophrenia Working Group of the Psychiatric Genomics Consortium, 2014) that have associated genetic risk with genes implicated in NMDAR function and plasticity at glutamatergic synapses, and also with genes showing association with altered connectivity measures on functional MRI (Rasetti et al., 2014). To further explore the impact of genetic risk, we combined genotype data with the RNA-seq data to assess the overlap in genetic regulation by brain region. We identified expression quantitative trait loci (eQTL) for each of the expression features and brain regions (Materials and Methods: eQTLs) using individuals with age > 13 (n=477). In HIPPO we found 11,237,357 eQTL associations (SNP-feature pairs, at FDR <1%) across genes, exons and junctions corresponding to 17,719 genes (**Figure 4 A**, **Figure S32 A**) and in DLPFC we found 15,766,398 eQTLs for 19,482 genes across the same spectrum of expression features. We then performed joint analyses in DLPFC and HIPPO to identify 205,618 (at FDR< 1%) brain region-dependent eQTLs (i.e. statistical interaction between genotype and region, **Figure 4 B**, **Figure S32 B**) corresponding to 1,484 genes (**Figure 4 C**, **Figure S32 C**) across all three expression feature types. These results show substantial regional specificity in genetic regulation of expression. The eQTL associations we identified showed low replication rates in GTEx v6 (GTEx Consortium, 2015) when considering directionality and p<0.05 but higher rates when only considering directionality (HIPPO: 31.9% vs 69.9%, DLPFC: 32.5% vs 70%, brain-region dependent: 24.6% vs 67.4% at the gene level), likely due to the much smaller sample sizes (GTEx: 92 and 82, BSP2: 397 and 395 for DLPFC and HIPPO respectively).

**Figure 4.**
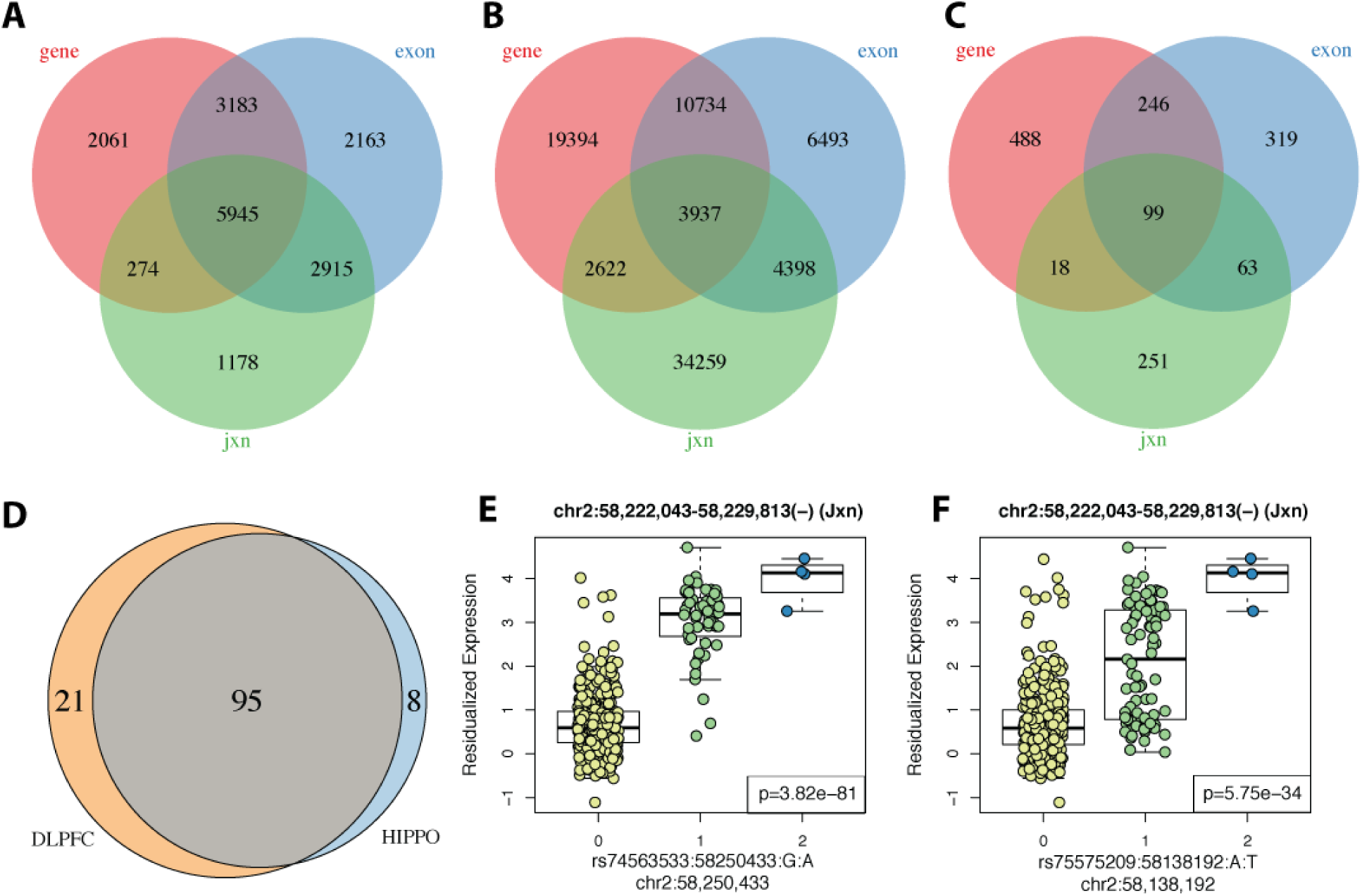
(**A**) HIPPO eQTLs grouped by gene ID. (**B**) region-dependent eQTLs grouped by SNP ID. (**C**) Region-dependent eQTLs group by gene ID. (**D**) Unique risk schizophrenia GWAS index SNPs from PCG2 (Pardiñas et al., 2018) by brain region that are in eQTLs. (**E**) Top HIPPO eQTL among schizophrenia risk SNPs. This exon-exon junction skips exon 4/14 of the *FANCL* gene. (**F**) Index schizophrenia risk SNP corresponding to the proxy SNP from (E).

We explored whether any of these eQTLs were previously-identified schizophrenia GWAS risk variants (Pardiñas et al., 2018). Among the HIPPO eQTLs, we found eQTL associations to 60 risk SNPs such as rs42945 and expression features from the *NDRG4* gene (**Figure S33 A-B**) that has been previously implicated in sudden cardiac death among users of antipsychotics in patients with schizophrenia (Watanabe et al., 2017). Similarly, 3 risk SNPs (rs4144797, rs12293670, and rs324015) were brain region-dependent eQTLs that involved features from genes *LRP1*, *NRGN* and *NGEF* (**Figure S34 A-D**) and have been previously been identified as methylation QTL SNPs in DLPFC (Jaffe et al., 2016). While risk SNPs rs12293670 and rs4144797 were specific to the DLPFC, with weak to no eQTL signal in HIPPO, rs324015 showed opposing eQTL directionality by brain region to features in *LRP1* (**Figure S34 A-D**).

To more fully characterize regional-specific eQTLs in the context of schizophrenia risk SNPs, we carried out a targeted eQTL analysis by using samples age >13 (**Figure S32 D**, n=477, Materials and Methods: Risk loci) and GWAS variants. Using rAggr (Barrett et al., 2005) we identified proxy SNPs that have a linkage disequilibrium (LD) r2 score greater than 0.8 to the 179 index GWAS (Pardiñas et al., 2018) risk SNPs and used 163 risk loci that were common (MAF > 5%) and well-imputed in our genotype data (**Figure S32 E**, Materials and Methods: risk loci). Of these 163 risk loci, 124 (76%) were either eQTLs (FDR<1%) in HIPPO or DLPFC with 95 variants (76.6% of the eQTLs) shared across both regions (**Figure 4 D**). This percentage of GWAS significant SNPs showing association with gene expression is more than 50% greater than previous reports, again illustrating the incompleteness of assumptions about eQTLs based on only limited surveys of brain regions. Overall, 5,510 and 6,780 out of the 9,692 proxy and index SNPs were eQTLs (FDR<1%) in HIPPO and DLPFC spanning 1,731 and 2,525 different expression features respectively (**Figure S32 F**). The top eQTL in HIPPO corresponds to an exon skipping event in the *FANCL* gene (**Figure 4 E**) and involves proxy SNP rs74563533. The corresponding index SNP rs75575209 is also an eQTL (**Figure 4 F**) with the same exon-exon junction (chr2:58,222,043–58,229,813 in hg38 coordinates). The eQTL involving this junction and proxy SNP rs74563533 was also the top result in DLPFC in BSP1 (Jaffe et al., 2018) (p=2.796e-85, chr2:58,449,178-58,456,948 in hg19 coordinates). *FANCL* has been previously linked to Fanconi anemia disease (Meetei et al., 2003) and is involved in the DNA repair pathway (Machida et al., 2006). From the 103 and 116 risk loci in these eQTLs, 38 (36.9%) and 37 (31.9%) of the loci pair to a single gene in HIPPO and DLPFC (**Figure S32 G**) showcasing that in both brain regions roughly two thirds of the risk loci (63.1 and 68.1% respectively) are ambiguous to resolve. Through our eQTL browser resource http://eatl.brainsea.org/ we have made all eQTLs sets available for further exploration: global HIPPO and DLPFC eQTLs, brain-region dependent eQTLs, and schizophrenia risk HIPPO and DLPFC eQTLs.

## Discussion

In this second phase of the BrainSeq consortium project, we generated and processed 900 RNA-seq samples from human postmortem brains from 551 individuals of which 286 (52%) were affected with schizophrenia (**Figure S1**). We used the RiboZero library preparation method to preserve non-coding RNA, and examined two brain regions - the hippocampus (HIPPO) and dorsolateral prefrontal cortex (DLPFC) - thus expanding our previous analysis of 495 (175 with schizophrenia) DLPFC samples analyzed using polyA+ RNA-seq (Jaffe et al., 2018). This second phase expands the variety of analyses that can be carried out to further investigate differences among the two of the brain regions most consistently implicated in schizophrenia as well as their differences at given time points across development (**Figure S1**). As part of BrainSeq Phase II, we also generated genotype information and identified expression quantitative trait loci (eQTLs), thereby linking genetic risk variants for schizophrenia to biological consequences. By analyzing the expression data with multiple complementary annotation methods (gene, exon, exon-exon junction, transcript), we gleaned a more complete understanding of differences in the transcriptome (annotated and unannotated) (**Figure S1**).

Perhaps not surprisingly, we found that the expression in HIPPO and DLPFC is more similar in prenatal age than in adult age, with widespread differences across development and cell type composition as estimated using DNA methylation among non-psychiatric controls. These results are important for further understanding if expression differences associated with a given disorder vary by brain region or underlying cell types. Synaptic membrane, GTPase activity and regulation, calcium signaling and RNA degradation are among the cellular components, molecular functions and pathways that are different between the two brain regions. This information might be important when developing drugs, most of which will likely affect both the DLPFC and HIPPO.

Given that SCZD cases generally have a longer postmortem interval and lower pH than non-psychiatric controls (**Figure S16 A-B**) and that RNA degradation is a major and virtually universal confounder in SCZD case-control analyses, we adapted the experiment-based quality surrogate variable (qSVA) framework (Jaffe et al., 2017) for more than one brain region (**Figure S17**). We generated 40 RiboZero RNA-seq samples in toto across both DLPFC and HIPPO to identify degradation-associated expressed regions. We then quantified those regions in BrainSeq Phase II and used them to identify quality surrogate variables (qSVs) for each brain region. Using the qSVs for each brain region we were able to substantially reduce the effect of RNA degradation in our SCZD case-control analyses (**Figure S19**). By expanding the qSVA framework for two regions we have laid the groundwork for analyzing multiple brain regions with this framework, which involves carrying out degradation experiments for each new brain region.

We identified 48 and 245 differentially expressed genes among SCZD cases and non-psychiatric controls in HIPPO and DLPFC (FDR<5%), respectively, with little overlap among these genes and at higher FDR thresholds or other expressed features. This finding, in itself, highlights the molecular heterogeneity of this illness across these regions. We determined that genes that are significantly co-expressed between HIPPO and DLPFC (FWER<5%) were enriched for immune processes but showed no relation with the SCZD case versus non-psychiatric control differential expression signal. Interestingly, at the individual subject-level, we found that HIPPO and DLPFC have significantly (p<0.05) lower transcriptome-wide correlation at the gene, exon, exon-exon and transcript levels (**Figure S29**) in SCZD cases compared to non-psychiatric controls, which may echo and could provide molecular insights into *in vivo* evidence of altered connectivity between these regions (Friston et al., 2016; Meyer-Lindenberg et al., 2001; Weinberger et al., 1992). To our knowledge, this is the first evidence that gene expression is less coherent between these regions in schizophrenia. Among the GO terms with significant correlation differences we identified dendritic spine neck and positive regulation of long term synaptic depression (**Figure S31 A-C**), which had significant decreased correlation in SCZD cases and have been previously linked to schizophrenia (Crabtree and Gogos, 2014; Hasan et al., 2012; Penzes et al., 2011). We further found more supporting evidence linking the lysosome pathway (**Figure S31 D**) to SCZD, thus also complementing previous findings (Zhao et al., 2015).

We lastly characterized extensive genetic regulation of gene expression with significantly different effects across the two brain regions: we identified 205,618 brain region-dependent eQTLs (at FDR < 1%), corresponding to 1,484 genes, at the gene, exon, or exon-exon junction level. These brain region-dependent eQTLs included clinically relevant risk variants as five schizophrenia risk loci showed significant differential regional regulation. We also identified millions of eQTLs in both HIPPO and DLPFC (FDR<1%) across genes, exons and exon-exon junctions. A sub-analysis focusing on schizophrenia GWAS risk loci showed that 124 of the 163 risk loci observed in our dataset are eQTLs with 95 (76% of the eQTLs) being shared across HIPPO and DLPFC. Furthermore, we found that 6,970 risk SNPs (GWAS index and their proxies) are an eQTL in either brain region. In each brain region, over 60% of the risk loci are associated with more than one gene, highlighting the complexity of SNP association for this disease and the potential difficulty in interpreting the eQTL results. Given the complexity and richness of these eQTL findings, we have made all five sets of eQTL results (global and risk-focused for each brain region, plus regional-dependent ones) publicly available via the user friendly LIBD eQTL browser at http://eatl.brainsea.org/ to facilitate independent analyses.

Overall, we showed extensive regional specificity of developmental and genetic regulation, and schizophrenia-associated expression differences between the hippocampus and dorsolateral prefrontal cortex and their molecular correlation. These findings suggest that some components of the transcriptional correlates of development and genetic risk for schizophrenia are brain region-specific. Thus, it may be necessary to incorporate regionally targeted therapies for schizophrenia. These data and analysis methods, generated as part of the BrainSeq Phase II project, strengthen the foundation of our understanding of schizophrenia and enhance our ability to find new ways to improve the lives of individuals affected by this disease.

## Materials and Methods

### Postmortem brain tissue acquisition and processing

As previously described in Jaffe *et al.* 2016 (Jaffe et al., 2016), postmortem human brain tissue was obtained by autopsy primarily from the Offices of the Chief Medical Examiner of the District of Columbia, and of the Commonwealth of Virginia, Northern District, all with informed consent from the legal next of kin (protocol 90-M-0142 approved by the NIMH/NIH Institutional Review Board). Additional post-mortem prenatal, infant, child and adolescent brain tissue samples were provided by the National Institute of Child Health and Human Development Brain and Tissue Bank for Developmental Disorders (http://www.BTBank.org) under contracts N01-HD-4-3368 and N01-HD-4-3383. Postmortem human brain tissue was also provided by donation with informed consent of next of kin from the Office of the Chief Medical Examiner for the State of Maryland (under Protocol No. 12-24 from the State of Maryland Department of Health and Mental Hygiene) and from the Office of the Medical Examiner, Department of Pathology, Homer Stryker, M.D. School of Medicine (under Protocol No. 20111080 from the Western Institute Review Board). The Institutional Review Board of the University of Maryland at Baltimore and the State of Maryland approved the protocol, and the tissue was donated to the Lieber Institute for Brain Development under the terms of a Material Transfer Agreement. Clinical characterization, diagnoses, and macro- and microscopic neuropathological examinations were performed on all samples using a standardized paradigm, and subjects with evidence of macro- or microscopic neuropathology were excluded, as were all subjects with any psychiatric diagnoses. Details of tissue acquisition, handling, processing, dissection, clinical characterization, diagnoses, neuropathological examinations, and quality control measures were further described previously (Lipska et al., 2006). Postmortem tissue homogenates of the prefrontal cortex (dorsolateral prefrontal cortex, DLPFC, BA46/9) were obtained from all subjects.

### RNA sequencing

Total RNA was extracted from samples using the RNeasy Plus Mini Kit (Qiagen). Paired-end strand-specific sequencing libraries were prepared from 300ng total RNA using the TruSeq Stranded Total RNA Library Preparation kit with Ribo-Zero Gold ribosomal RNA depletion (Illumina). An equivalent amount of synthetic External RNA Controls Consortium (ERCC) RNA Mix 1 (Thermo Fisher Scientific) was spiked into each sample for quality control purposes. The libraries were sequenced on an Illumina HiSeq 3000 at the LIBD Sequencing Facility, after which the Illumina Real Time Analysis (RTA) module was used to perform image analysis and base calling and the BCL converter (CASAVA v1.8.2) was used to generate sequence reads, producing a mean of 125.2 million 100-bp paired-end reads per sample.

### RNA-seg processing pipeline

Raw sequencing reads were quality checked with FastQC (Babraham Bioinformatics, 2016), and where needed leading bases were trimmed from the reads using Trimmomatic (Bolger et al., 2014) as appropriate. Quality checked reads were mapped to the hg38/GRCh38 human reference genome with splice-aware aligner HISAT2 version 2.0.4 (Kim et al., 2015). Feature-level quantification based on GENCODE release 25 (GRCh38.p7) annotation was run on aligned reads using featureCounts (subread version 1.5.0-p3) (Liao et al., 2014) with a mean 45.3% (SD=7.4%) of mapped reads assigned to genes. Exon-exon junction counts were extracted from the BAM files using regtools (McDonnell Genome Institute) v. 0.1.0 and the `bed_to_j uncs` program from TopHat2 (Kim et al., 2013) to retain the number of supporting reads (in addition to returning the coordinates of the spliced sequence, rather than the maximum fragment range) as described in (Jaffe et al., 2018). Annotated transcripts were quantified with Salmon version 0.7.2 (Patro et al., 2017) and the synthetic ERCC transcripts were quantified with Kallisto version 0.43.0 (Bray et al., 2016). For an additional QC check of sample labeling, variant calling on 740 common missense SNVs was performed on each sample using bcftools version 1.2. We generated strand-specific base-pair coverage BigWig files for each sample using `bam2wig. py` version 2.6.4 from RSeQC (Wang et al., 2012) and `wigToBigwig` version 4 from UCSC tools (Kent et al., 2010). **Table S1** includes demographics for different subsets of samples and differences (if any) among technical covariates.

### Genotype data processing

Genotype data was processed and imputed as previously described (Jaffe et al., 2018). Briefly, genotype imputation was performed on high-quality observed genotypes (removing low quality and rare variants) using the prephasing/imputation stepwise approach implemented in IMPUTE2 (Howie et al., 2009) and Shape-IT (Delaneau et al., 2008), with the imputation reference set from the full 1000 Human Genomes Project Phase 3 dataset (1000 Genomes Project Consortium et al., 2015), separately by Illumina platform using genome build hg19. We retained common variants (MAF > 5%) that were present in the majority of samples (missingness < 10%) that were in Hardy Weinberg equilibrium (at p>1×10^−6^) using the Plink tool kit version 1.90b3a (Purcell et al., 2007). Multidimensional scaling (MDS) was performed on the autosomal LD-independent construct genomic ancestry components on each sample, which can be interpreted as quantitative levels of ethnicity – the first component separated the Caucasian and African American samples. This processing and quality control steps resulted in 7,023,860 common variants in this dataset of 551 unique subjects. We remapped variants to hg38 first using the dbSNP database (Sherry et al., 2001) (from v142 on hg19 to v149 on hg38) and then the liftOver tool (Hinrichs et al., 2006) for unmapped variants (that were dropped in dbSNP v149). eQTL analyses were performed using the 7,023,286 variants with hg38 coordinates (574 variants did not liftOver).

### Quality control

After completing the preprocessing pipeline, samples were checked for quality control measures. All samples had expected ERCC concentrations. Forty-two samples with poor alignment rates (<70%), gene assignment rates (<20%), and mitochondrial mapping rates (>6%, DLPFC only) were dropped. Next, for each sample we correlated the genotypes of missense single nucleotide variants (SNVs) found in the processing pipeline to those same SNVs based on genotyping described above, and dropped ten samples based on possible incorrect sample labeling, where the genotypes were lowly correlated to the same sample number, or highly correlated do a different sample number. Ultimately 900 samples passed quality control checks (HIPPO n=447, DLPFC n=453).

### Feature filtering bv expression levels

We filtered lowly expressed features using the `expression_cutof f ( )` function from the jaffelab (Collado-Torres and Jaffe, 2017) package v0.99.18 resulting in cutoffs based on the mean expression across all 900 samples: RPKM 0.25 for genes, RPKM 0.30 for exons, RP10M 0.46 for exon-exon junctions and 0.32 TPM for transcripts (**Figure S3**). 24,652 genes (42.5%), 396,583 exons (69.4%), 297,181 exon-exon junctions (35.5%) and 92,732 transcripts (46.8%) passed the expression filters. **Figure S4** shows the number of expressed features passing the expression cutoffs grouped by gene ids.

### Differential expression bv brain region in prenatal and adult samples

Using the neuropsychiatric control samples (**Figure S2 B**, **Table S1**) for the adult age group (age >= 18, HIPPO n=238, DLPFC n=222) or the prenatal group (age < 0, HIPPO and DLPFC n=28) we identified differentially expressed features by brain region while adjusting for age, technical covariates (mitochondrial mapping rate, total assigned gene rate, RIN) and ethnicity (first five principal components based on the genotype data) as shown in **Equation 1**. We used the voom method (Law et al., 2014) for genes, exons and exon-exon junctions and calculated the t-statistics for β_1_ = 0 using limma (Ritchie et al., 2015) v3.34.5 while adjusting for repeated measures using `dupiicateCorreiation ( )`. Resulting p-values were Bonferroni-adjusted within each feature type.

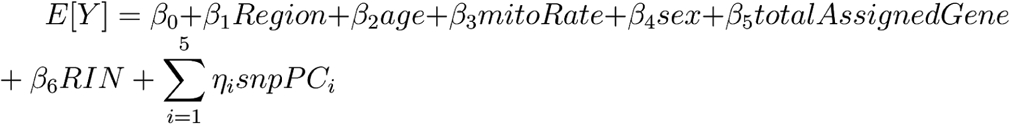

**Equation 1**. Full model for the differential expression analysis by brain region subsetted to specific age groups.

The full list and statistics for the differentially expressed features are available in **Table S2**.

### DNA methvlation

DNA methylation was assessed using the Illumina HumanMethylation450 ( “450k“) microarray as previously described (Jaffe et al., 2016). Samples from both brain regions were jointly normalized using stratified quantile normalization in the minfi Bioconductor package (Aryee et al., 2014). We retained a single array in the case of duplicates by choosing the sample that had the closest quality profile (via Methylated and Unmethylated signal intensity) to all other arrays. We implemented *in silico* estimation of the relative proportions of five cell types (ESCs, ES-derived NPCs, and derived dopamine neurons from culture, and adult cortex neuronal and non-neuronal cells from adult tissue) using the reference profile described in Jaffe et al, 2016 (Jaffe et al., 2016) with the deconvolution algorithm described by Houseman et al (Houseman et al., 2012).

### Differential expression over development

Using the neuropsychiatric control samples (**Figure S2 B**, **Table S1**, n=614) we fit a linear spline model with break points at birth/0, 1,10, 20, and 50 years of age while adjusting for the brain region (DLPFC was set as the reference, n=300, HIPPO n=314), sex, technical covariates (mitochondrial mapping rate, total assigned gene proportion and RIN) and ethnicity (first five principal components based on the genotype data). We computed F-statistics based on the null (**Equation 2**) and full (**Equation 3**) models shown below where we tested for an interaction between the age splines and the region indicator. For genes, exons and exon-exon junctions we used the voom method (Law et al., 2014) for normalizing the expression. To take into account the correlation induced by measuring two brain regions from the same individuals (in most cases) we used the `dupiicateCorreiation ( )` function in limma where the `block` and `design` arguments were set to the individual identifier and the intercept and region indicator, respectively. The resulting estimated correlation was passed to the `lmFit ( )` function in limma (Ritchie et al., 2015) version 3.34.5. Statistics were then calculated using `eBayes ( )` and `topTabie ( )` with the coef argument set to the additional terms in the full model (**Equation 3**). Resulting p-values were Bonferroni-adjusted within each feature type.

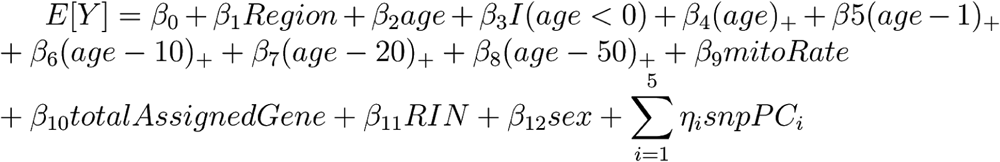

**Equation 2**. Null model for differential expression over development where *x*_+_ = *max*(0, *x*).

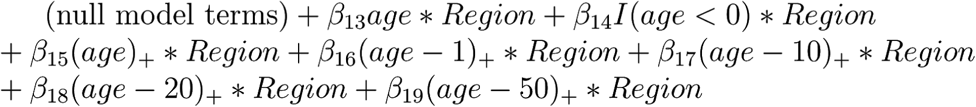

**Equation 3**. Additional terms present in the full model absent from the null model (**Equation 2**) that are tested in the F-statistic.

The full list and statistics for the differentially expressed features are available in **Table S2**

### Replication analysis with BrainSpan

To assess replication, we downloaded the fastq files from BrainSpan (BrainSpan, 2011) and processed the samples using the same RNA-seq processing pipeline and software versions. We retained only the samples with region code DFC and HIP and subsetted the measured features to those retained in our expression filtering cutoffs for BrainSeq Phase II (**Figure S2 C**). We then used the same models and procedure as for **Equation 1** for the differential expression by brain region analysis, and **Equation 2** and **Equation 3** for determining differential expression over development: p-value in BrainSpan had to be < 0.05 to be considered as replicating. For the differential expression by brain region analysis we required the log fold change to be consistent by directionality.

### Degradation data generation

The qSVA algorithm begins with measuring brain tissue degradation in a separate RNA-seq dataset to determine regions most-associated with degradation as illustrated in **Figure S17**. In this study, degradation was tested at four different time points for tissue from two brain regions, HIPPO and DLPFC, from five individuals (Jaffe et al., 2017). An aliquot of ~100 mg of pulverized tissue for each brain region from each donor was left on dry ice, and placed at room temperature until reaching the respective time interval, at which point the tissue was placed back onto dry ice (Jaffe et al., 2017). The four time intervals tested were 0, 15, 30, and 60 minutes, with the 0-minute aliquot remaining on dry ice for the entirety of the experiment, and RNA extraction began immediately after the end of the final time interval (Jaffe et al., 2017).

### Determining multi-reaion quality surrogate variables (qSVs’)

Raw fastq files were processed with the same RNA-seq processing pipeline as the BrainSeq Phase II data using the same software versions. The base-pair coverage RNA-seq expression data was normalized to 40 million reads using recount.bwtool (Ellis et al., 2018) version 0.99.28, and a cutoff of 5 reads was used to identify the expressed regions (ERs) of interest using derfinder version 1.12.6 (Collado-Torres et al., 2017). Transcript features most susceptible to RNA degradation were then identified using a regression model for degradation time adjusting for the brain region and individuals as shown in **Equation 4**. Individuals were adjusted for to account for variability due to subject effect, since brain tissue degradation was measured in the same brain at 4 different time points. The top 1000 ERs most associated with degradation (by FDR) were then quantified in the BrainSeq Phase II data using recount.bwtool.

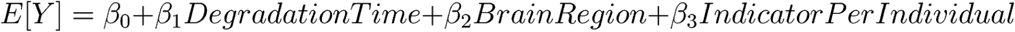

**Equation 4**. Linear model for identifying degradation-associated expressed regions.

We also used a model (shown in **Equation 5**) that included an interaction term between degradation time and brain region and found 620 ERs with a significant interaction term (FDR<5%) of which 603 had a significant degradation time term in the initial model. The genomic coordinates of the degradation-associated ERs used are available in **Table S4**.

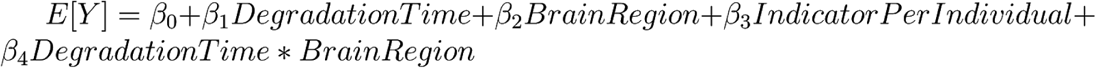

**Equation 5**. Same as **Equation 4** but also including an interaction term between degradation time and brain region.

From our exploratory data analysis of all 900 BrainSeq Phase II samples (**Figure S16**) we determined that it was best to drop the HIPPO Gold samples and samples age < 17 due to the lack of early SCZD cases (**Table S1**) before determining the actual qSVs (right side of **Figure S17**). We found three sets of qSVs with the BE algorithm (Buja and Eyuboglu, 1992) using sva (Leek and Storey, 2007) version 3.26.0: one for DLPFC samples (k=15, n=379), one for HIPPO samples (k=16, n=333) and one for both regions combined (k=22, n=712). The DLPFC and HIPPO top qSVs have similar relationships to RIN and SCZD diagnosis status as shown in **Figure S18**.

### DEaual plots

To assess the performance of the qSVA approach we used three different linear regression models for each brain region: a naïve model with only the SCZD diagnosis term (**Equation 6**), a model with all covariates we adjust for excluding qSVs (**Equation 7**), and a full model with all adjustment covariates and qSVs (**Equation 8**). We then compared the log2 fold change by SCZD diagnosis against the log2 fold change by degradation time for each brain region as shown in **Figure S19**.

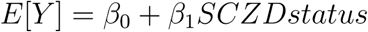

**Equation 6**. Linear regression model for the case-control analysis with a naive model.

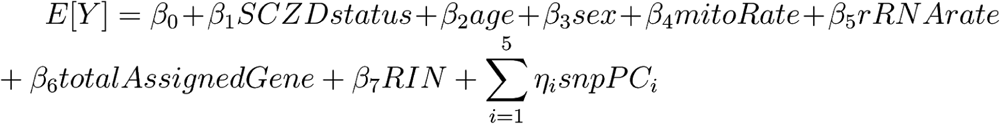

**Equation 7**. Linear regression model for the case-control analysis using all adjustment covariates except the quality surrogate variables (qSVs).

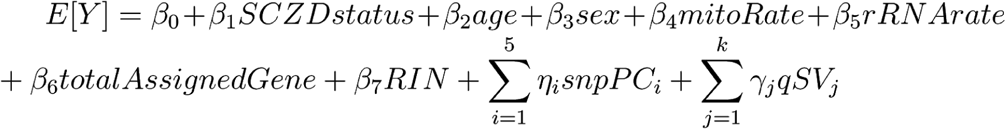

**Equation 8**. Linear regression model for the case-control analysis where *k* was determined by the BE algorithm (Buja and Eyuboglu, 1992) using sva (Leek and Storey, 2007) version 3.26.0 for each brain region separately (**Table S1**; DLPFC k=15, n=379; HIPPO k=16, n=333).

### Differential expression between SCZD cases and non-psvchiatric controls

Using voom (Law et al., 2014) from limma (Ritchie et al., 2015) version 3.34.9 we identified differentially expressed features using the model described in **Equation 8** for each brain region for genes, exons and exon-exon junctions. For transcripts we skipped the voom step. For comparisons with BrainSeq Phase 1 (Jaffe et al., 2018) and the CommonMind Consortium dataset (Fromer et al., 2016) we used the log fold changes we determined previously (Jaffe et al., 2018) and matched the sets by gene ids. The full list and statistics for the differentially expressed features are available in **Table S2**.

To identify differentially expressed features by the interaction between SCZD diagnosis status and brain region, we expanded **Equation 8** to include the interaction term and used the same methodology as previously described. The qSVs used in this analysis were determined acrossed both the DLPFC and HIPPO degradation matrices (N=712) resulting in k=22 qSVs.

### Gene ontology and gene set enrichment analyses

Unless otherwise noted, we used the compareCluster() function from clusterProfiler (Yu et al., 2012) version 3.6.0 for gene ontology (Ashburner et al., 2000; The Gene Ontology Consortium, 2017) and KEGG (Kanehisa et al., 2017) enrichment analyses with parameters `pvaiuecutof f=o. l` and `qvaiuecutof f=o. 05` with the set of Ensembl gene ids expressed in genes, exons and exon-exon junctions as the background universe (**Figure S4**). For gene set enrichment analysis in the SCZD case-control model, we used the `gseGO ()` function from clusterProfiler with default parameters using Ensembl gene ids.

### Visualization of differential expression results

We used the `cieaningY ()` function from the jaffelab package (Collado-Torres and Jaffe, 2017) version 0.99.20 to regress out adjustment covariates from our main differential expression models (**Equation 1**, **Equation 3**, **Equation 8**) to visualize the expression as shown in **Figure 2 C** and **Figure 3 D**. Doing so helps visualize what the model is observing.

### Venn diagrams

Colored venn diagrams were made using the VennDiagram R package version 1.6.18 while non-colored venn diagrams were made using gplots version 3.0.1.

### Brain region co-expression analyses

Using the age >17 RNA-seq samples from the SCZD case-control analysis, we identified the individuals with measured expression in both DLPFC and HIPPO (n=265 subjects, 530 RNA-seq samples). We computed the log2(RPKM + 0.5) for gene and exon expression levels, log2(RP10M + 0.5) for exon-exon junctions and log(TPM + 0.5) for transcripts. For DLPFC and HIPPO separately, we then used the `cieaningY ()` function from the jaffelab package (Collado-Torres and Jaffe, 2017) version 0.99.21 to remove all the covariates from **Equation 8** while keeping the intercept and SCZD diagnosis effects.

At the gene level, we computed the Pearson correlation for each gene across all 265 individuals producing one correlation value per gene. We then permuted the individual labels 1000 times to compute a null distribution of the gene correlation values and calculated FWER p-values using `derfinder: : : . caicPval ()` function (Collado-Torres et al., 2017). With the genes that had a significant correlation among brain regions (FWER<5%) we identified enriched GO terms and KEGG pathways using the same parameters previously described. We did this process for both the raw expression (say log2(RPKM + 0.5) at the gene level) and the “cleaned “ expression (post use of the `cieaningY ()` function).

We computed correlations at the individual level for all 4 expression summarizations (gene, exon, exon-exon junction, transcript) across all features, resulting in one correlation value per individual and expression level. We performed two-sided t-tests comparing the mean correlation among SCZD cases and non-psychiatric controls for each expression level. We grouped all the genes expressed in our dataset by their GO level 3 category for the biological process, cellular component and molecular function ontologies as well as by their KEGG pathway id. For each of these 4 gene sets, we calculated the individual level correlation between DLPFC and HIPPO, their two-sides t-test p-value for a difference in mean correlation, and corrected the p-values using the FDR method. GO terms and KEGG pathways with a significant difference are listed in **Table S3**.

We further computed correlations across brain regions using the sets of genes with significant differential expression by SCZD status identified in our previous analysis for both brain regions as well as in either the BSP1 and CMC datasets.

### Identifying eQTLs

MatrixEQTL (Shabalin, 2012) version 2.2 was used to identify eQTLs for each brain region using samples age > 13 (**Figure S32 D**, DLPFC n=397, HIPPO n=395) with either the log2(RPKM+1) for genes and exons, log2(RP10M+1) for exon-exon junctions or log2(TPM+1) for transcripts. For the eQTL analysis we adjusted for SCZD diagnosis status, sex, SNP PCs as well as expression PCs as shown in **Equation 9**. To determine the number of expression PCs to adjust (*p*) for we used the `num. sv ()` function from the sva package with the model in **Equation 10** and the `vfiiter=50000` parameter. Then the `Matrix eQTL main ()` was used to identify the eQTLs using the parameters `pvOutputThreshold. cis=0.001`, `pvOutputThreshold=0`, `useModel=modelLINEAR, cisDist=5e5.`

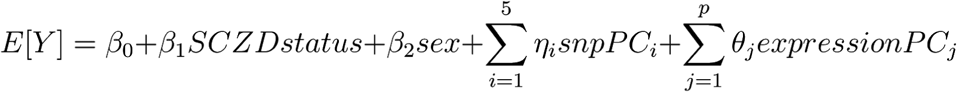

**Equation 9**. Main eQTL model for DLPFC and HIPPO.

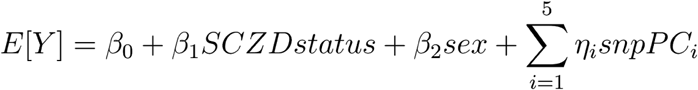

**Equation 10**. Model used for determining the number of expression PCs to adjust for in the eQTL analysis.

To identify eQTLs that have different effects by brain region, we ran a similar analysis using all the age > 13 samples (n=792) using the model from **Equation 11** and by changing some parameters Of `Matrix_eQTL_main ()` to `useModel=modelLINEAR_CROSS`, `cisDist=2.5e5`.

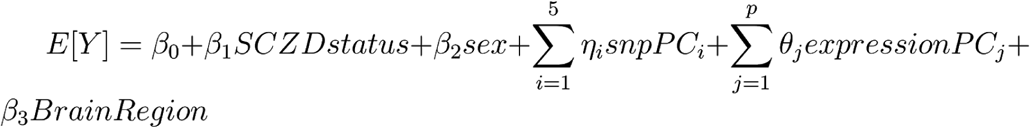

**Equation 11**. Model used for the brain region interaction eQTL analysis between DLPFC and HIPPO. `num. sv ()` from the sva package was used to determine the number of expression PCs to adjust for.

### Risk loci

We downloaded the list of 179 schizophrenia GWAS risk SNPs determined by the Psychiatric Genomics Consortium (Pardiñas et al., 2018). We used this list as input to the rAggr tool available at raaar.usc.edu (Barrett et al., 2005) to identify proxy markers in linkage disequilibrium (LD>0.8) with the list of the 179 index SNPs based on the 1000 Genomes Project Phase 3 database. A maximum distance of 500kb and minumum MAF of 0.001 were used as cutoffs, and reference populations used were African, Americas, East Asian, and European, based on the races present in our samples. Of the 44 index SNPs absent in our genotype data, 28 (64%) had at least one proxy present, so we considered 163 out of 179 risk loci (91%) and 135 index SNPs we directly analyzed. The rAggr software returned a set of 10,981 markers that included both the index and proxy SNPs, of which we found 9,736 of them present in our genotype data (**Figure S32 D**). We then used those 9,736 SNPs to identify eQTLs for each brain region by repeating a similar analysis as described above using **Equation 9** and **Equation 10**, with the exception of now using `pvOutputThreshoid. cis=1` when running `Matrix_eQTL_main ()` for the set of risk SNPs.

### GTEx eQTL replication

We downloaded the GTEx v6 (GTEx Consortium, 2015) HIPPO ( “Brain - Hippocampus “) and DLPFC [ “Brain - Frontal Cortex (BA9) “] fastq files from SRA (Leinonen et al., 2011) using fastq-dump as well as their corresponding genotype data for samples passing the GTEx quality controls (SMAFRZE labeled as “USE ME “). RNA-seq data was processed using the same pipeline and expression features were subsetted to those observed in BrainSeq Phase II. Genotype data was processed as previously described and variantes were subsetted to those observed in BrainSeq Phase II. eQTL analyses were run using matrixEQTL (Shabalin, 2012) for all the expression features and all the variants that had a significant eQTL association (FDR<0.01) in any of the previous 5 eQTL analyses. Replication was assessed for the DLPFC, HIPPO and the brain region interaction eQTLs by directionality or by nominal p-value <0.05.

## Acknowledgements

We thank R. Zielke, R. D. Vigorito and R. M. Johnson of the National Institute of Child Health and Human Development Brain and Tissue Bank for Developmental Disorders at the University of Maryland for providing fetal, child and adolescent brain specimens. The Genotype-Tissue Expression (GTEx) Project was supported by the Common Fund of the Office of the Director of the National Institutes of Health. Additional funds were provided by the NCI, NHGRI, NHLBI, NIDA, NIMH and NINDS. Donors were enrolled at Biospecimen Source Sites funded by NCI/SAIC-Frederick, Inc. (SAIC-F) subcontracts to the National Disease Research Interchange (10XS170), Roswell Park Cancer Institute (10XS171) and Science Care, Inc. (X10S172). The Laboratory, Data Analysis, and Coordinating Center (LDACC) was funded through a contract (HHSN268201000029C) to The Broad Institute, Inc. Biorepository operations were funded through an SAIC-F subcontract to the Van Andel Institute (10ST1035). Additional data repository and project management were provided by SAIC-F (HHSN261200800001E). The Brain Bank was supported by supplements to University of Miami grants DA006227 and DA033684 and to contract N01MH000028. Statistical methods development grants were made to the University of Geneva (MH090941 and MH101814), the University of Chicago (MH090951, MH090937, MH101820, MH101825), the University of North Carolina - Chapel Hill (MH090936 and MH101819), Harvard University (MH090948), Stanford University (MH101782), Washington University St Louis (MH101810) and the University of Pennsylvania (MH101822). The data used for the analyses described in this manuscript were obtained from dbGaP accession number phs000424.v6.p1 on October 6, 2015.

*Members of the BrainSeq consortium include*: Mitsuyuki Matsumoto, Takeshi Saito, Katsunori Tajinda, Daniel Hoeppner, Nicholas J. Brandon, Alan Cross, David Andrew Collier, John N Calley, Cara Lee Ann Ruble, Brain J Eastwood, Philip J Ebert, David Charles Airey, Yupeng Li, Yushi Liu, Karim Malki, Bradley Bryan Miller, James E Scherschel, Hong Wang, Maura Furey, Derrek Hibar, Hartmuth Kolb, Michael Didriksen, Lasse Folkersen, Tony Kam-Thong, Dheeraj Malhotra, Patricio O’Donnell, Simon (Hualin) Xi, Jie Quan, Joo Heon Shin, Andrew E. Jaffe, Rujuta Narurkar, Richard E. Straub, Amy Deep-Soboslay, Thomas M. Hyde, Joel E. Kleinman and Daniel R. Weinberger.

## Author contributions

- L.C.-T.: Conceptualization, Formal Analysis, Methodology, Software, Supervision, Visualization, Writing – Original Draft Preparation, Writing – Review & Editing
- E.E.B.: Formal Analysis, Software, Visualization, Writing – Original Draft Preparation, Writing – Review & Editing
- A.P.: Formal Analysis, Writing – Original Draft Preparation
- J.H.S.: Investigation, Supervision
- R.E.S.: Conceptualization, Methodology, Supervision, Writing – Review & Editing
- R.A.: Investigation
- S.A.S.: Formal Analysis, Visualization, Writing – Review & Editing
- W.S.U.: Visualization
- BrainSeq consortium: Conceptualization, Funding Acquisition, Project Administration
- C.V.: Formal Analysis
- R.T.: Investigation
- A.D.-S.: Data Curation
- T.M.H.: Conceptualization, Investigation, Supervision
- J.E.K.: Conceptualization, Data Curation, Supervision
- D.R.W.: Conceptualization, Funding Acquisition, Methodology, Supervision, Writing – Review & Editing
- A.E.J.: Conceptualization, Formal Analysis, Funding Acquisition, Methodology, Supervision, Writing – Original Draft Preparation, Writing – Review & Editing

## Data availability and materials

Raw and processed data is available from ^*****^**TBD** Code is available through GitHub at https://aithub.com/Lieberlnstitute/brainseg_phase2 and https://github.com/Lieberlnstitute/osva_brain.

---

**Figure S1.**
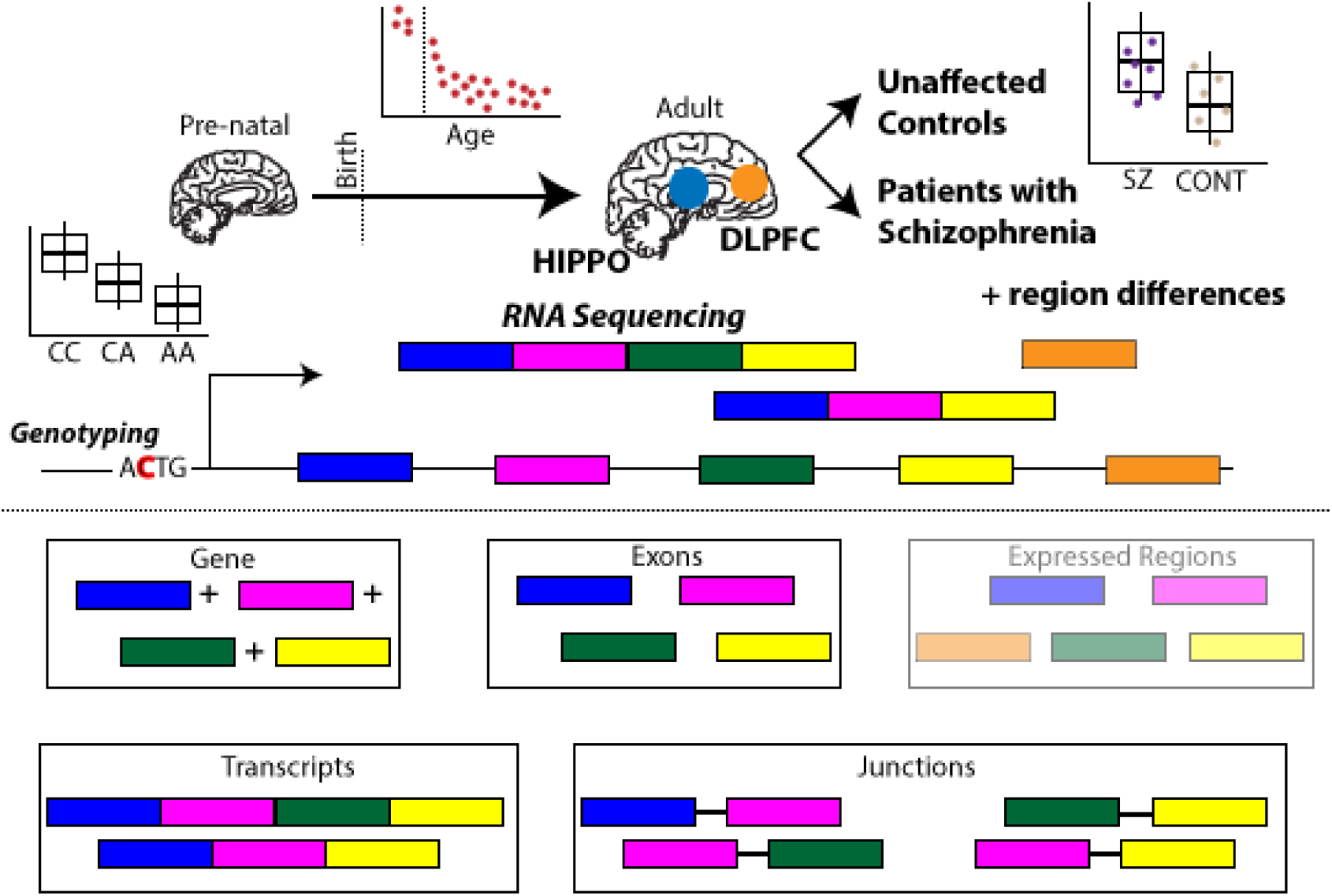
BrainSeq Phase II RNA-seq overview. RNA-seq samples from the hippocampus (HIPPO) and dorsolateral prefrontal cortex (DLPFC) across the human lifespan were sequenced. The individuals were also genotyped with some being patients with SCZD and others being non-psychiatric controls. Differences in expression across age, across HIPPO and DLPFC and between SCZD status were assessed. Expression was evaluated at 4 feature levels: gene, exons, exon-exon junctions and transcripts. The data is also available to perform expressed regions analyses.

**Figure S2.**
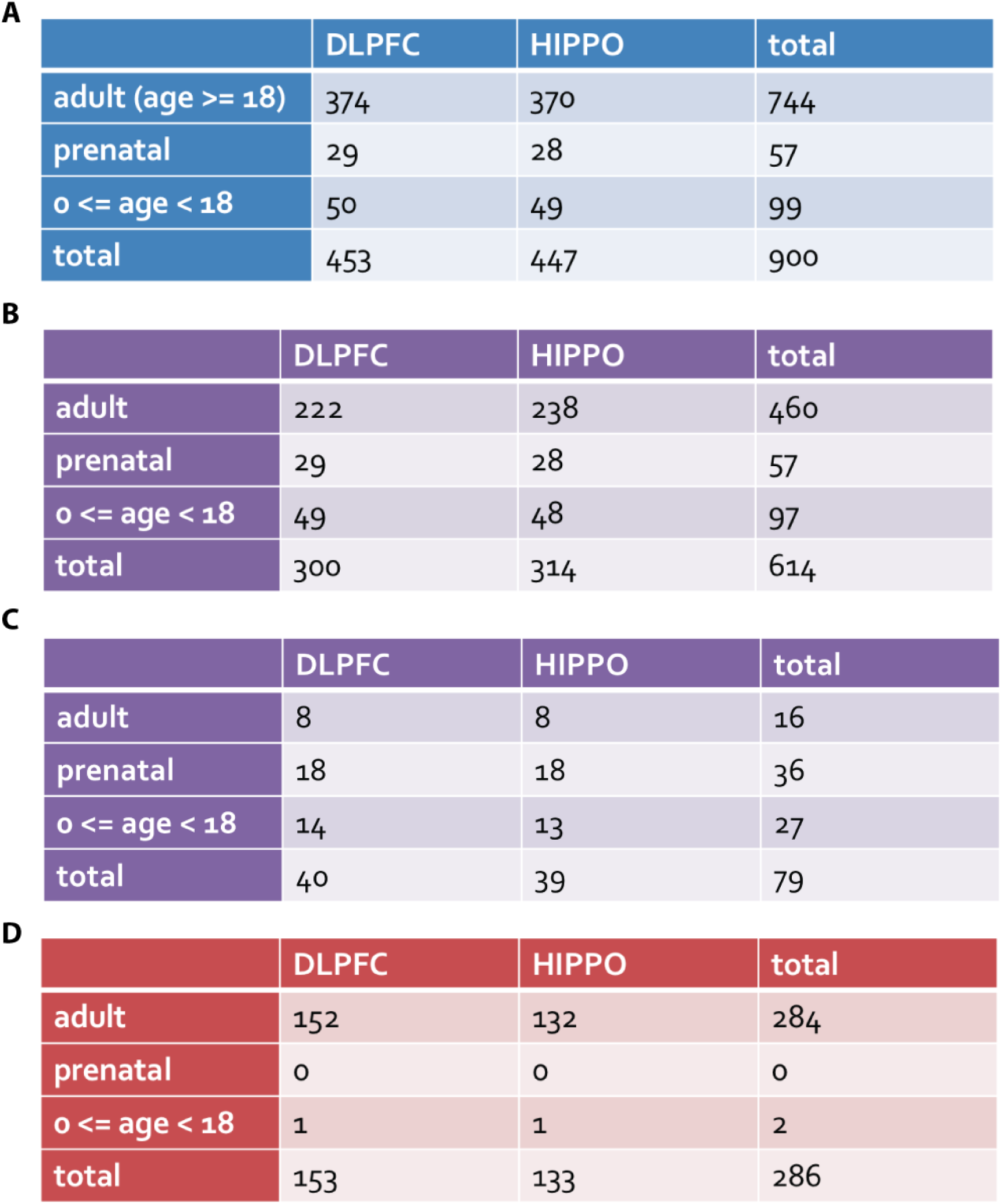
RNA-seq samples in BrainSeq Phase II and BrainSpan. (**A**) All BrainSeq Phase II RNA-seq samples. (**B**) Non-psychiatric control samples in BrainSeq Phase II. (C) Non-psychiatric control samples in BrainSpan. (**D**) SCZD samples in BrainSeq Phase II.

**Figure S3.**
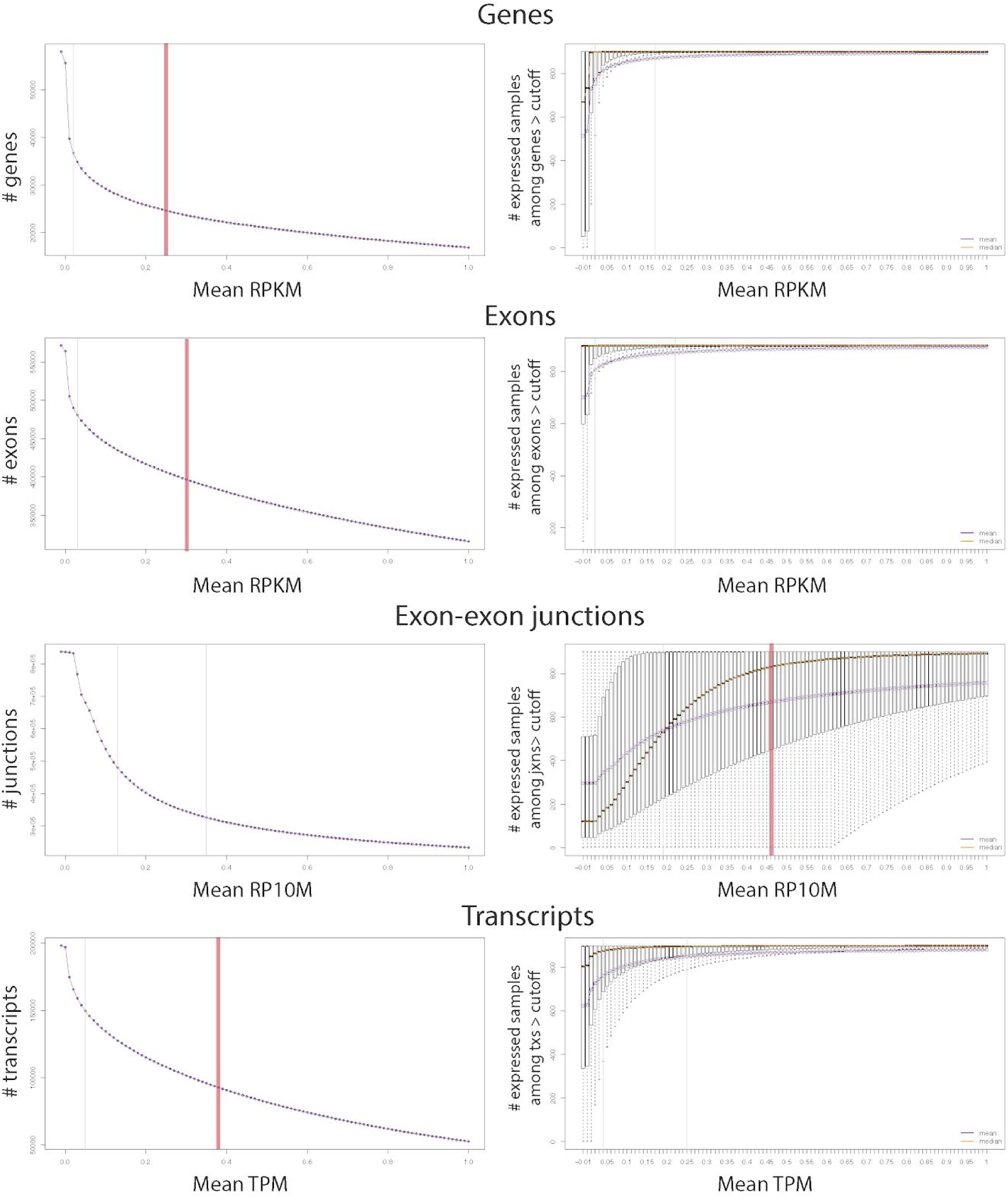
Determination of the low expression filters. (**Left**) Mean expression and the number of features passing the cutoff. (**Right**) Mean expression compared to the number of expressed samples (non-zero expression) among the features that pass the cutoff. The maximum suggested cutoff (highlighted in red) was used for each feature type.

**Figure S4.**
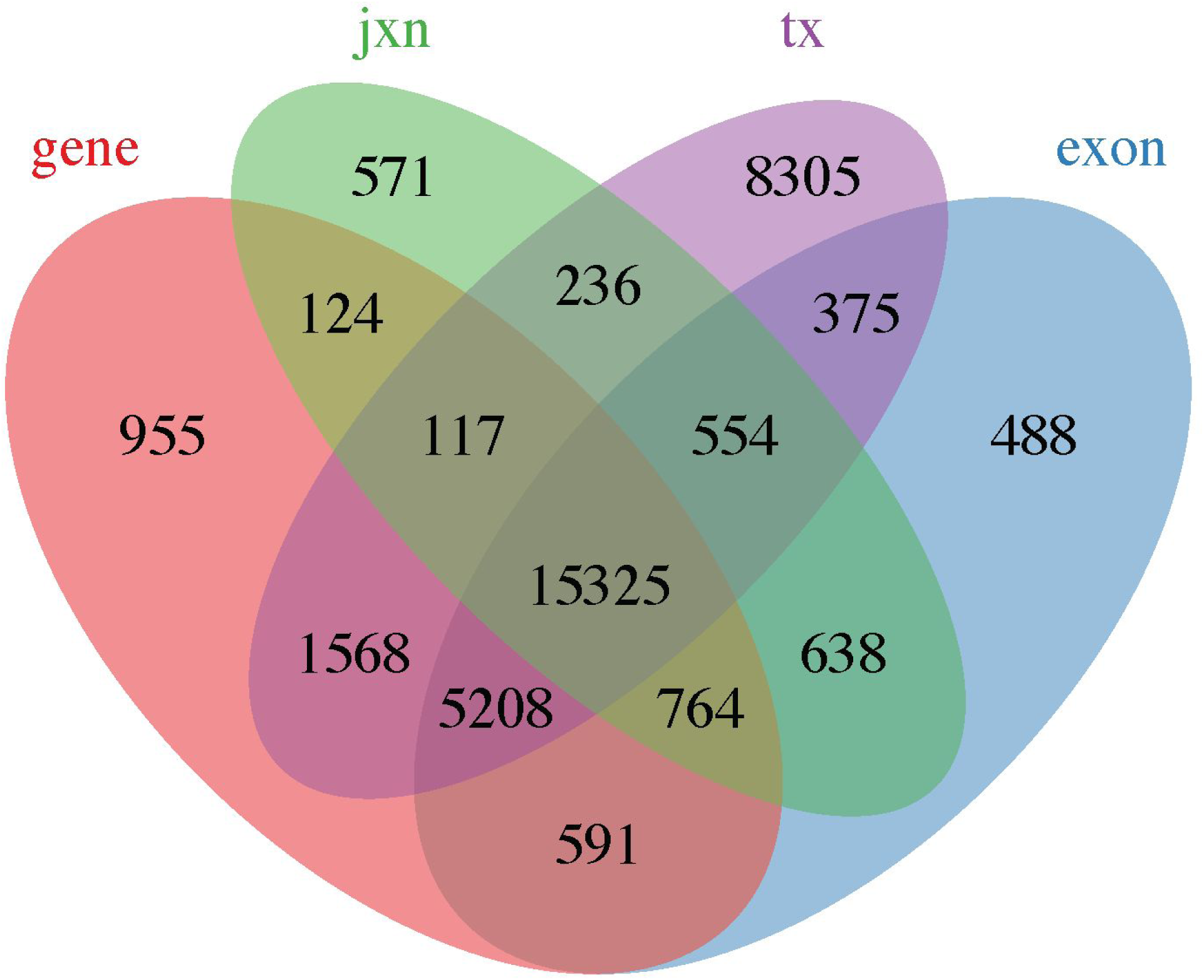
Expressed features grouped by gene ID. Venn diagram of expressed features grouped by Ensembl gene ID.

**Figure S5.**
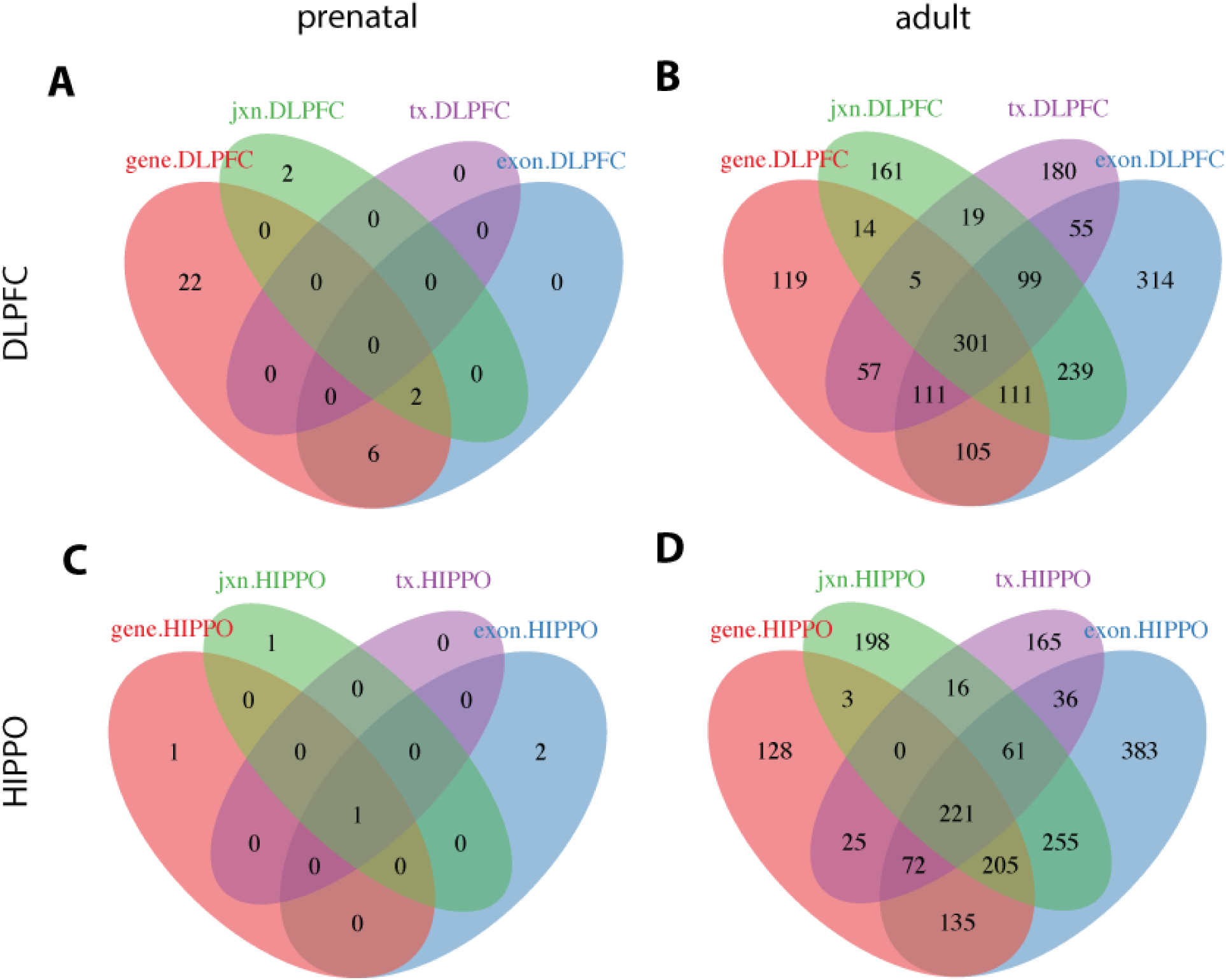
Region-specific differentially expressed features. Venn diagrams of differentially expressed features between DLPFC and HIPPO grouped by Ensembl gene ids. (**A**) DE features in prenatal samples with higher expression in DLPFC. (**B**) DE features in adult samples with higher expression in DLPFC. (**C**) DE features in prenatal samples with higher expression in HIPPO. (**D**) DE features in adult samples with higher expression in HIPPO.

**Figure S6.**
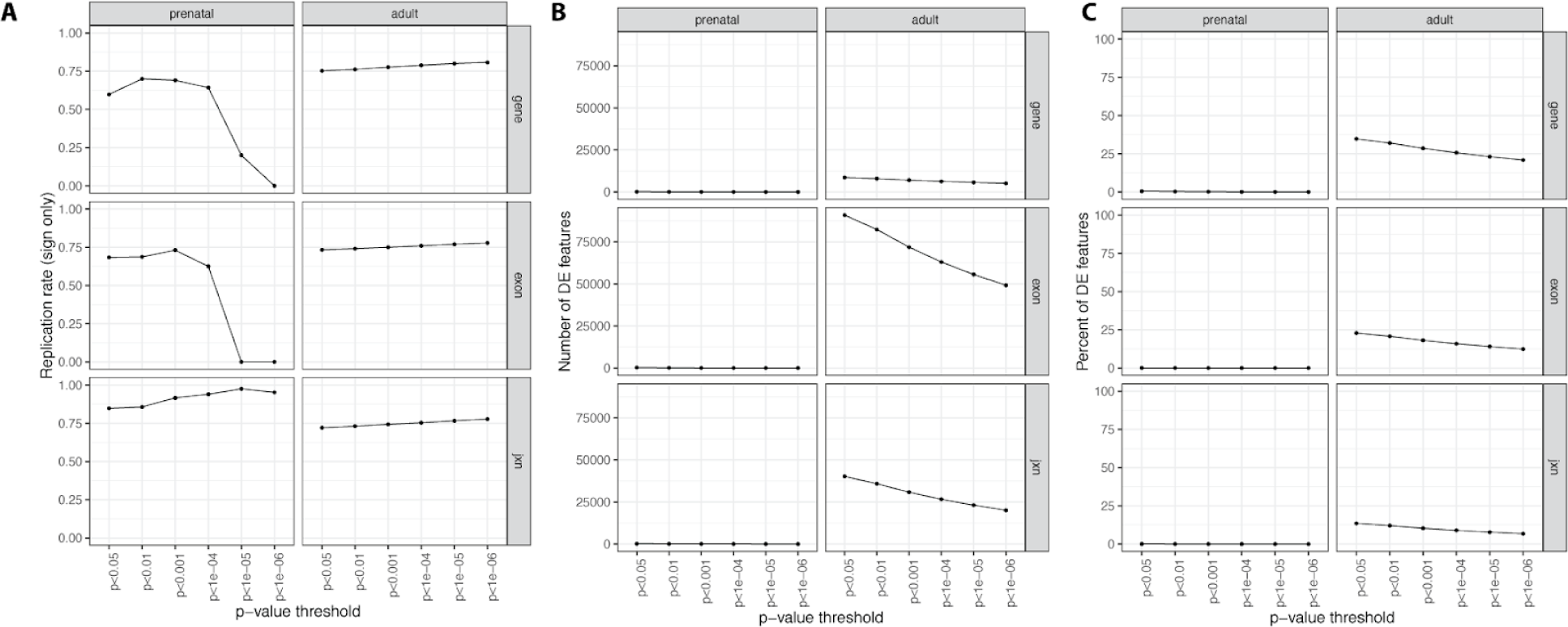
Region-specific differentially expressed replication rate with BrainSpan. (**A**) Replication rate when considering directionality only instead of consistent directionality and p-value < 0.05. (**B**) Number and percent (**C**) of differentially expressed features across the different p-value thresholds.

**Figure S7.**
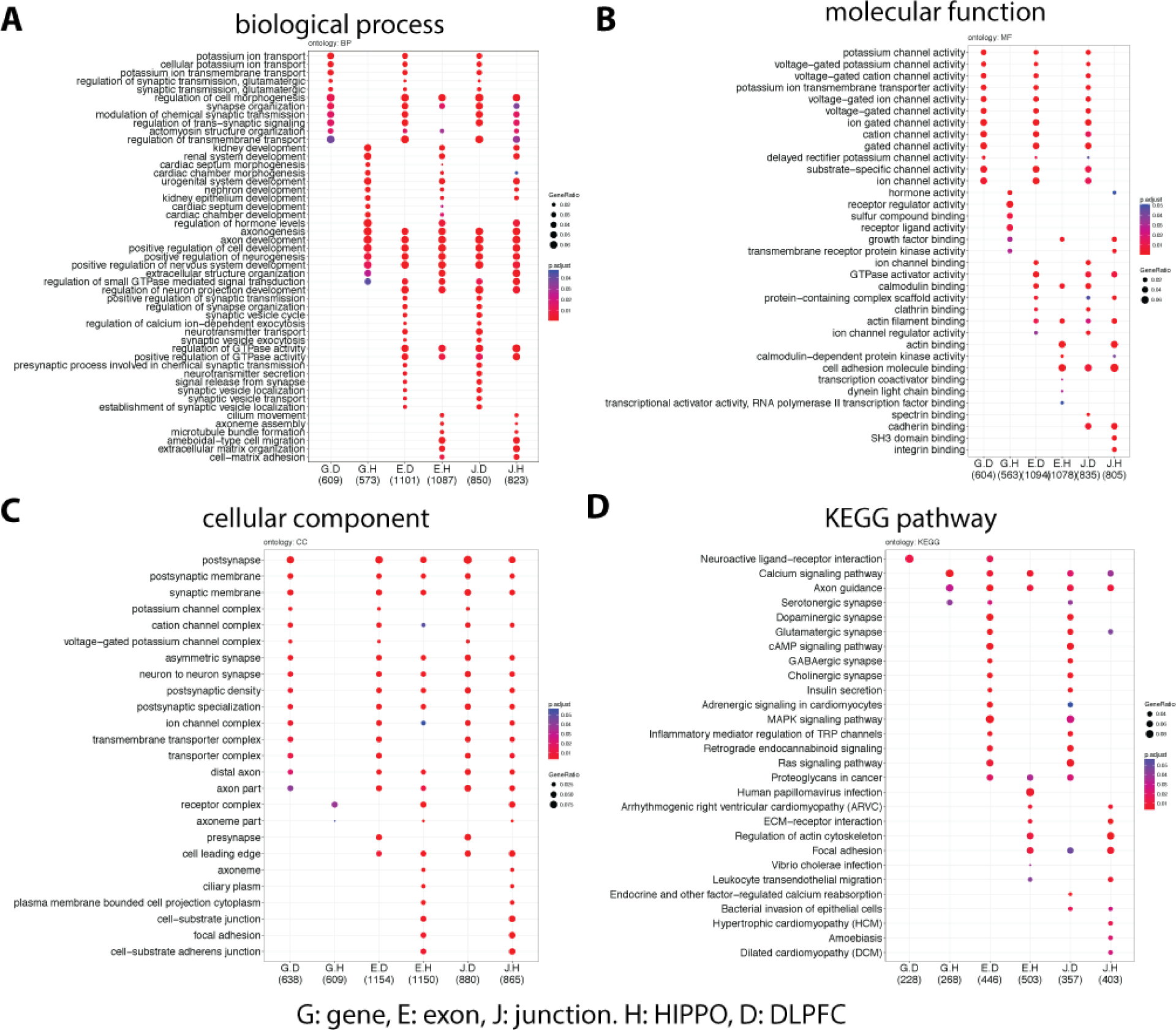
Enriched GO terms for region-specific adult genes. Gene ontology enrichment results for differentially expressed features between HIPPO and DLPFC in adult samples. Enriched (**A**) biological processes, (**B**) molecular function, (**C**) cellular component ontologies and (**D**) Kyoto Encyclopedia of Genes and Genomes (KEGG) pathways at the gene, exon or junction feature levels separated by higher expression in HIPPO or DLPFC.

**Figure S8.**
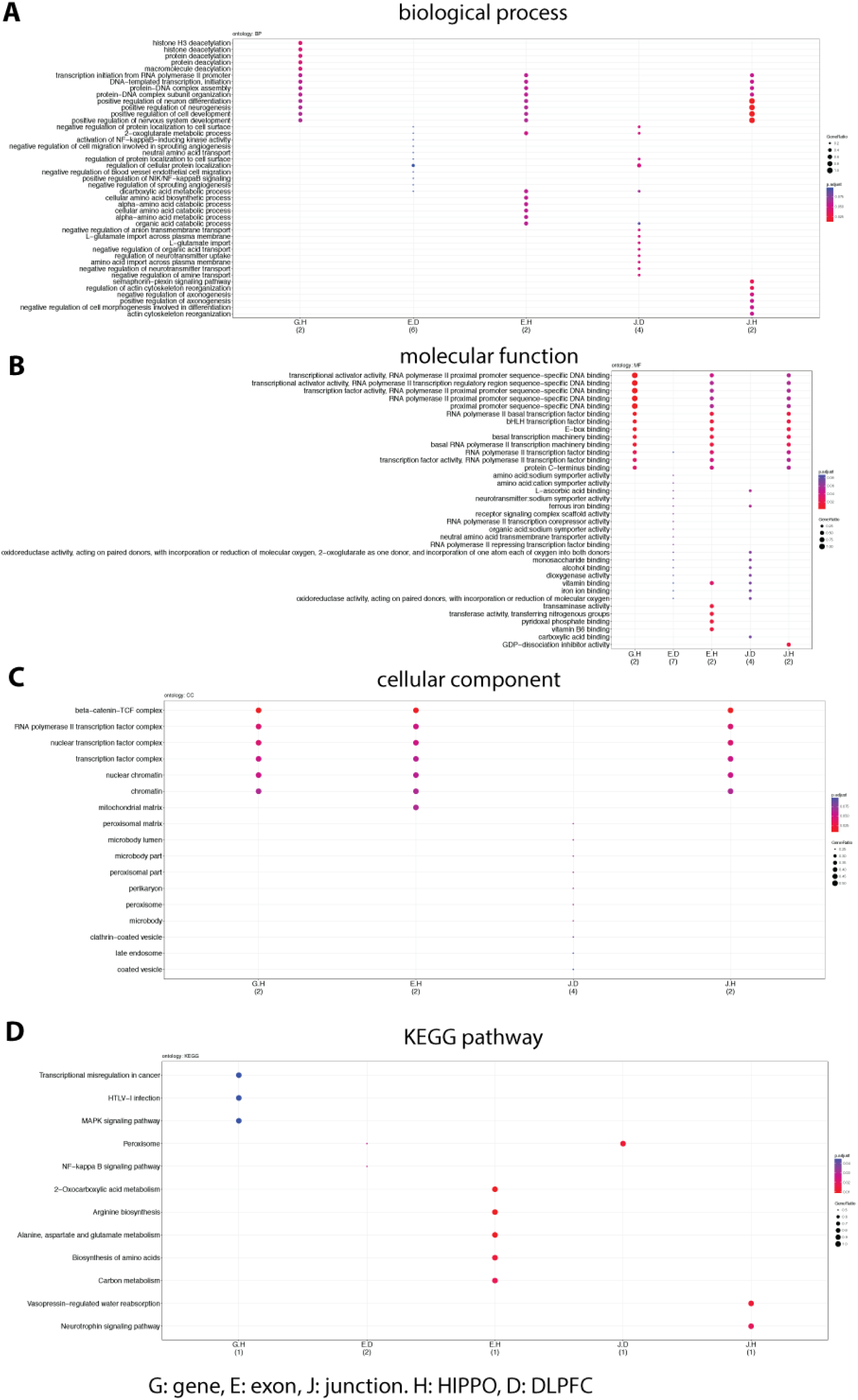
Enriched GO terms for region-specific prenatal genes. Gene ontology enrichment results for differentially expressed features between HIPPO and DLPFC in prenatal samples. Enriched (**A**) biological processes, (**B**) molecular function, (**C**) cellular component ontologies and (**D**) Kyoto Encyclopedia of Genes and Genomes (KEGG) pathways at the gene, exon or junction feature levels separated by higher expression in HIPPO or DLPFC.

**Figure S9.**
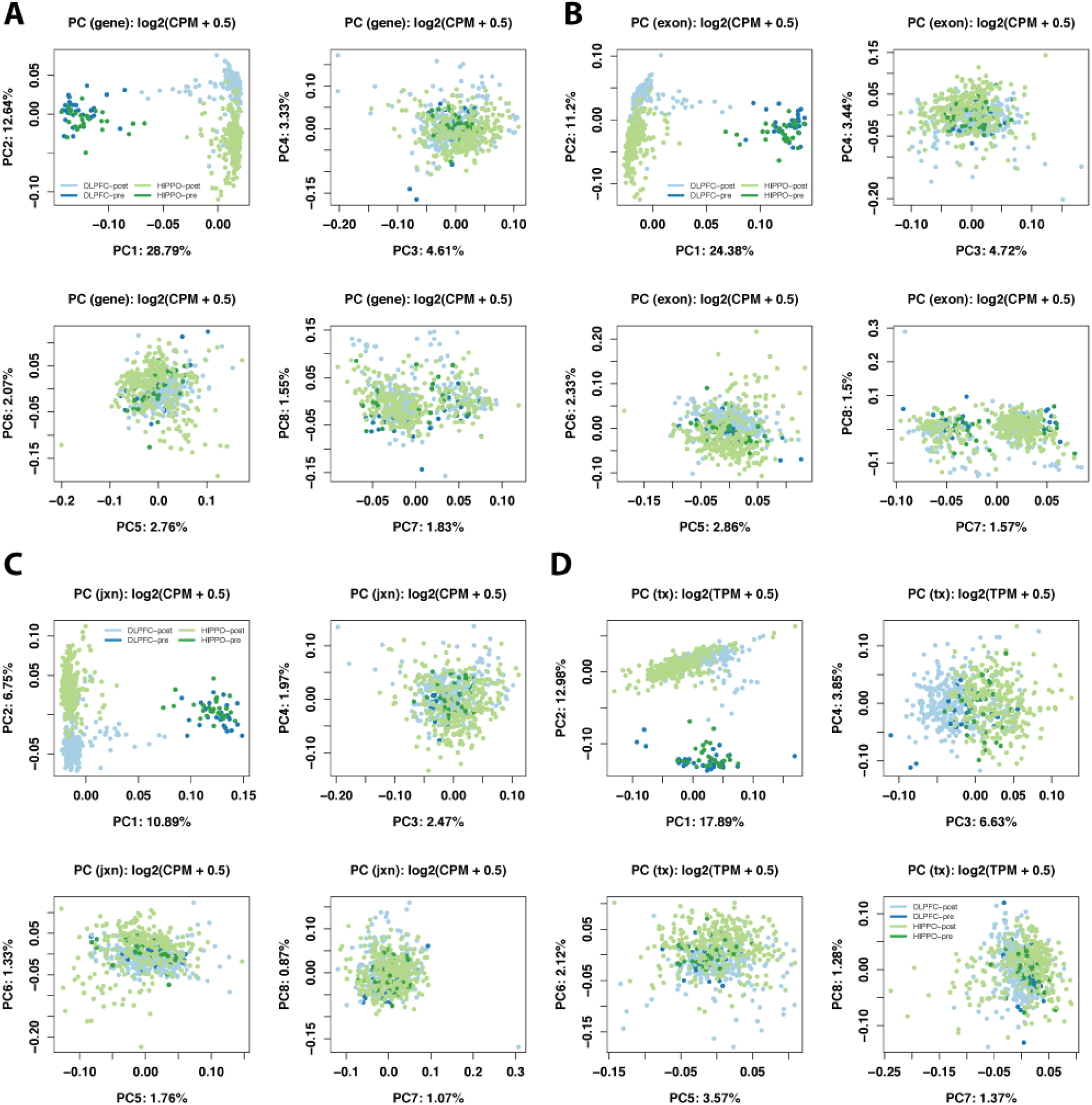
Top PCs by brain region and age. Principal components for each feature across the 614 samples used for the development analysis colored by pre- and postnatal status as well as brain region. Top 8 PCs for (**A**) gene, (**B**) exon, (**C**) exon-exon junctions and (**D**) transcripts.

**Figure S10.**
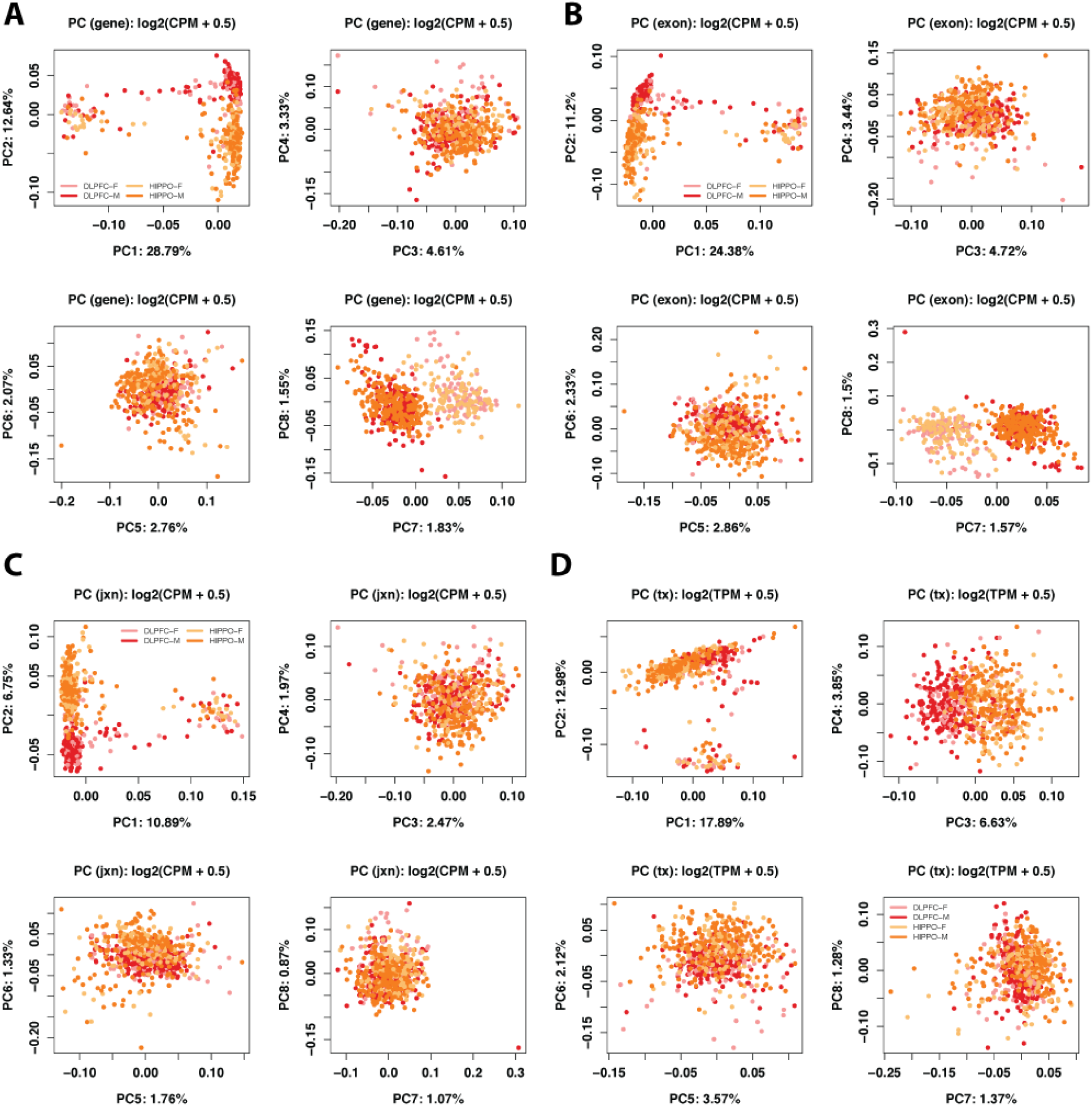
Top PCs by brain region and sex. Principal components for each feature across the 614 samples used for the development analysis colored by sex and brain region. Top 8 PCs for (**A**) gene, (**B**) exon, (**C**) exon-exon junctions and (**D**) transcripts.

**Figure S11.**
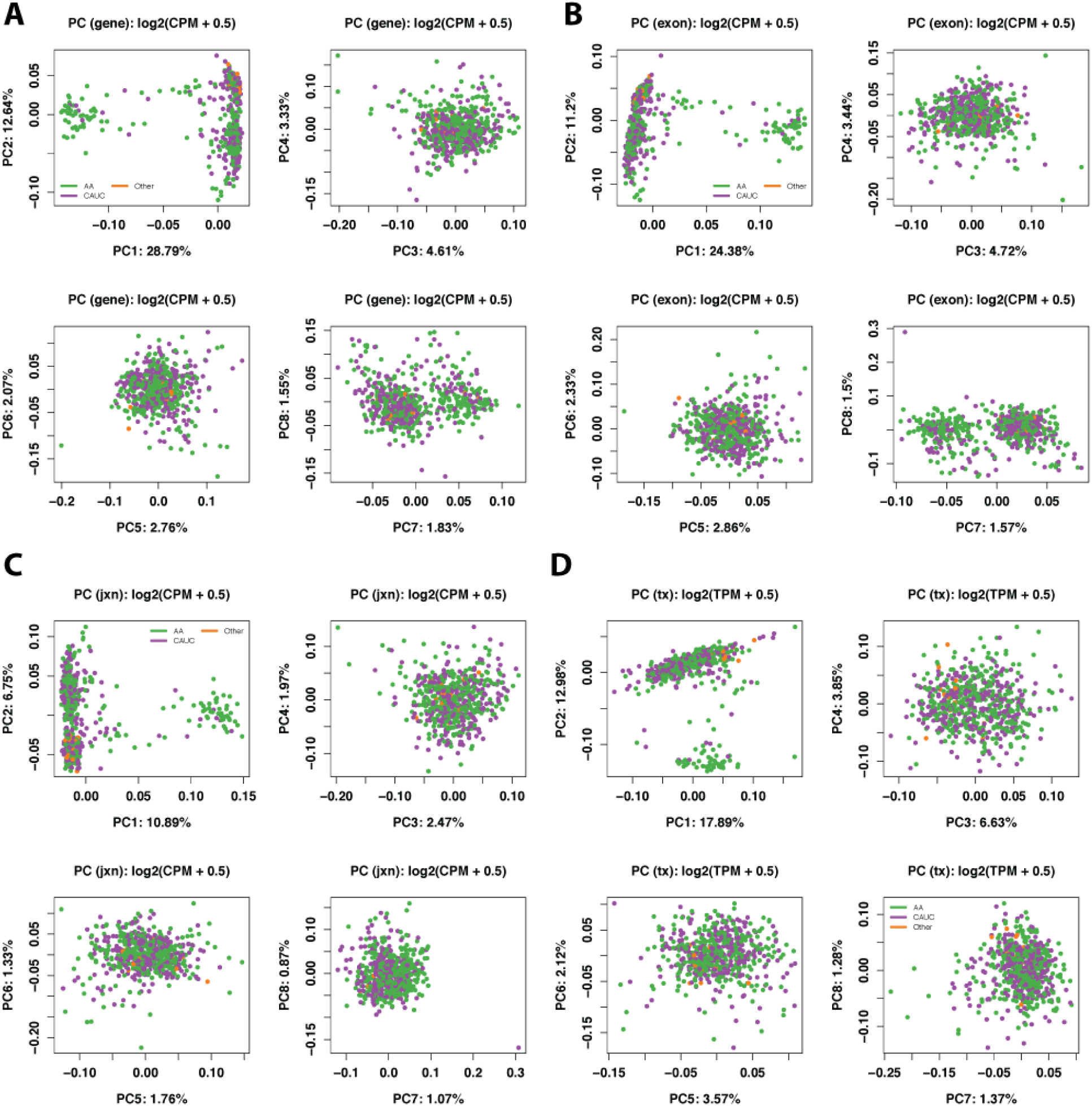
Top PCs by race. Principal components for each feature across the 614 samples used for the development analysis colored by race. Top 8 PCs for (**A**) gene, (**B**) exon, (**C**) exon-exon junctions and (**D**) transcripts.

**Figure S12.**
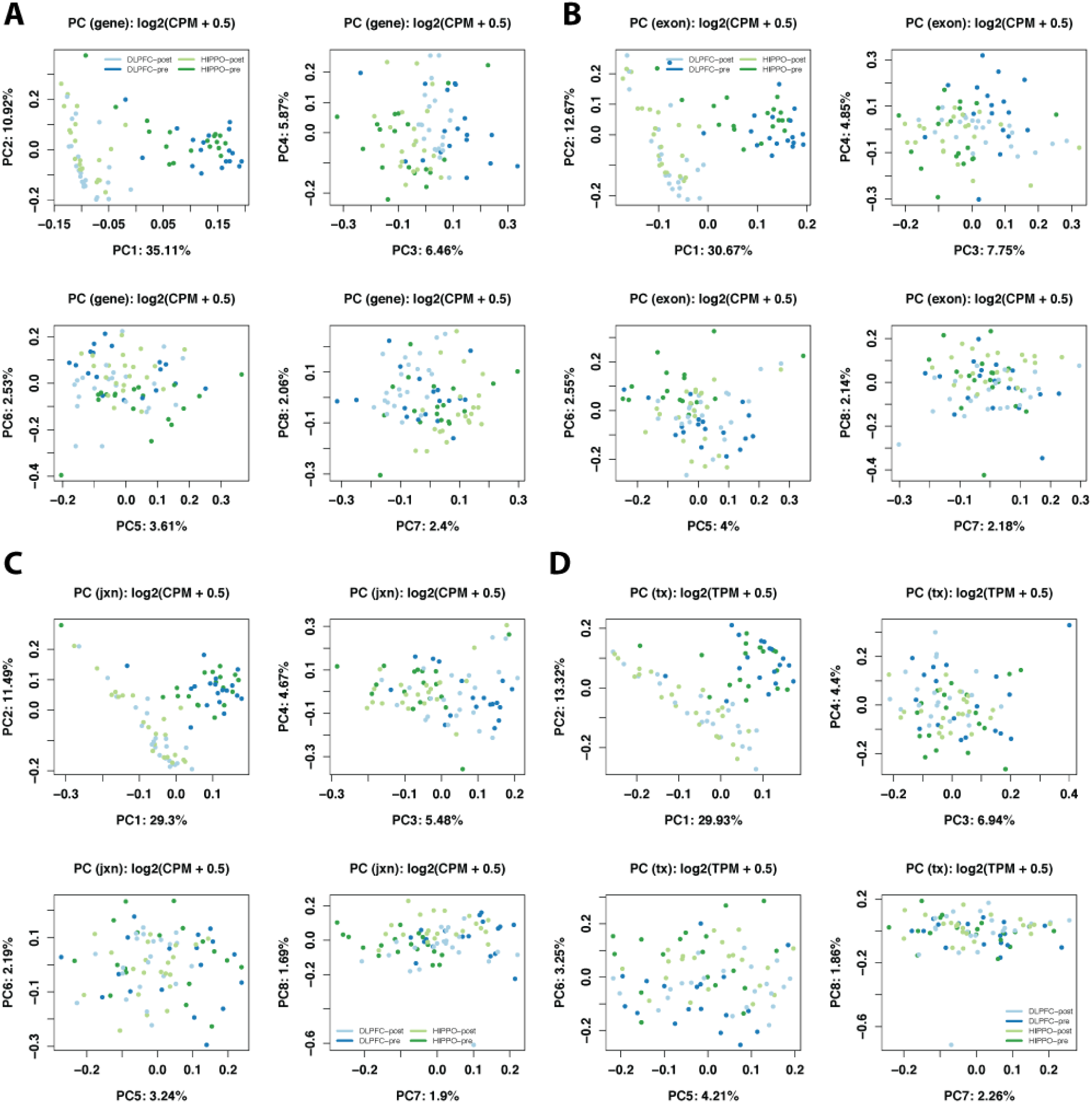
Top PCs by brain region and age in BrainSpan. Principal components for each feature across 79 the BrainSpan samples used for the development replication analysis colored by pre- and postnatal status as well as brain region. Top 8 PCs for (**A**) gene, (**B**) exon, (**C**) exon-exon junctions and (**D**) transcripts.

**Figure S13.**
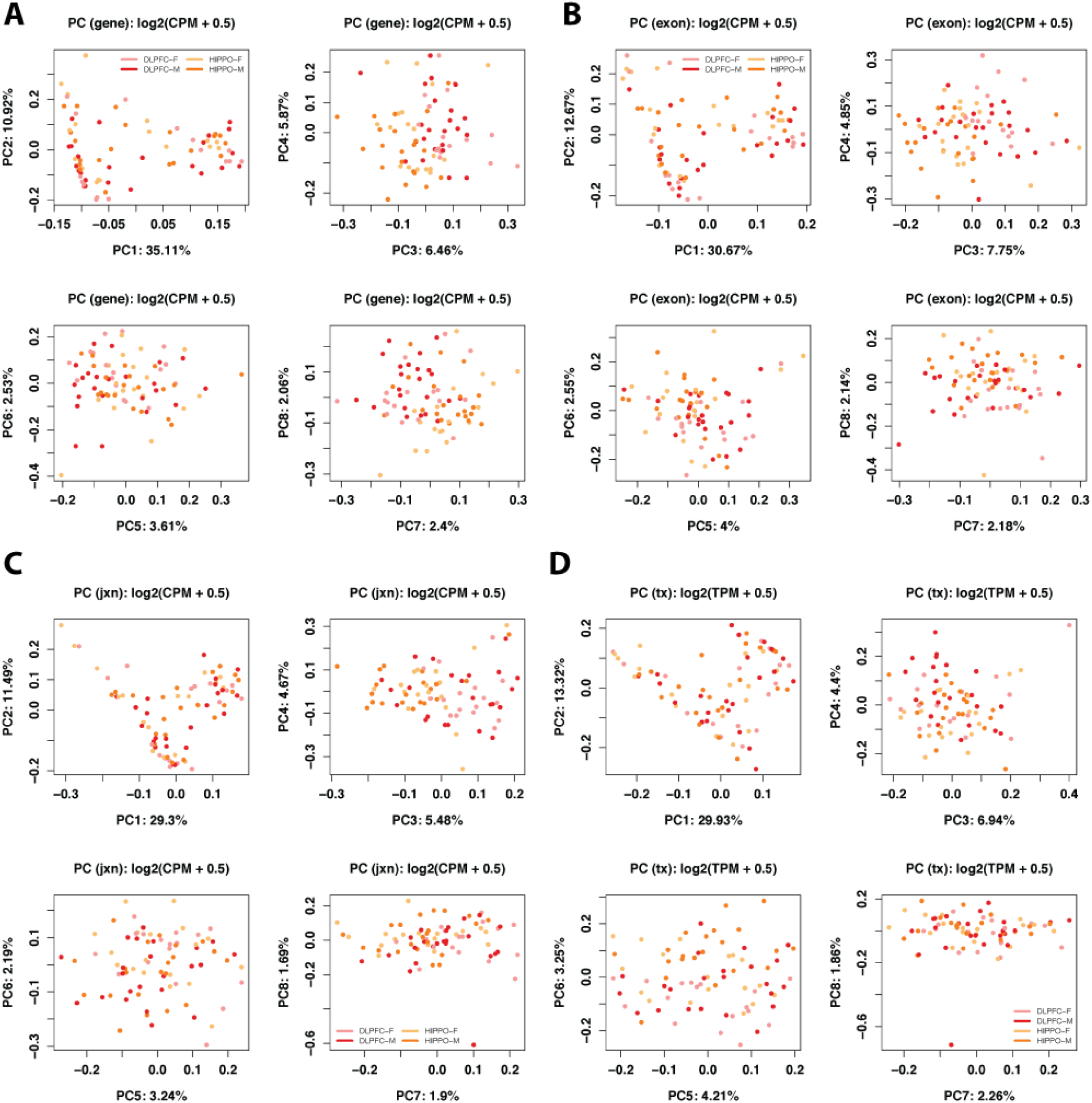
Top PCs by brain region and sex in BrainSpan. Principal components for each feature across the 79 BrainSpan samples used for the development analysis colored by sex and brain region. Top 8 PCs for (**A**) gene, (**B**) exon, (**C**) exon-exon junctions and (**D**) transcripts.

**Figure S14.**
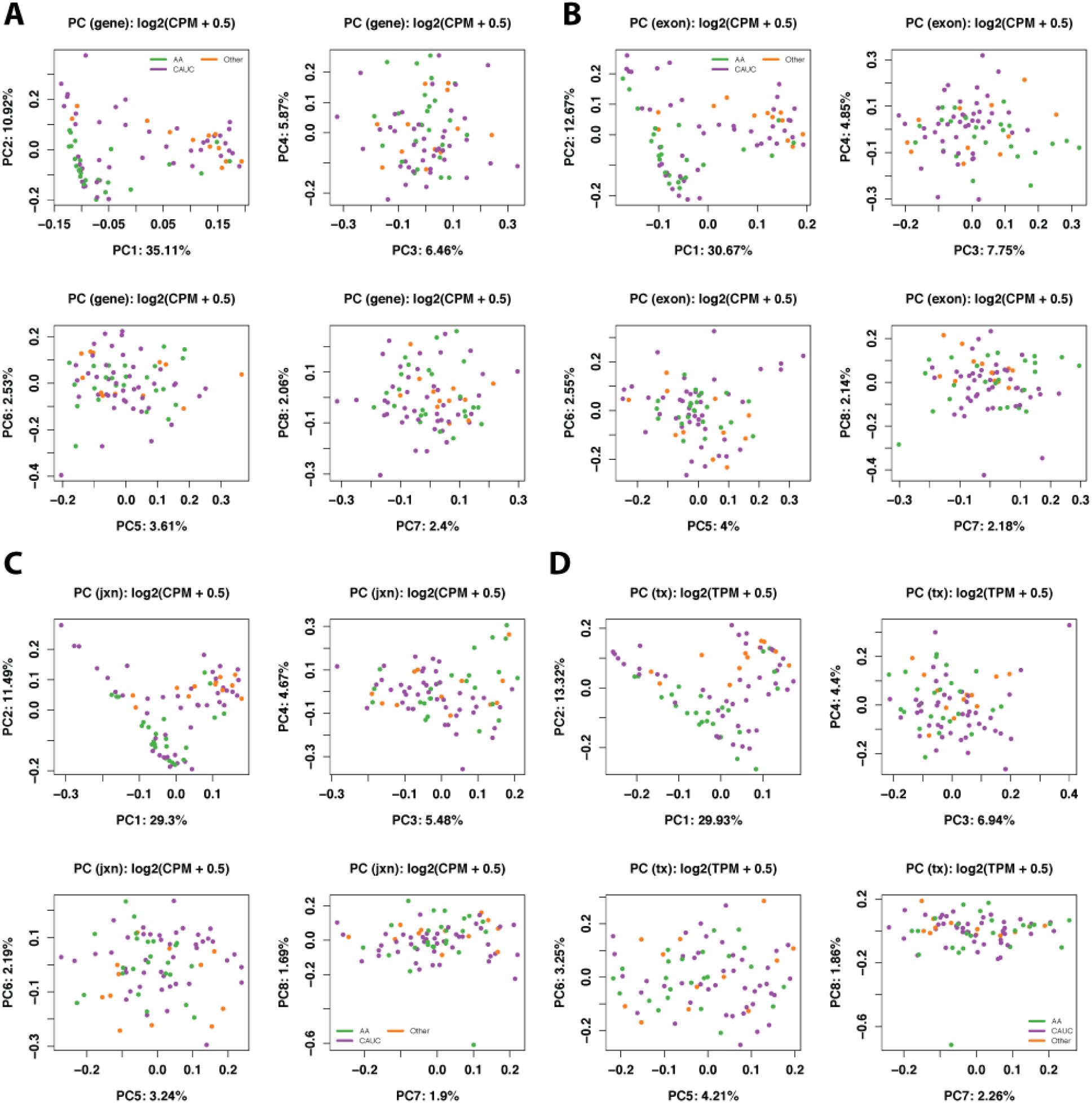
Top PCs by race in BrainSpan. Principal components for each feature across the 79 BrainSpan samples used for the development replication analysis colored by race. Top 8 PCs for (**A**) gene, (**B**) exon, (**C**) exon-exon junctions and (**D**) transcripts.

**Figure S15.**
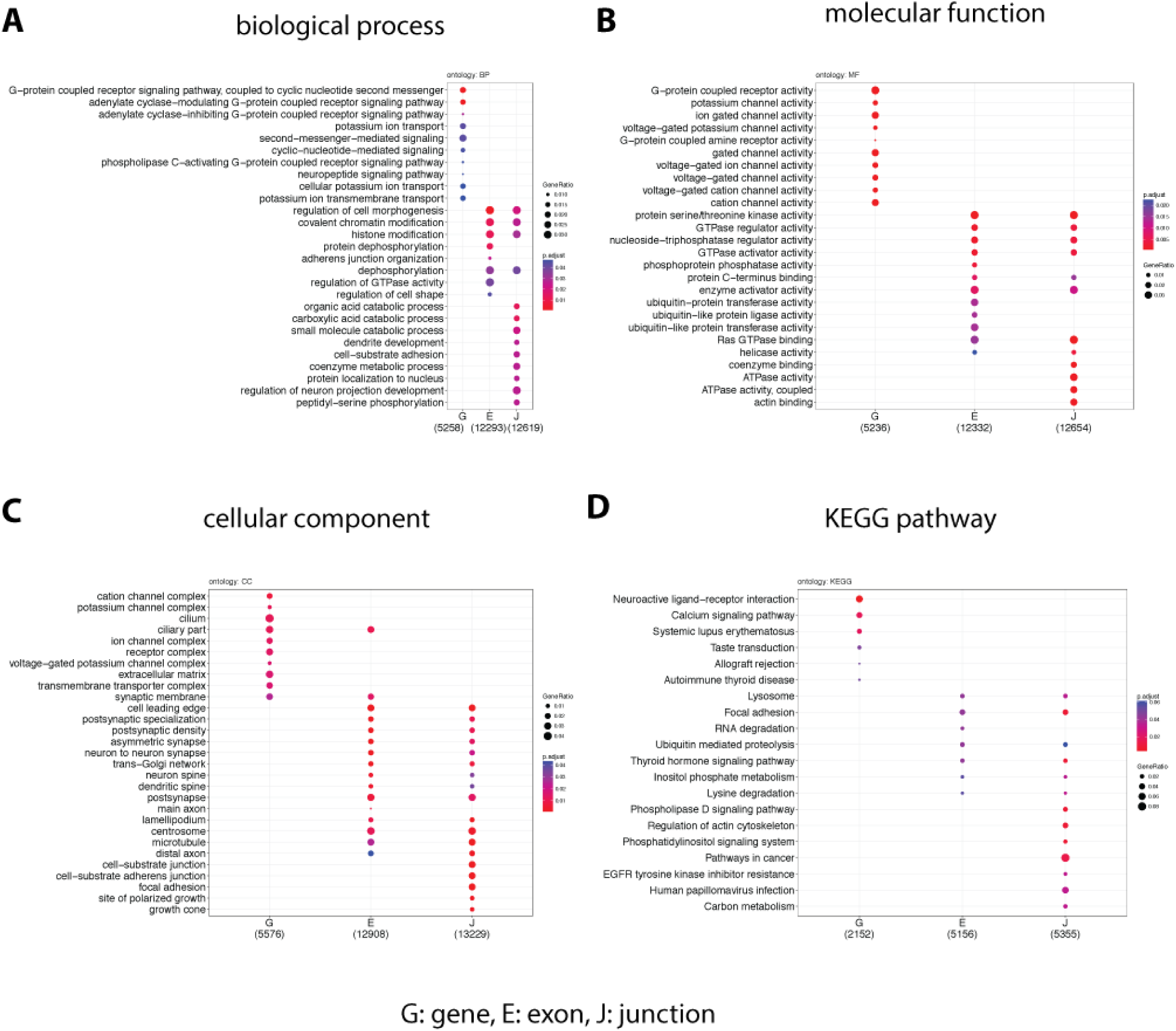
Enriched GOs across development. Gene ontology enrichment results for differentially expressed features across development as modeled with age splines. Enriched (**A**) biological processes, (**B**) molecular function, (**C**) cellular component ontologies and (**D**) Kyoto Encyclopedia of Genes and Genomes (KEGG) pathways at the gene, exon or junction feature levels.

**Figure S16.**
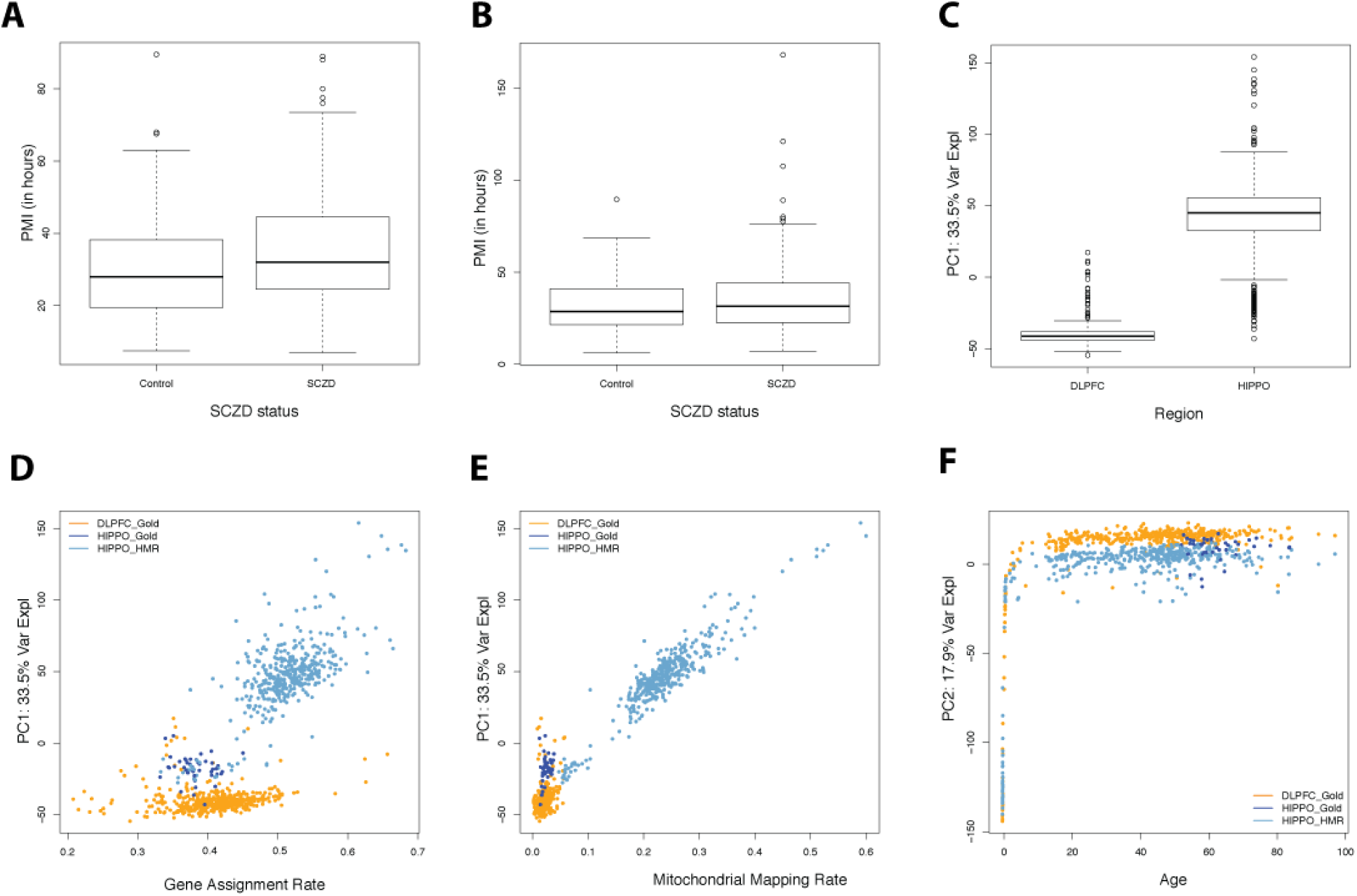
PMI and PCA EDA with all 900 samples. Post mortem interval (PMI) measured in hours for (**A**) HIPPO and (**B**) DLPFC separated by SCZD status. Using all 900 BrainSeq Phase II RNA-seq samples, the first gene-level principal component (PC) is associated with brain region (**C**), gene assignment rate (**D**) and mitochondrial mapping rate (**E)**. The second PC is associated with age (**F**), in particular with the prenatal vs postnatal age boundary. Samples in the bottom row are colored by brain region and RNA-seq preparation kit.

**Figure S17.**
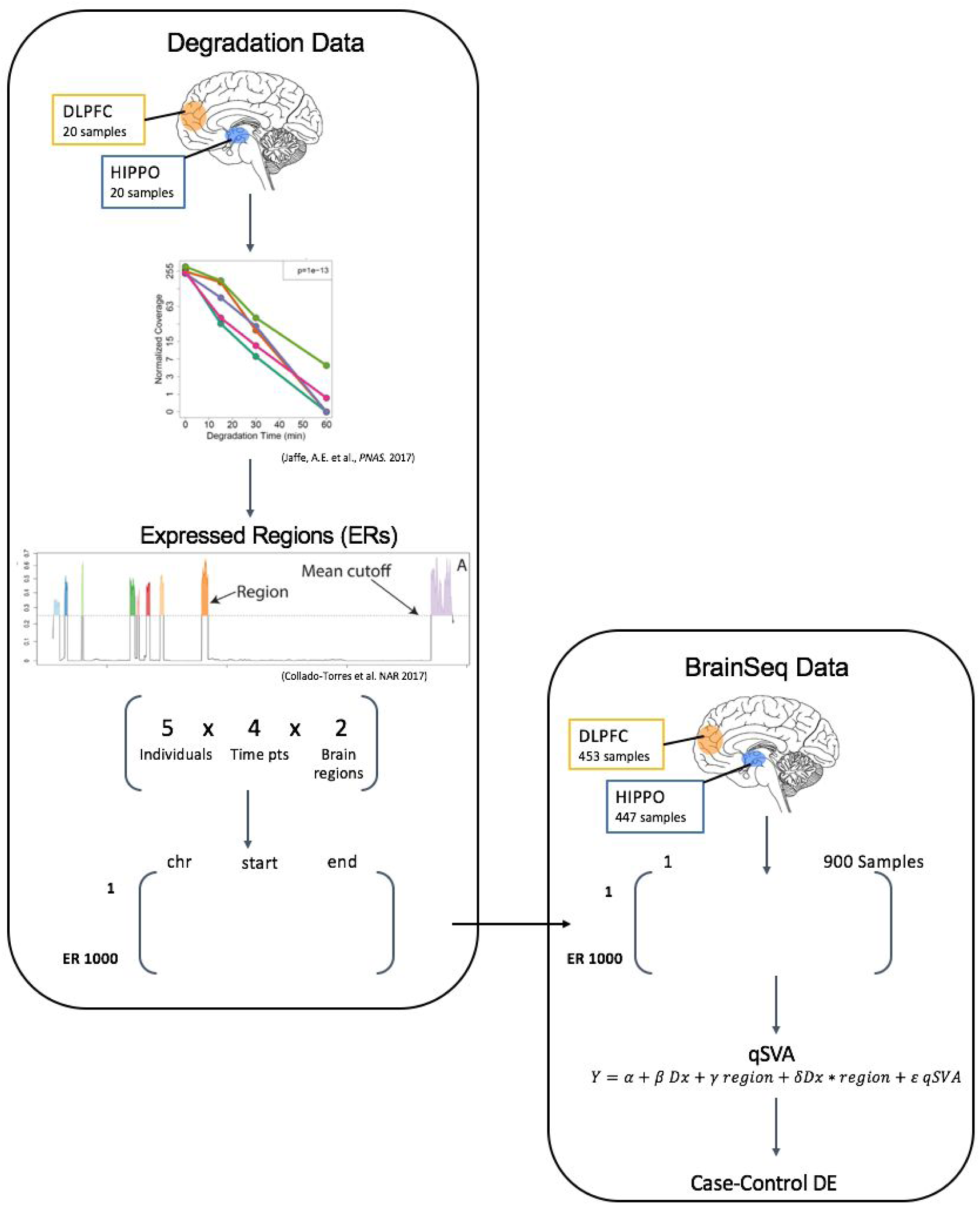
qSVA overview. Workflow illustration for a complete analysis using degradation data and case-control data to implement the qSVA method. The degradation data, a few samples across time, are used to infer expressed regions that are associated with degradation. We save the genome coordinates of the top 1000 regions. We then quantify the expression of those regions in our case-control data (BrainSeq Phase II) to obtain a degradation matrix. Using this matrix we compute the quality surrogate variables (qSVs) that we can then use as adjustment covariates in a case-control differential expression analysis. We identify the qSVs by performing PCA on the degradation matrix and choosing k number with the BE algorithm (Buja and Eyuboglu, 1992).

**Figure S18.**
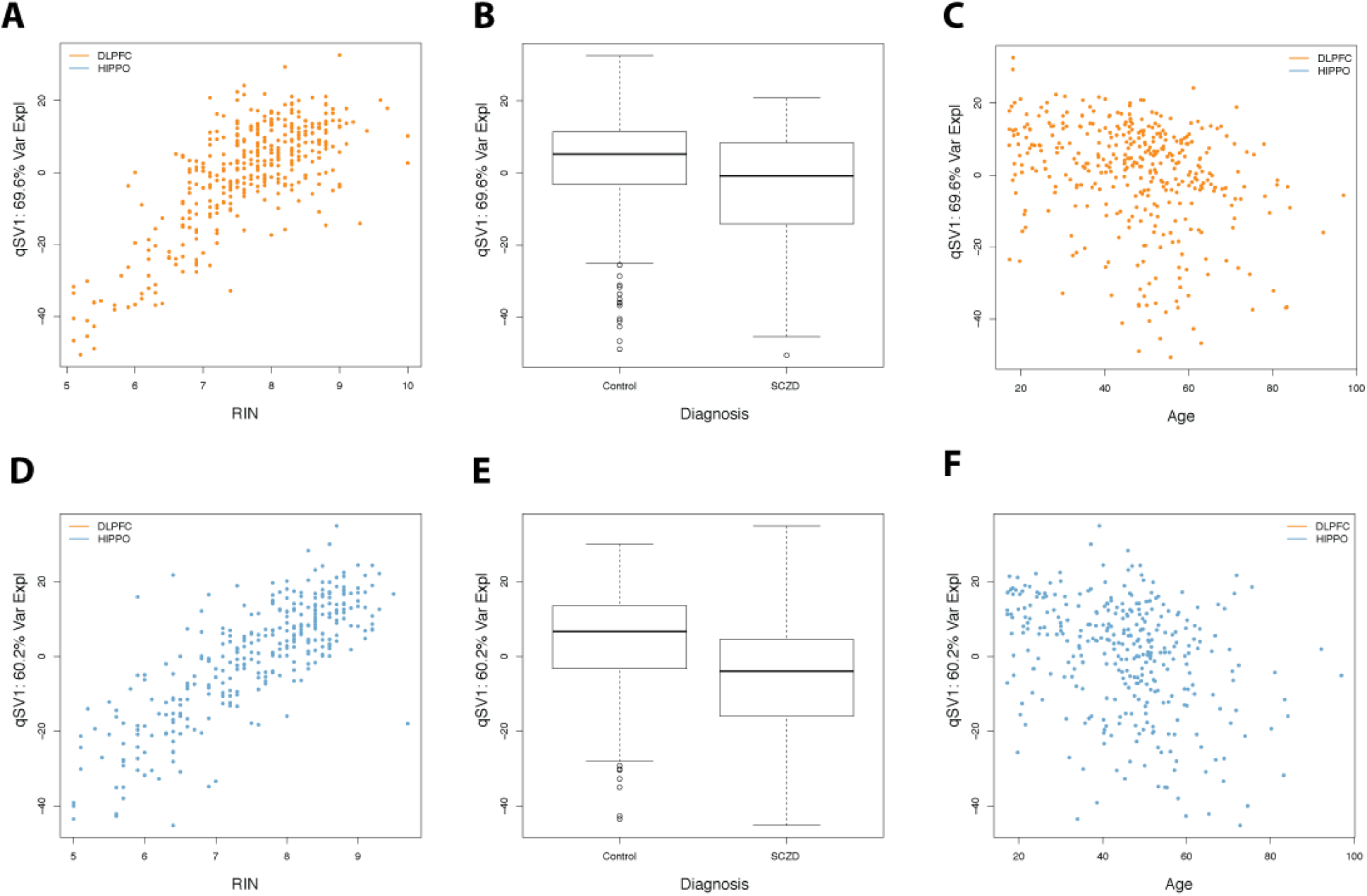
qSVA EDA for HIPPO and DLPFC. (**A**) First quality surrogate variable (qSV) from the DLPFC samples (n=379) is associated with RIN and SCZD diagnosis status (**B**). Similarly, the first qSV from the HIPPO samples (n=333) is associated with RIN (**D**) and SCZD diagnosis status (**E**). There is no strong association with age for the first qSV for both DLPFC (**C**) and HIPPO (**F**).

**Figure S19.**
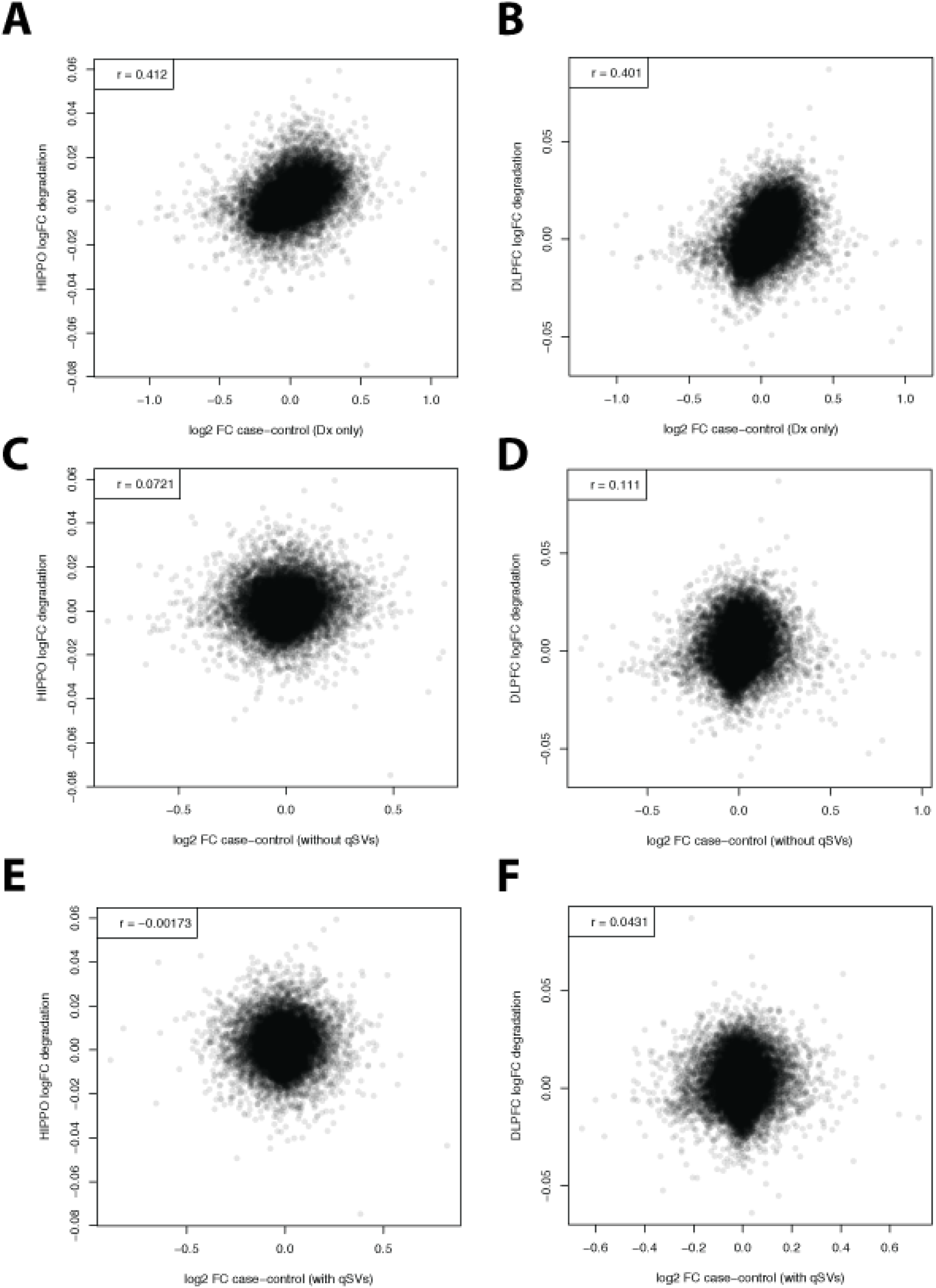
DEqual plots. Degradation quality (DEqual) plots for each brain region (HIPPO: left column, DLPFC: right column) comparing against a naïve diagnosis model (**A** and **B**), adjusting for covariates including known quality-associated metrics (**C** and **D**) and the final model adjusting for the region-specific qSVs and covariates (**E** and **F**). Correlation between the log2 fold change by SCZD status (x-axis) and log2 fold change by degradation in HIPPO or DLPFC is shown in the top left corner of each panel. DEqual plots are defined in Jaffe et al. (Jaffeetal., 2017).

**Figure S20.**
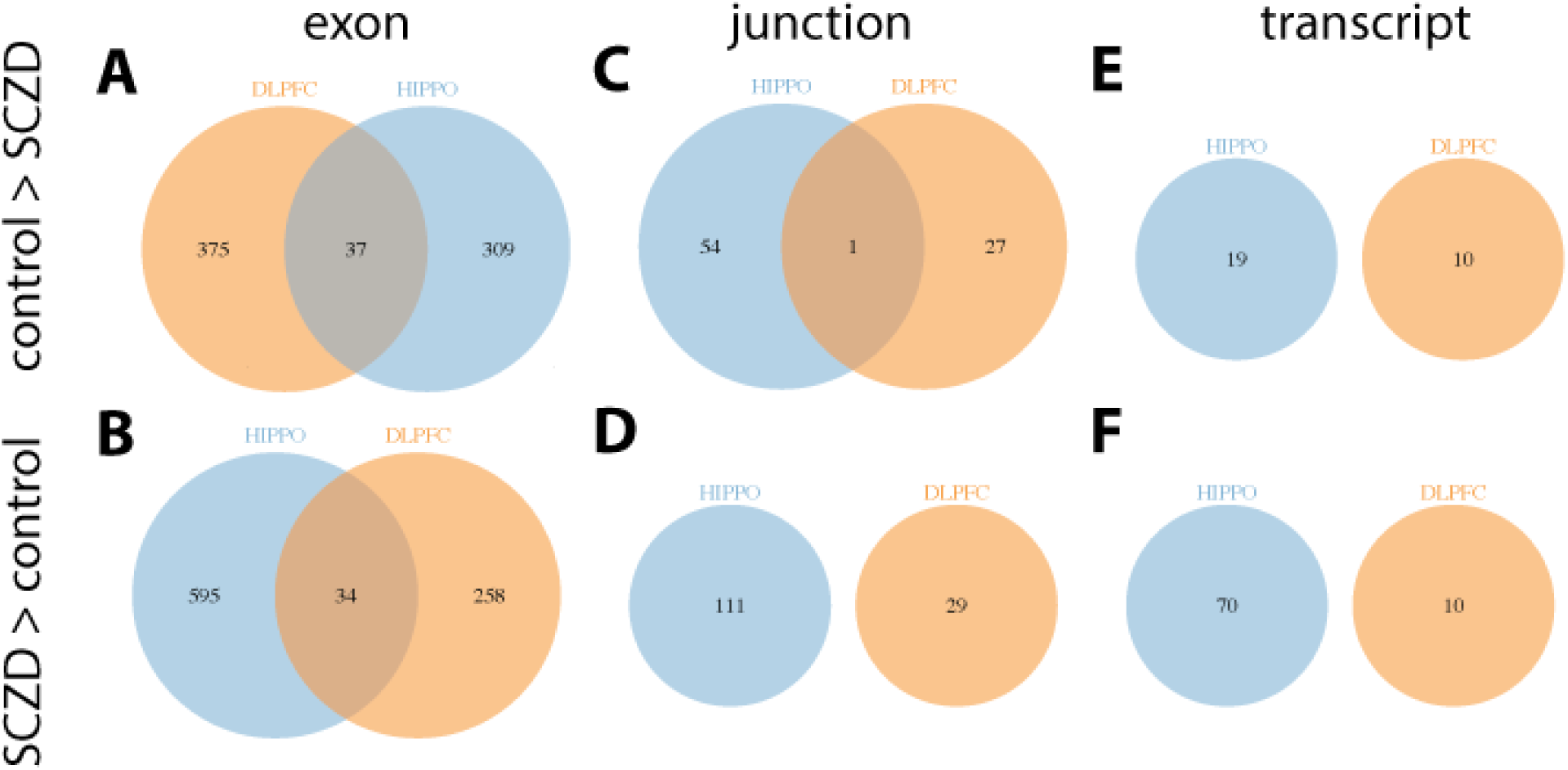
Feature DE by SCZD status. Venn diagrams of differentially expressed features between non-psychiatric controls and SCZD affected individuals by brain region (DLPFC: FDR <10%, HIPPO: FDR <20%). Exons (**A** and **B**), exon-exon junctions (**C** and **D**) and transcripts (**E** and **F**) with higher expression in non-psychiatric controls (**A**, **C** and **E**) or in SCZD cases (**B**, **D** and **F**).

**Figure S21.**
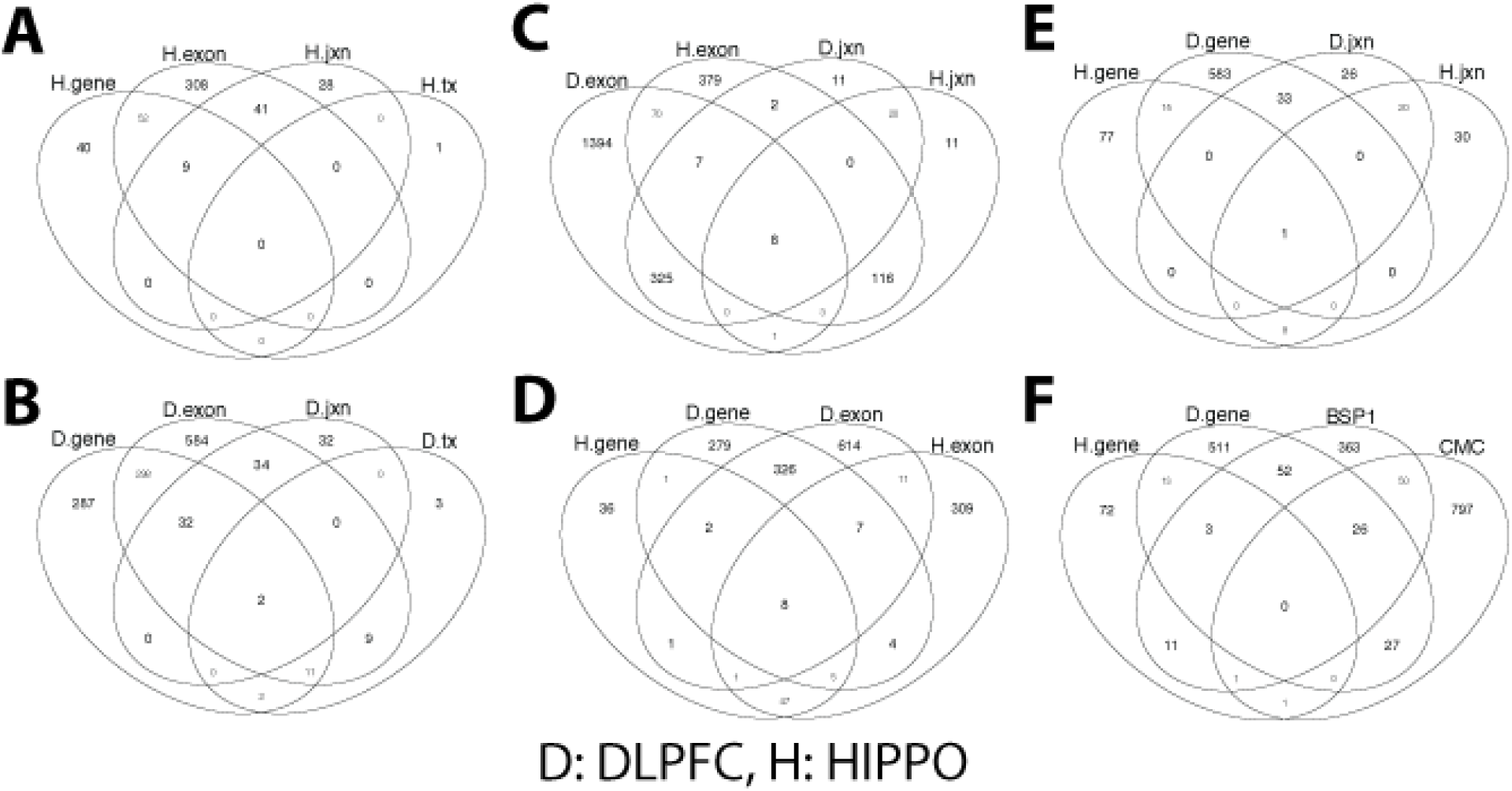
Model and feature DE by SCZD status. Venn diagrams of differentially expressed features (FDR<10%) between non-psychiatric controls and SCZD affected individuals by brain region and studies grouped by gene id. Differentially expressed features in HIPPO (**A**) or DLPFC (**B**). Exons and exon-exon junctions differentially expressed by SCZD status (FDR<10%) grouped by gene ID (**C**), gene and exons (**D**), as well as genes and exon-exon junctions (**E**) across brain regions. (**F**) Differentially expressed genes in BrainSeq Phases I and II as well as in the CommonMind Consortium (CMC).

**Figure S22.**
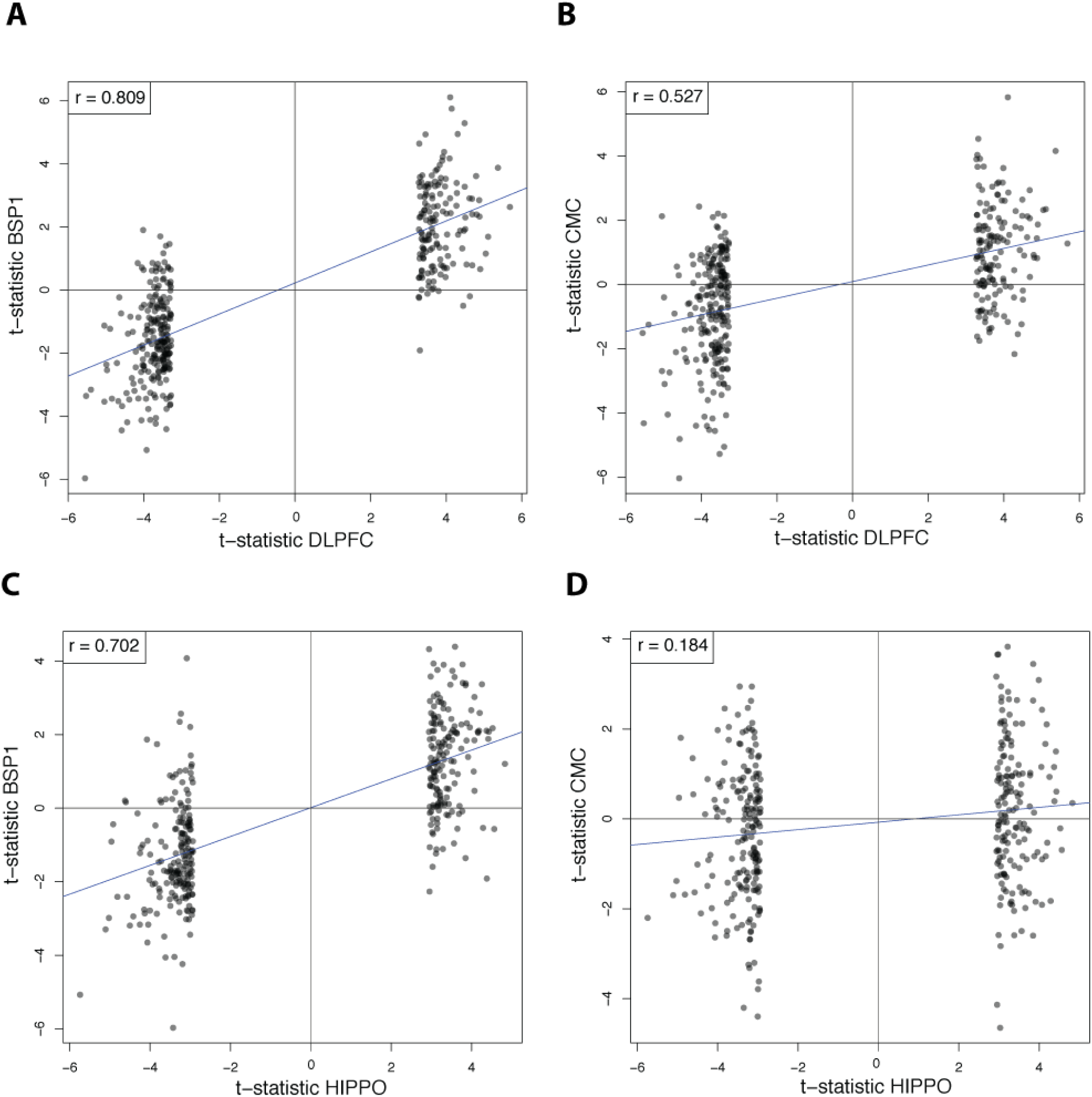
T-statistics across regions and datasets for top SCZD results. T-statistics for the top 400 differentially expressed genes between non-psychiatric controls and SCZD cases for either DLPFC (**A** and **B**) or HIPPO (**C** and **D**) compared against the t-statistics in BrainSeq Phase 1 DLPFC (**A** and **C**) and the CommonMind Consortium dataset (**B** and **D**). Correlation and the linear regression trend (blue line) are shown for each plot.

**Figure S23.**
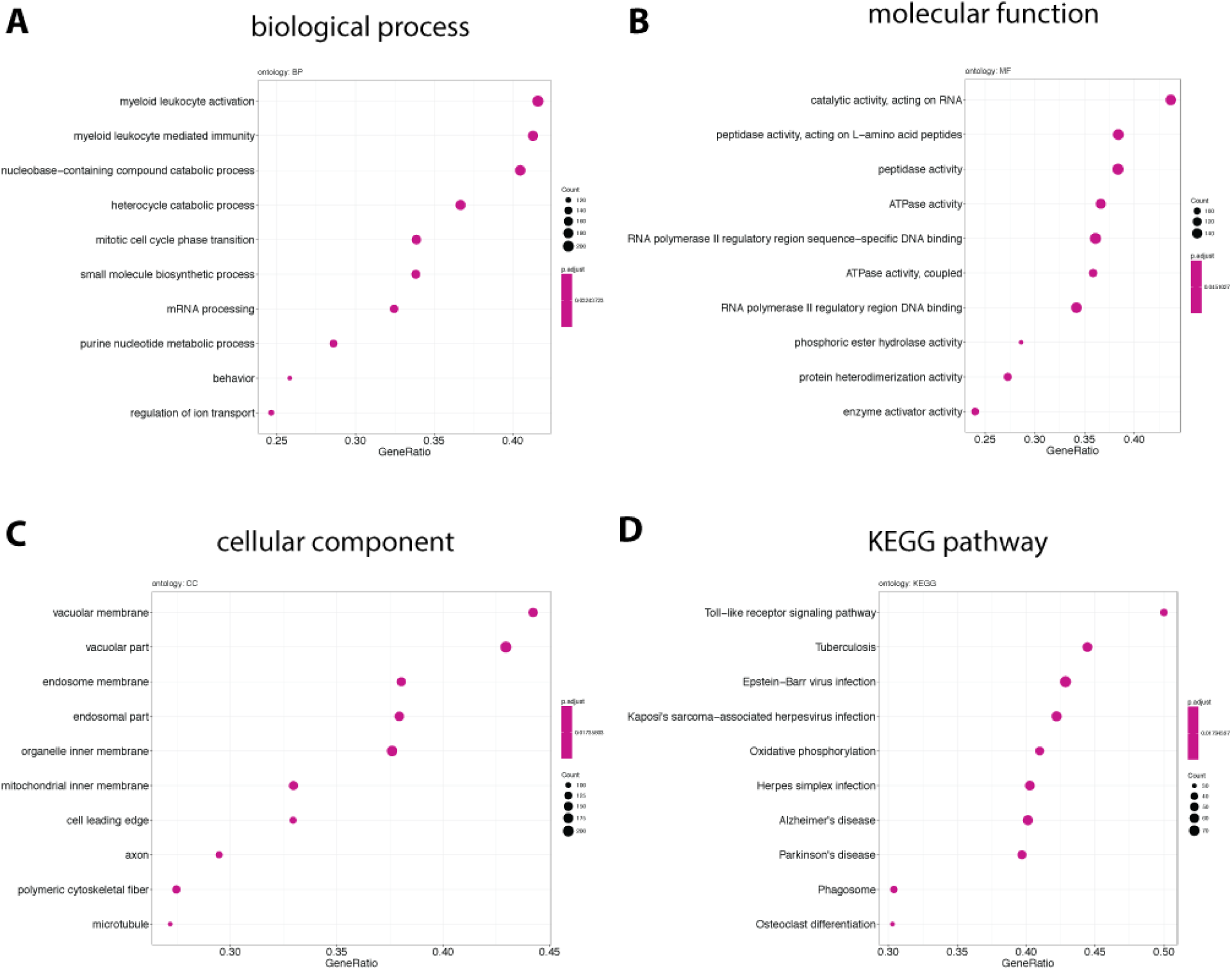
GSE DLPFC results. Gene set enrichment (GSE) results for differentially expressed features between SCZD cases and non-psychiatric controls in DLPFC. Enriched (**A**) biological processes, (**B**) molecular function, (**C**) cellular component ontologies and (**D**) Kyoto Encyclopedia of Genes and Genomes (KEGG) pathways. Biological process is duplicated here from **Figure 3** to facilitate comparisons.

**Figure S24.**
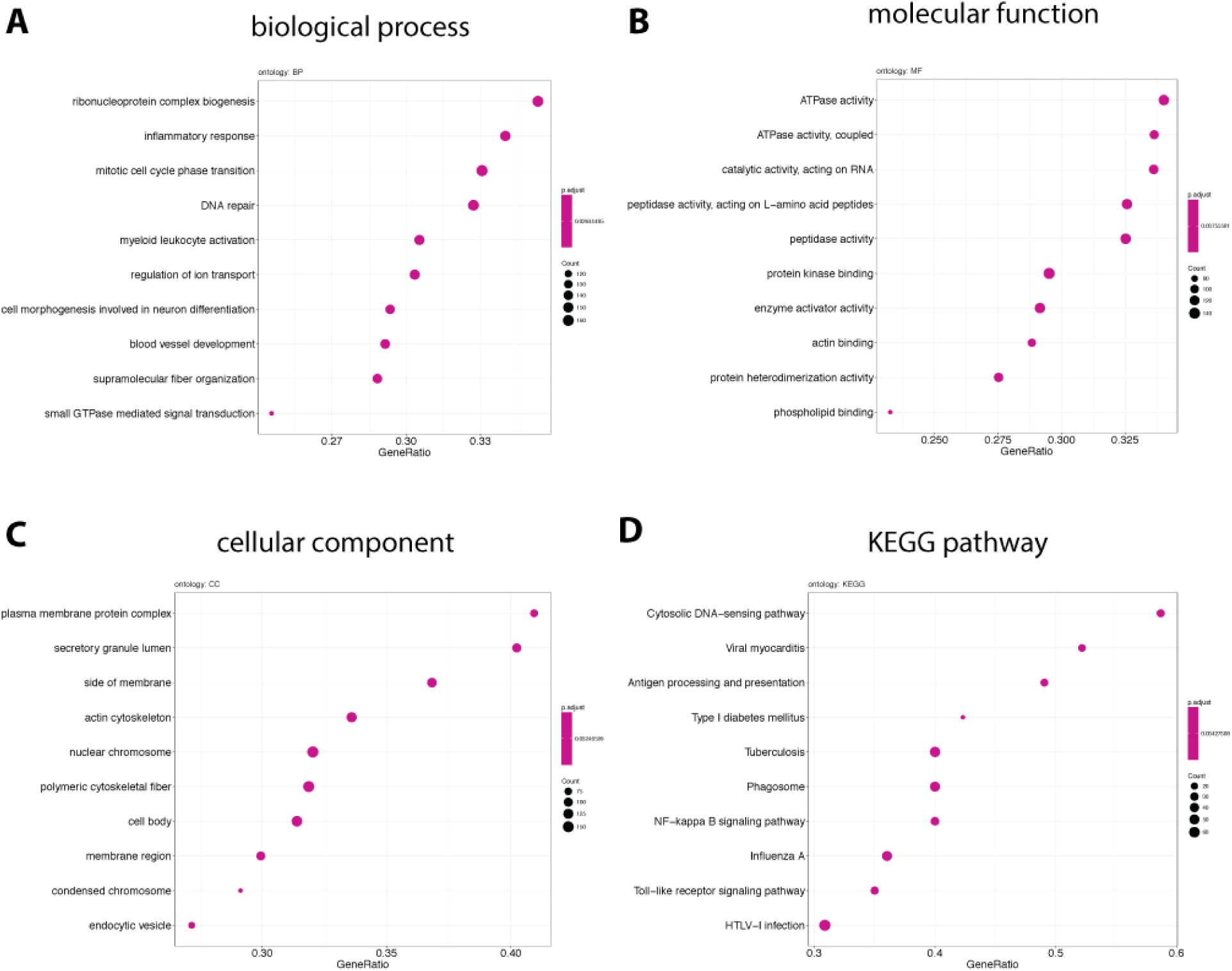
GSE HIPPO results. Gene set enrichment (GSE) results for differentially expressed features between SCZD cases and non-psychiatric controls in HIPPO. Enriched (**A**) biological processes, (**B**) molecular function, (**C**) cellular component ontologies and (**D**) Kyoto Encyclopedia of Genes and Genomes (KEGG) pathways. Biological process is duplicated here from **Figure 3** to facilitate comparisons.

**Figure S25.**
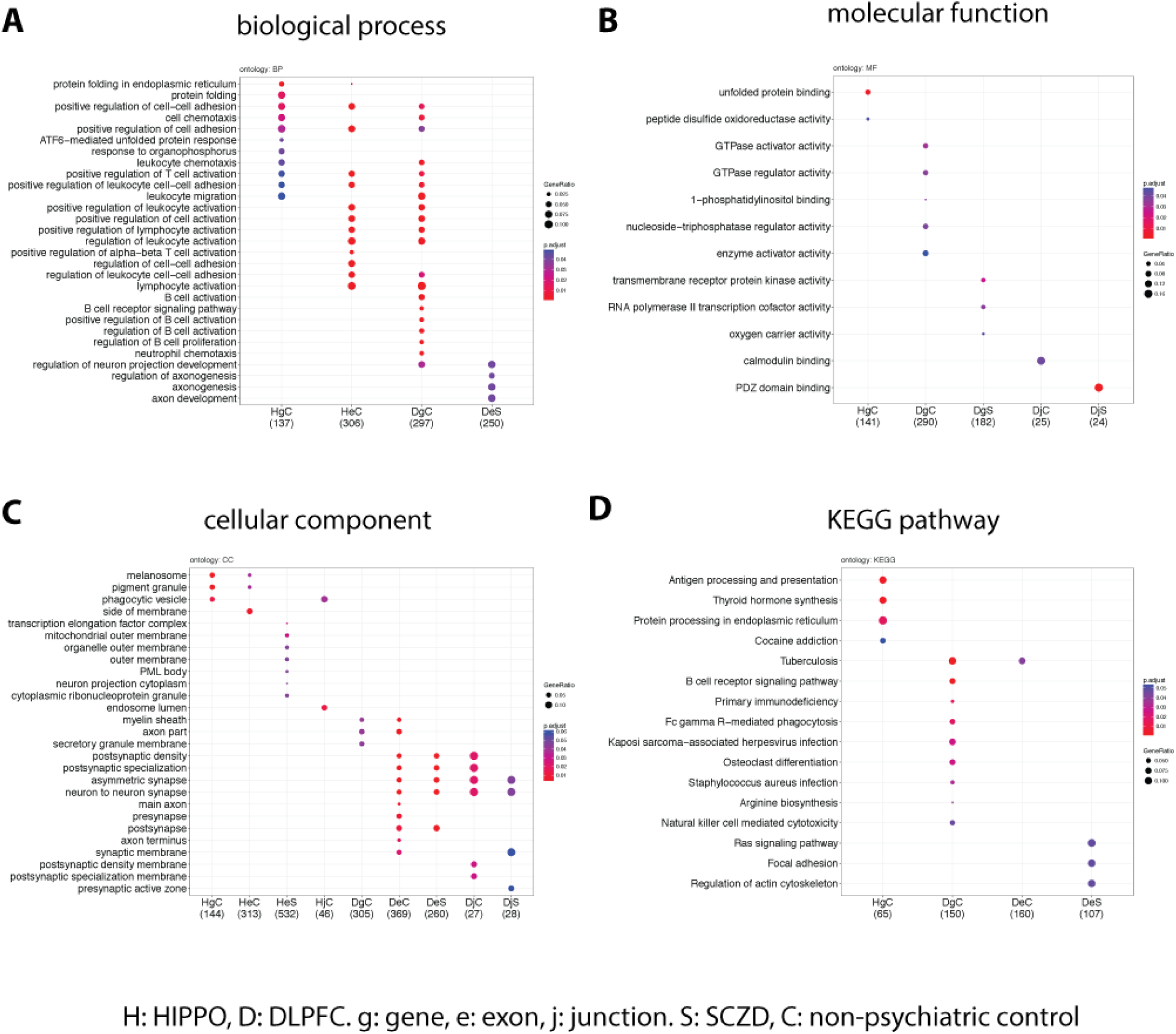
SCZD enriched GOs. Gene ontology enrichment results for differentially expressed features between SCZD cases and non-psychiatric controls either in the DLPFC or HIPPO brain region. Enriched (**A**) biological processes, (**B**) molecular function, (**C**) cellular component ontologies and (**D**) Kyoto Encyclopedia of Genes and Genomes (KEGG) pathways at the gene, exon or junction feature levels separated by higher expression in SCZD cases or non-psychiatric controls.

**Figure S26.**
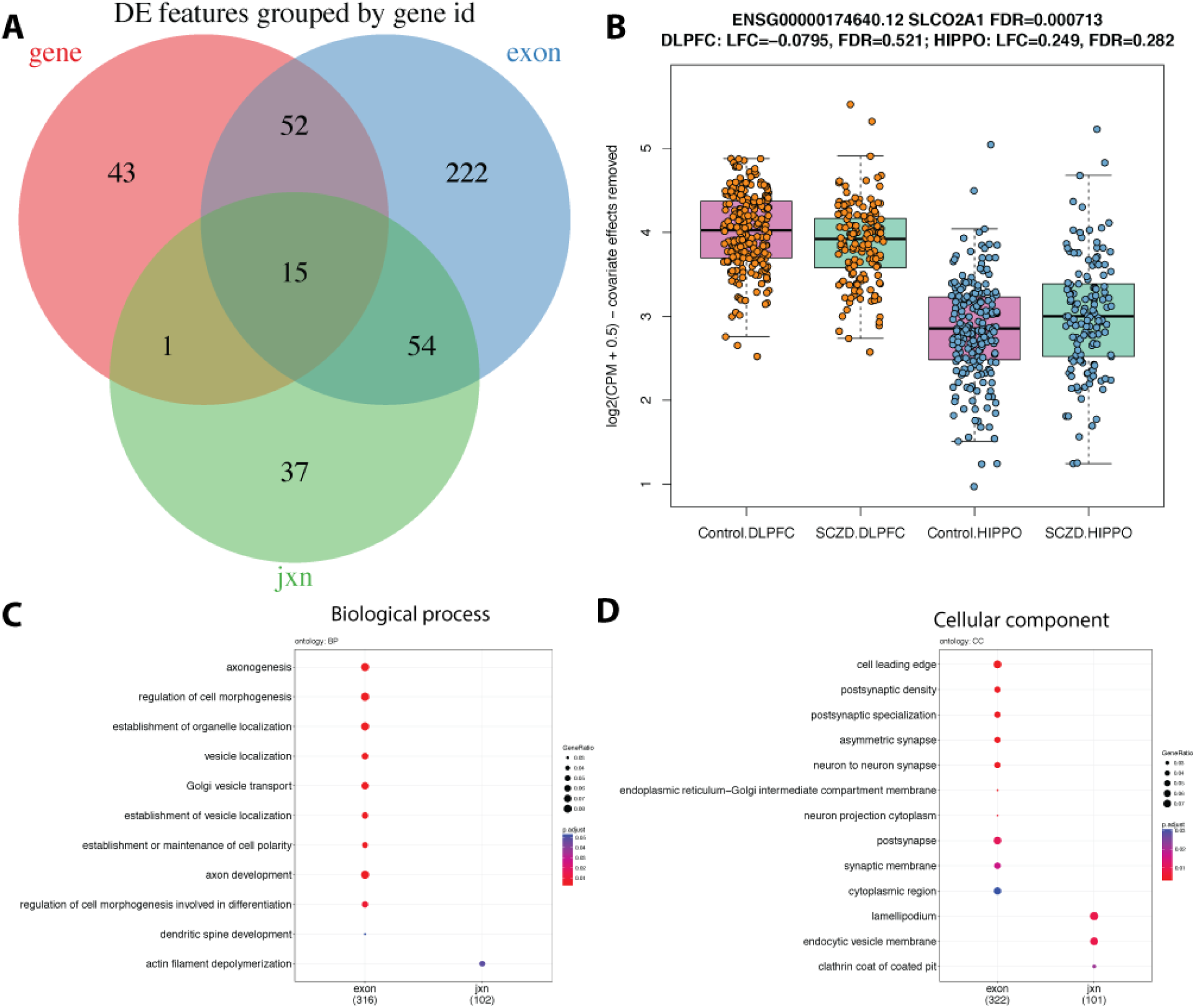
Differential expression by interaction between SCZD diagnosis status and brain region. (**A**) Differentially expressed features grouped by gene id excluding 29 un-annotated DE exon-exon junctions. (**B**) The *SCL02A1* gene has a significantly different relationship between expression and SCZD status across DLPFC (decreased in SCZD versus controls) and HIPPO (increased in SCZD versus controls); LFC: log2 fold change. Enriched biological processes (**C**) and cellular components (**D**) among the genes with DE signal at the exon and exon-exon junction levels. No terms were enriched among the 111 DEGs with gene support.

**Figure S27.**
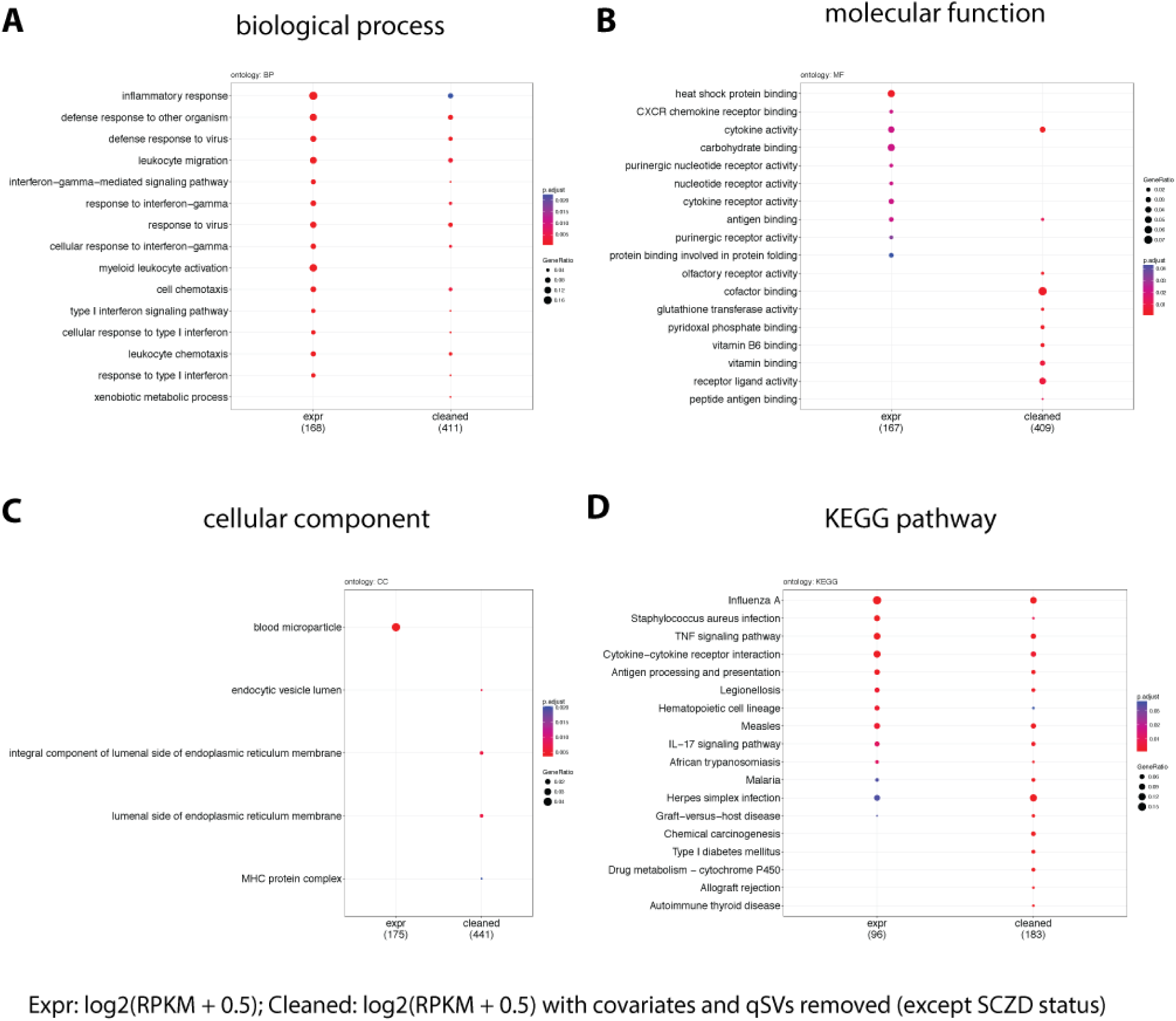
Enriched GOs among highly correlated genes in DLPFC and HIPPO adults. Gene ontology enrichment results for genes with significant correlation between HIPPO and DLPFC (FWER<5%) among the matched individuals from the SCZD case-control analysis (n=265 individuals). Enriched (**A**) biological processes, (**B**) molecular function, (**C**) cellular component ontologies and (**D**) Kyoto Encyclopedia of Genes and Genomes (KEGG) pathways at the gene level either with the raw expression values [log2(RPKM + 0.5)] or the cleaned expression values [log2(RPKM + 0.5) with covariates and qSVs removed while retaining the SCZD diagnosis effect].

**Figure S28.**
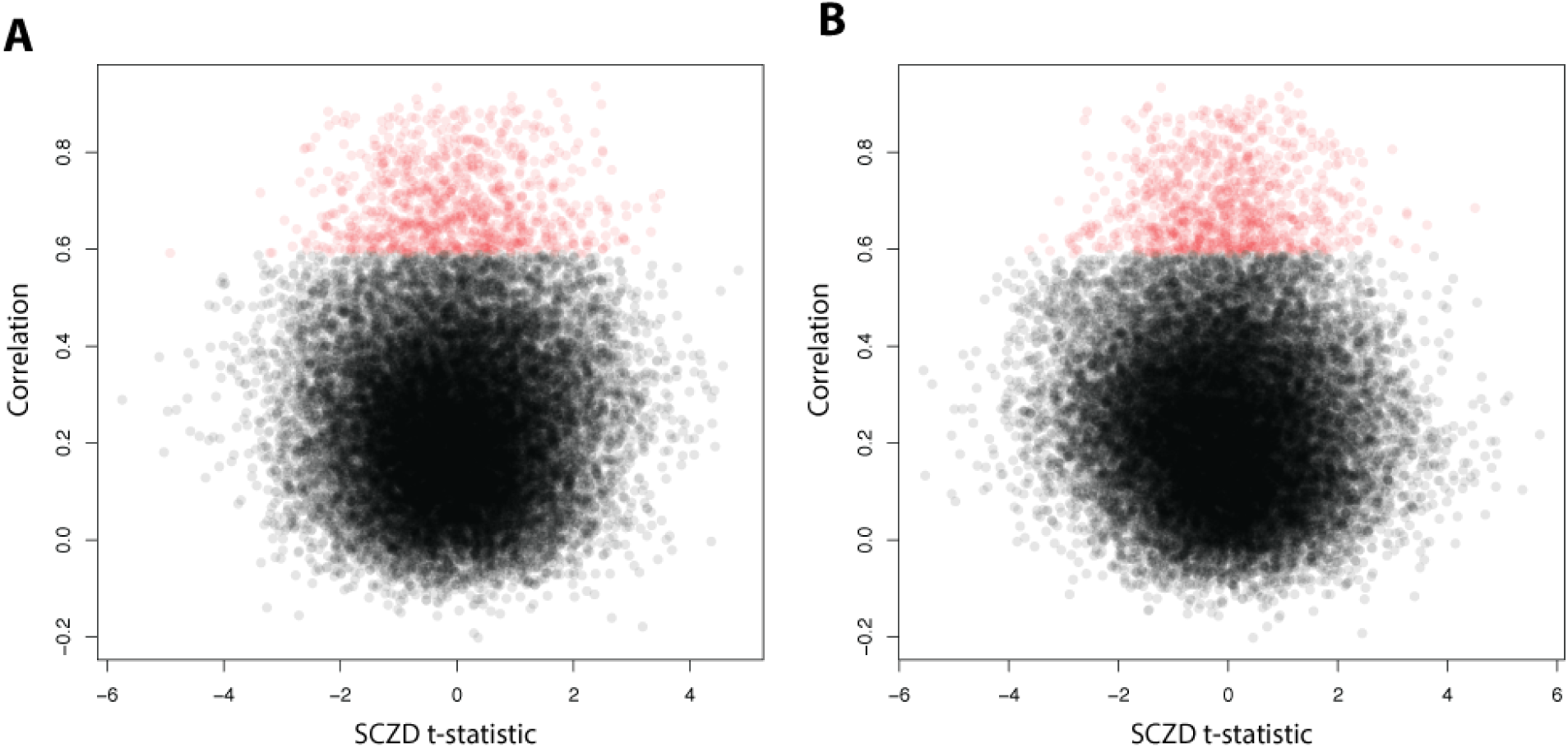
Correlated genes in DLPFC and HIPPO adults compared vs SCZD case-control t-statistic. Scatter plot of the correlation between HIPPO and DLPFC compared against the SCZD t-statistic with significantly correlated genes (FWER<5%) labeled in red. (**A**) HIPPO and (**B**) DLPFC SCZD t-statistics. The correlation was calculated with the cleaned expression values: log2(RPKM + 0.5) with covariates and qSVs removed while retaining the SCZD diagnosis effect.

**Figure S29.**
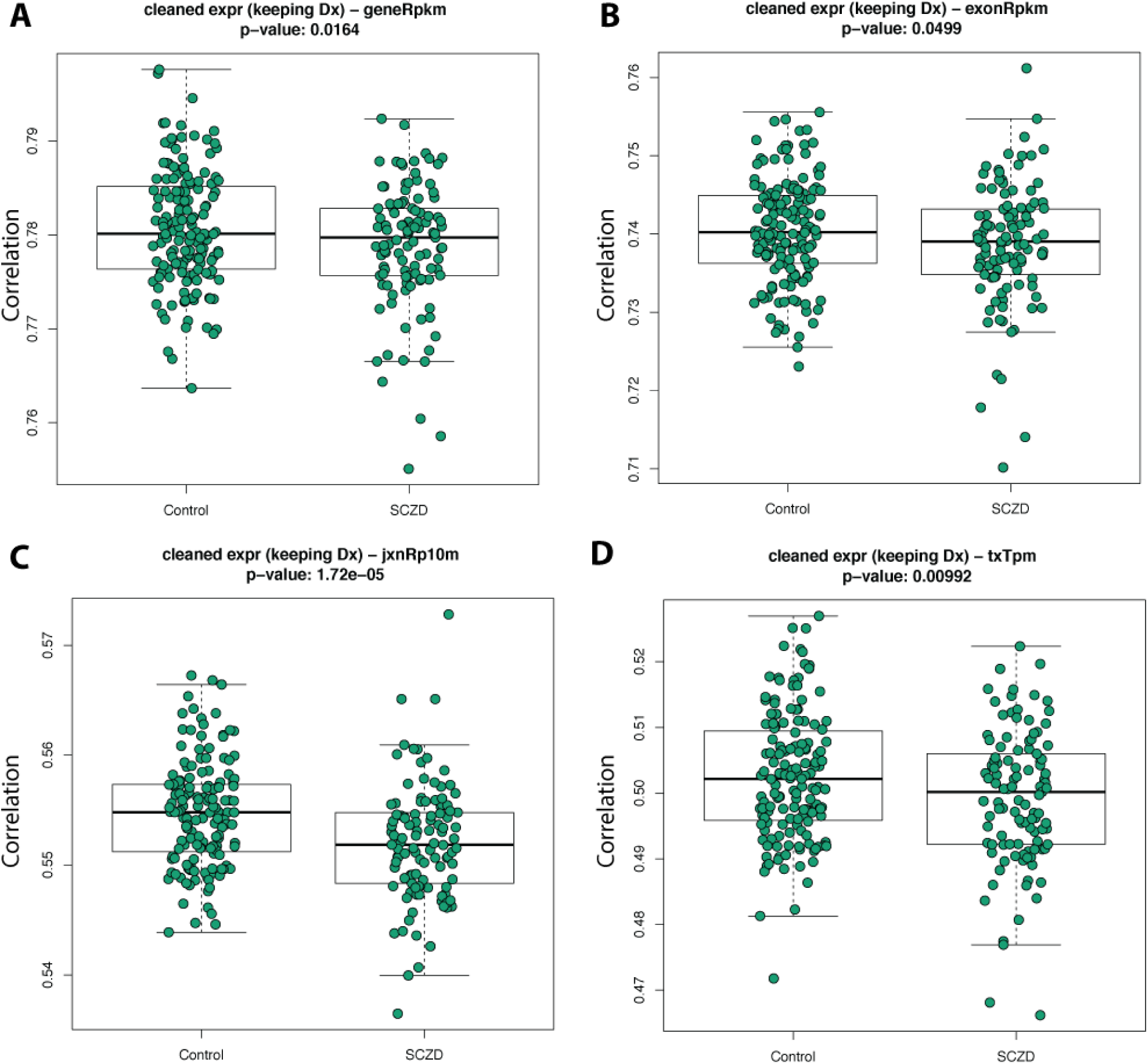
Correlation differences in HIPPO and DLPFC by SCZD diagnosis for all feature levels. Correlation for each expression feature across HIPPO and DLPFC per individual (n=265) shows that SCZD affected individuals have decreased correlation at the (**A**) gene, (**B**) exon, (**C**) exon-exon junction and (**D**) transcript levels compared to non-psychiatric controls. The correlation was calculated with the cleaned expression values: log2(RPKM + 0.5) for genes and exons, log2(RP10M + 0.5) for exon-exon junctions and log2(TPM + 0.5) for transcripts with covariates and qSVs removed while retaining the SCZD diagnosis effect.

**Figure S30.**
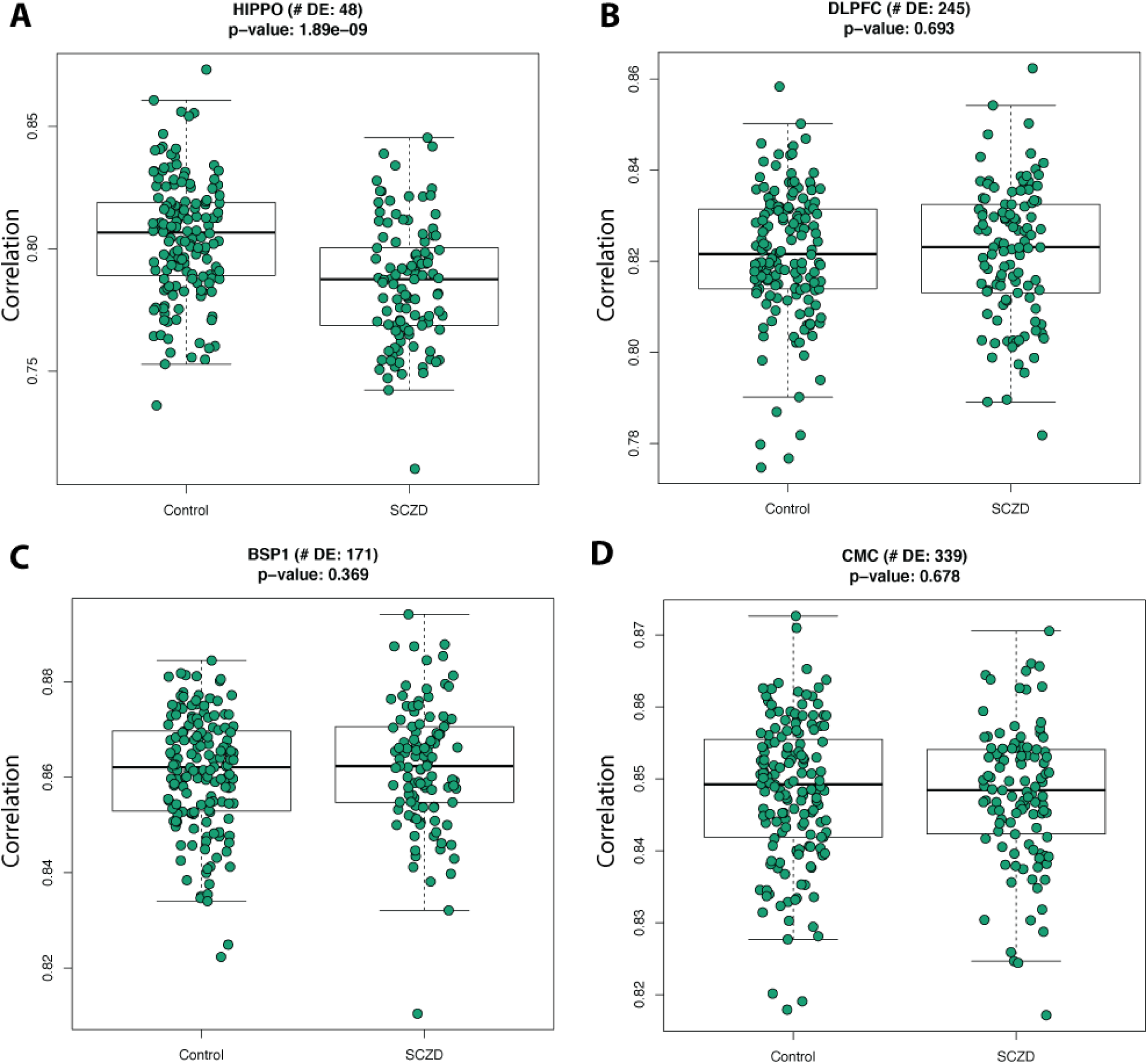
Correlation differences in HIPPO and DLPFC by SCZD diagnosis for SCZD differentially expressed genes. For genes differentially expressed by SCZD status, their correlation across HIPPO and DLPFC per individual (n=265) shows that SCZD have decreased correlation for the (**A**) HIPPO SCZD DEGs, but not for the (**B**) DLPFC, (**C**) BrainSeq Phase 1 (BSP1), or (**D**) CommonMind Consortium (CMC) DEGs.

**Figure S31.**
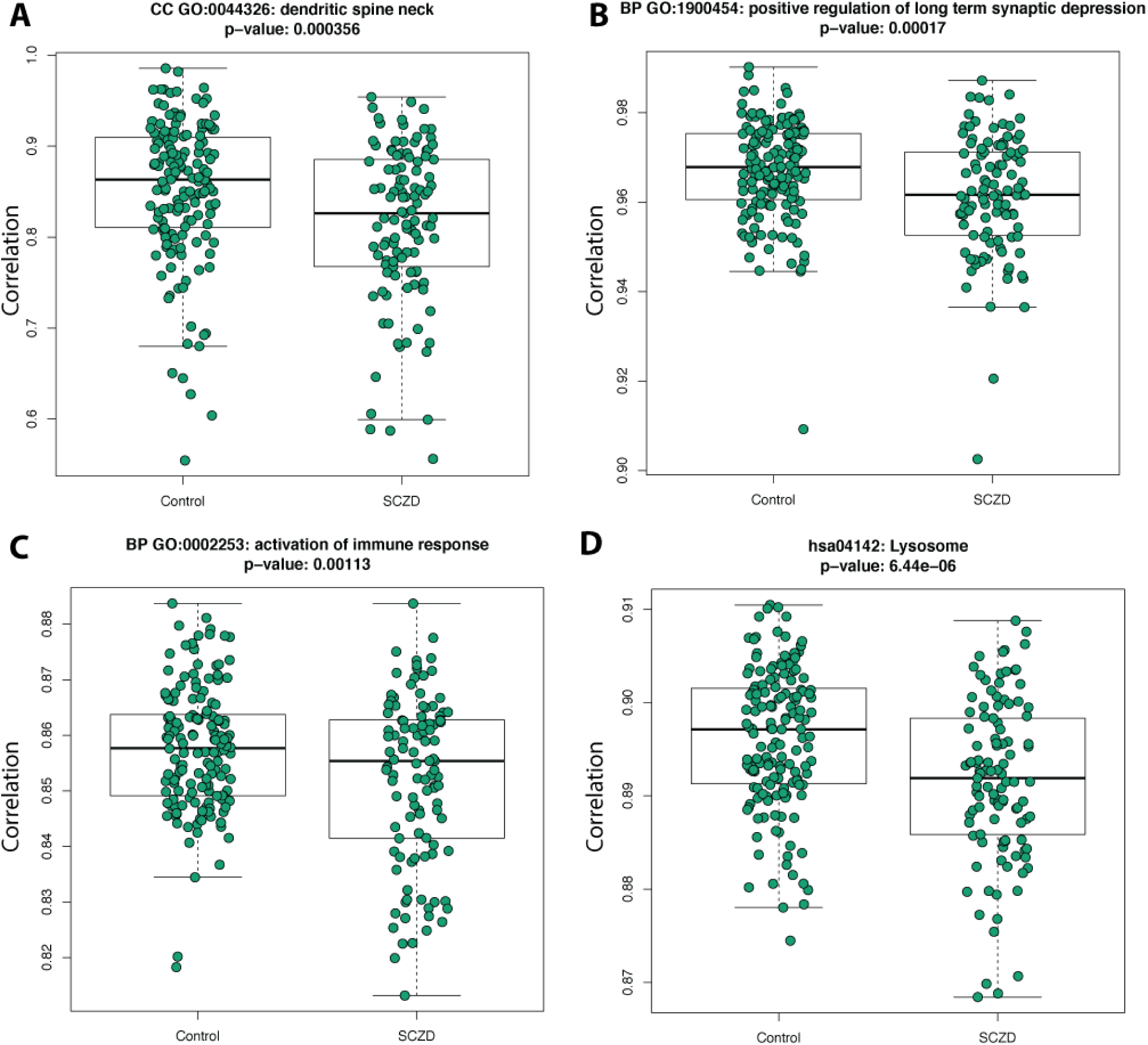
Highlighted GO and KEGG pathways with significant correlation differences in HIPPO and DLPFC by SCZD diagnosis. Sets of genes grouped by their membership to GO terms (level 3) and KEGG pathways that show differential correlation among individuals with SCZD and non-psychiatric controls. Highlighted significantly (FDR<5%) differentially correlated GO terms (**A**: 0044326, **B**: 1900454, **C**: 0002253) and KEGG pathway (**D**: hsa04142) among HIPPO and DLPFC. The correlation was calculated with the cleaned gene expression values: log2(RPKM + 0.5) with covariates and qSVs removed while retaining the SCZD diagnosis effect.

**Figure S32.**
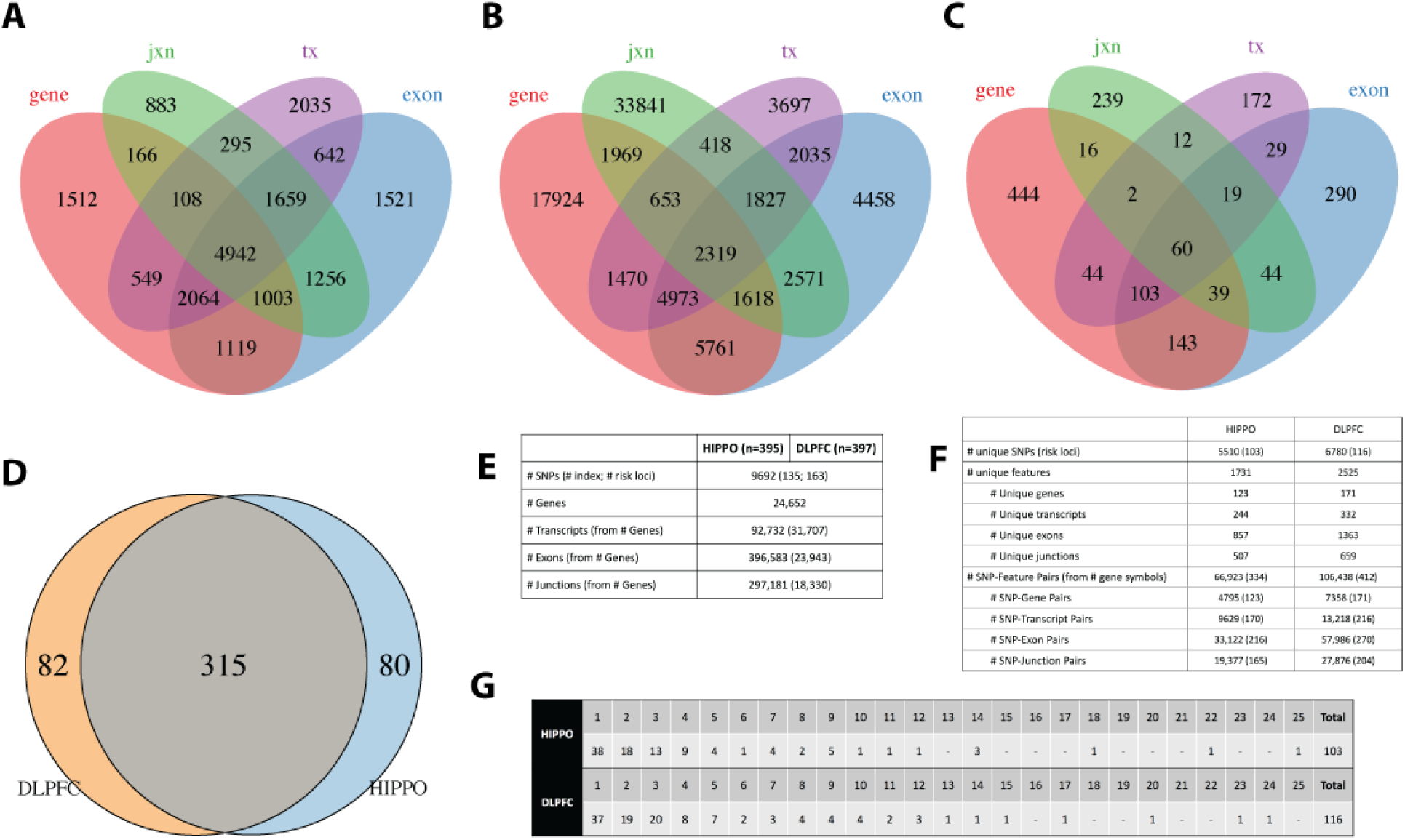
eQTLs by feature and risk analysis. (**A**) HIPPO eQTLs grouped by gene ID. (**B**) region-dependent eQTLs grouped by SNP ID. (**C**) region-dependent eQTLs group by gene ID. (**D**) Brain samples by individual used for the SNP GWAS risk analysis. (**E**) Input information for the SNP GWAS risk analysis. (**F**) SNP GWAS risk analysis results summary by brain region. (**G**) Number of risk GWAS loci that are eQTLs in BrainSeq Phase II that pair to genes by brain region.

**Figure S33.**
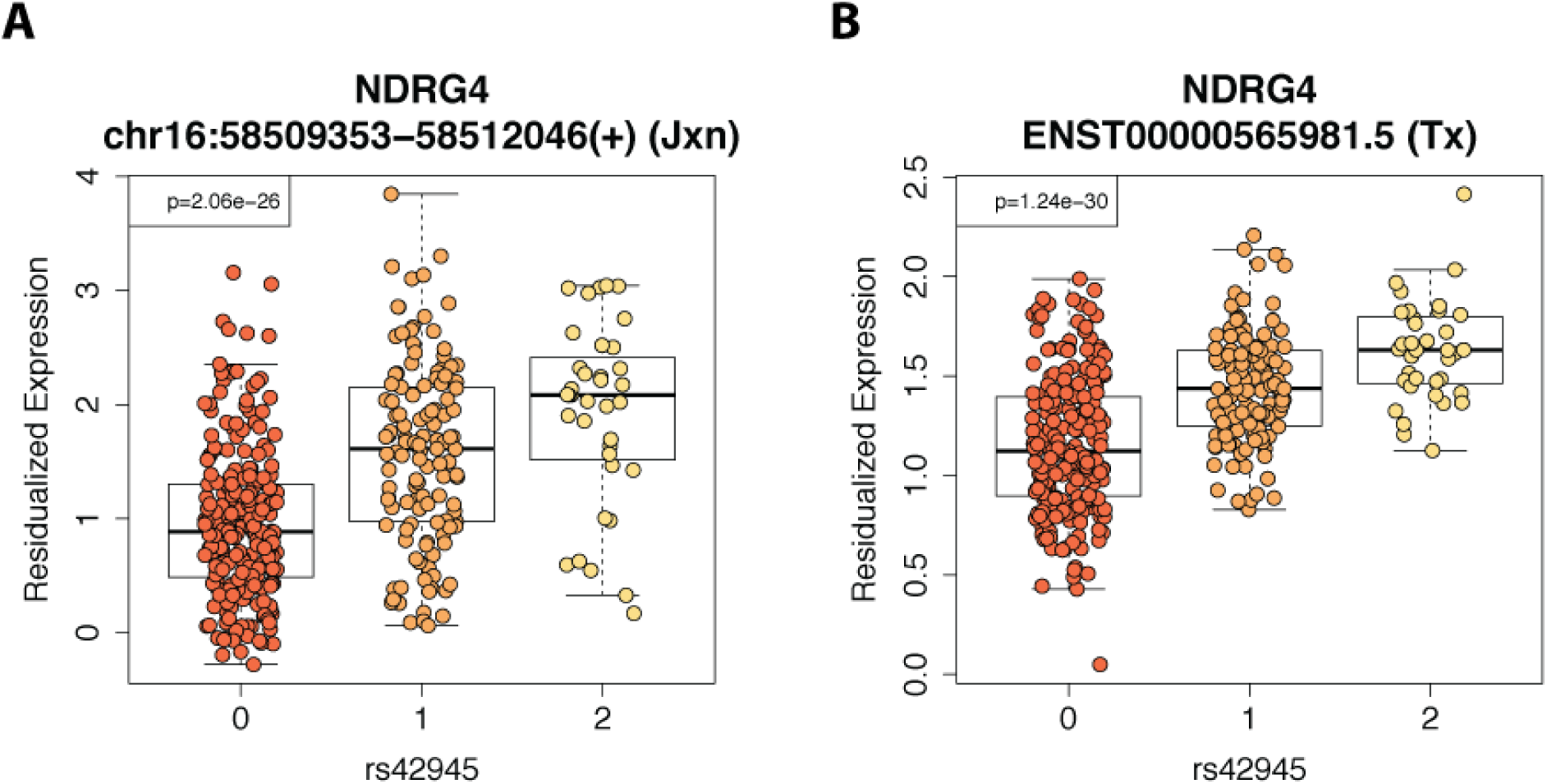
Example GWAS PGC2 index risk SNP eQTLs in HIPPO. An exon-exon junction (**A**) and a transcript (**B**) from the gene *NDRG4* are in eQTL associations with the schizophrenia risk snp rs42945 in HIPPO (and DLPFC: not shown). Residual expression is log2(RP10M + 1) or log2(TPM + 1) with SCZD diagnosis, sex, five ancestry PCs and expression PCs removed.

**Figure S34.**
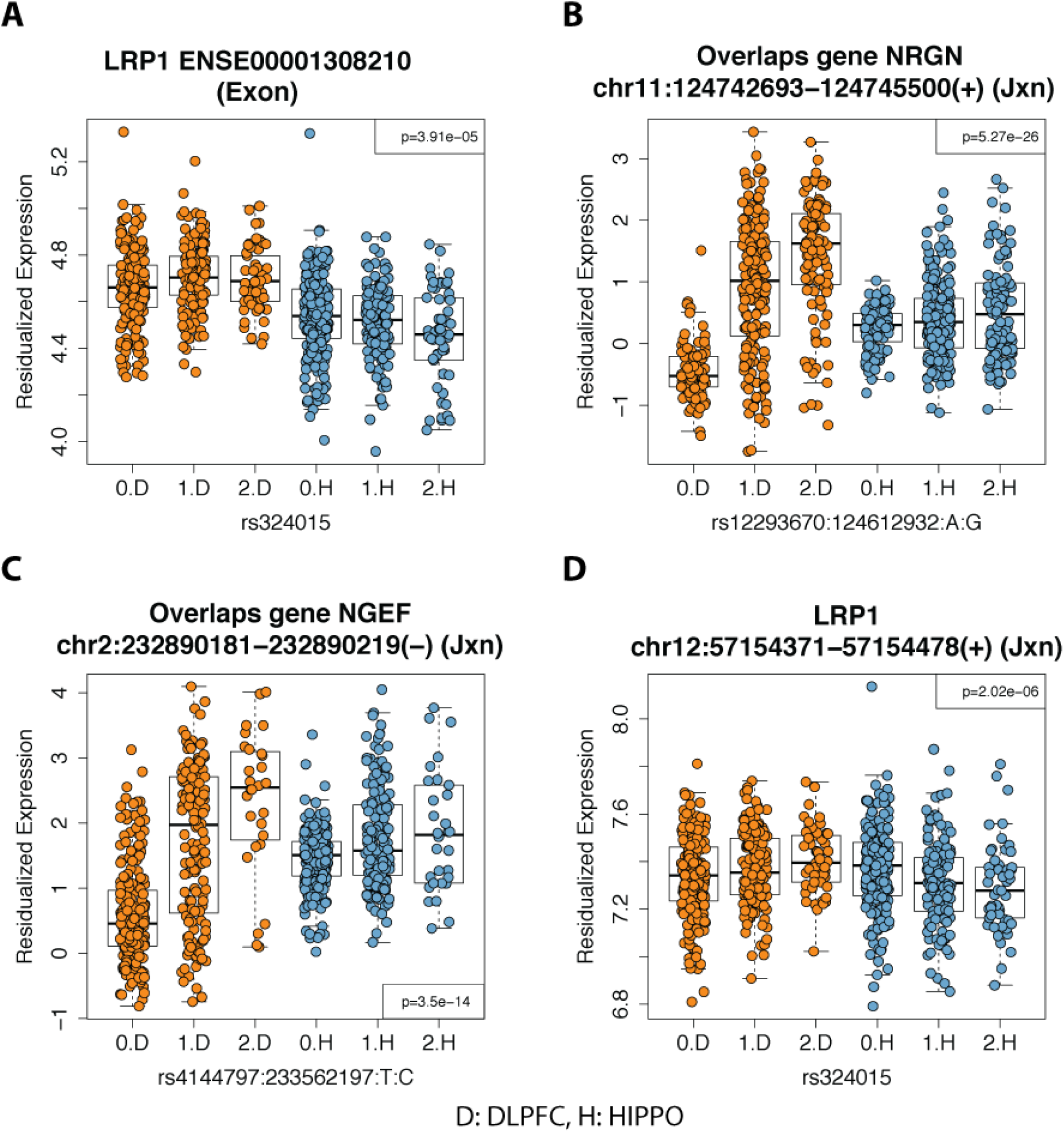
Brain-region dependent eQTLs that involve GWAS PCG2 index risk SNPs. GWAS PCG2 index risk snps rs324015 (**A** and **D**), rs12293670 (**B**) and rs4144797 (**C**) have brain-region dependent eQTL associations with DLPFC and HIPPO. They involve either an exon (**A**) or exon-exon junctions (**B-D**) for genes *LRP1*, *NRGN* and *NGEF.* Two of the exon-exon junctions are un-annotated (**B**, **C**) though they are in the same strand as the only gene they overlap: *NRGN* and *NGEF*, respectively. Residual expression is log2(RPKM + 1) or log2(RP10M + 1) with SCZD diagnosis, sex, five ancestry PCs and expression PCs removed while retaining the brain region effect. Orange: DLPFC, blue: HIPPO.

## Supplementary Tables

**Table S1**. Demographic and sequencing metrics by group for the RNA-seq samples. See file SupplementaryTablelxIsx.

**Table S2**. Differentially expressed features for each of the three main models: (1) HIPPO versus DLPFC developmental differences among non-psychiatric controls, (2) HIPPO versus DLPFC in both adults and prenatal age groups among non-psychiatric controls, (3) SCZD cases versus non-psychiatric controls for both DLPFC and HIPPO. See file SupplementaryTable2.tar.gz.

**Table S3**. Differentially (FDR<5%) correlated GO terms and KEGG pathways at the individual level between SCZD affected individuals and non-psychiatric controls. See file SupplementaryTable3.xlsx.

**Table S4**. Genomic coordinates of the top 1000 degradation-associated expressed regions across DLPFC and HIPPO. See file SupplementaryTable4.xlsx.

